# Genomic adaptations to aquatic and aerial life in mayflies and the origin of wings in insects

**DOI:** 10.1101/2019.12.29.888636

**Authors:** Isabel Almudi, Joel Vizueta, Alex de Mendoza, Chris Wyatt, Ferdinand Marletaz, Panos Firbas, Roberto Feuda, Giulio Masiero, Patricia Medina, Ana Alcaina, Fernando Cruz, Jessica Gómez-Garrido, Marta Gut, Tyler S. Alioto, Carlos Vargas-Chavez, Kristofer Davie, Bernhard Misof, Josefa González, Stein Aerts, Ryan Lister, Jordi Paps, Julio Rozas, Alejandro Sánchez-Gracia, Manuel Irimia, Ignacio Maeso, Fernando Casares

## Abstract

The first winged insects underwent profound morphological and functional transformations leading to the most successful animal radiations in the history of earth. Despite this, we still have a very incomplete picture of the changes in their genomes that underlay this radiation. Mayflies (Ephemeroptera) are one of the extant sister groups of all other winged insects and therefore are at a key phylogenetic position to understand this radiation. Here, we describe the genome of the cosmopolitan mayfly *Cloeon dipterum* and study its expression along development and in specific organs. We discover an expansion of odorant-binding proteins, some expressed specifically in the breathing gills of aquatic nymphs, suggesting a novel sensory role for gills. In contrast, as flying adults, mayflies make use of an enlarged set of opsins and utilise these visual genes in a sexually dimorphic manner, with some opsins expressed only in males. Finally, to illuminate the origin of wings, we identify a core set of deeply conserved wing-specific genes at the root of the pterygote insects. Globally, this is the first comprehensive study of the structure and expression of the genome of a paleopteran insect and shows how its genome has kept a record of its functional adaptations.

## Introduction

The first insects colonized the land more than 400 Myr ^1^. But it was only after insects evolved wings that this lineage (the Pterygotes) became the most prominent animal group in terms of number and diversity of species, and completely revolutionized Earth ecosystems. The development of wings also marked the appearance of hemimetabolous development. Concomitant with the development of wings, pterygotes also developed an hemimetabolous life cycle ^2^, with two clearly differentiated living phases (non-flying juveniles and flying adults). This allowed them to specialize functionally and exploit two entirely different ecological niches. This is still the life cycle of Ephemeroptera (mayflies) and Odonata (damselflies and dragonflies), the extant Paleoptera (“old wing”) orders, the sister group of all other winged insects (Figure 1a), and which undergo a radical ecological switch where aquatic nymphs metamorphose into terrestrial flying adults. The appearance of wings and the capacity to fly greatly increased the capability of insects for dispersal, escape and courtship and allowed them access to previously unobtainable nutrient sources, while establishing new ecological interactions. This ‘new aerial dimension in which to experience life’^3^ created new evolutionary forces and constraints that since, have been continuously reshaping the physiology, metabolism, morphology and sensory capabilities of different pterygote lineages-evolutionary changes that should ultimately be traced back to genomic features associated to these adaptations. However, our knowledge of these genomic features is still very incomplete, mostly because paleopteran lineages (Figure 1a), have not been extensively studied from a genomic and transcriptomic perspective.

**Figure 1.**
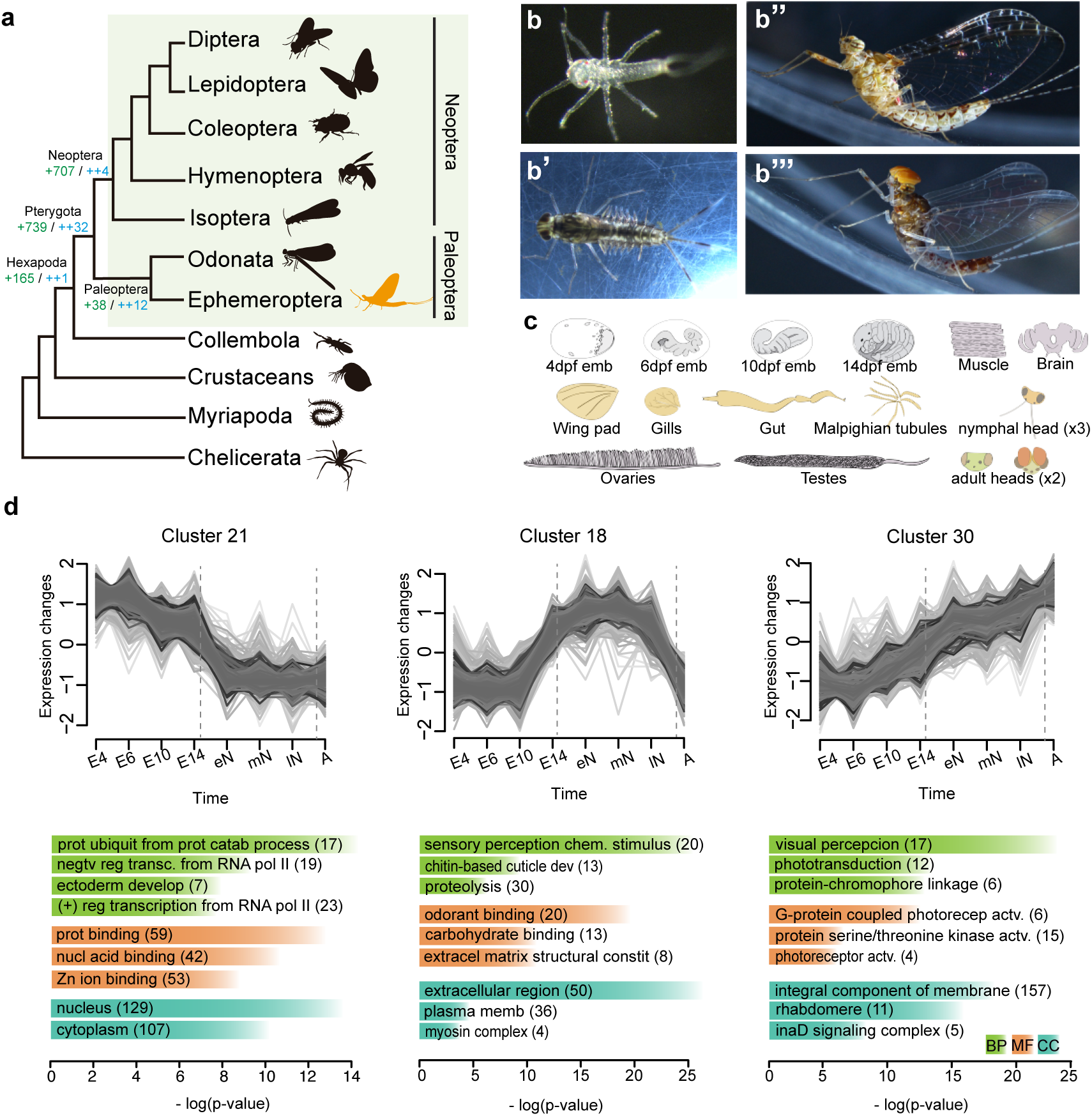
Genome and transcriptomes of the mayfly *C. dipterum*. **a**, Simplified arthropod phylogeny highlighting Pterygota insects (green rectangle). Ephemeroptera (mayflies, in orange) are within Paleoptera, the sister group to all other winged insects. Number of gene gains (in green) and “core” gene gains (in blue) is shown at the root of selected lineages. **b**, *C. dipterum* individuals. **b**, Freshly hatchling. **b’**, Late nymph. **b’’**, Female adult. **b’’’**, Male adult. **c**, Cartoons depicting organs and developmental stages used for the transcriptome profiling. In grey, embryonic stages used: 4 days post fertilization (dpf) or germ disc stage, 6 dpf or segmented embryo stage, 10 dpf or “revolution” stage and 14 dpf or pre-nymph stage. In orange, nymphal tissues: wing pads, gills, heads, gut and Malpighian tubes. In pale pink, adult organs: muscle, brain, female heads, male heads, ovaries and testis. **d**, Three M-Fuzz clusters showing high expression during embryonic, nymphal and adult stages (E.4: 4 days post fertilization (dpf) embryo, E.6: 6 dpf embryo, E.10: 10 dpf embryo, E.14 14 dpf embryo, eN: early nymph head, mN: mid nymph head, lN: late nymph head, A: adult head). Each of them contains genes that are enrichment in GO terms related to developmental processes, chitin and chemical stimuli perception and visual perception, respectively (BP: Biological Process, CC: Cellular component, MF: Morphological Function).

Mayflies are an ideal group to fill this gap of knowledge. By living in both terrestrial and aquatic environments, mayflies had to develop different sensory, morphological and physiological adaptations to each of these ecological niches. For example, mayflies have abdominal gills during the aquatic stages, a feature that places them in a privileged position to assess the different hypotheses accounting for the origin of wings ^4-9^. Moreover, some mayfly families exhibit a striking sexual dimorphism in their visual systems, which in the case of the Baetidae family includes the presence of a second set of very large compound eyes in males (Figure 1b). All these features make mayflies a fundamental order to investigate the origin of evolutionary novelties associated with the conquest of new habitats. The recent establishment of a continuous culture system of the mayfly *C. dipterum*, a cosmopolitan Baetidae species with a life cycle of just six weeks, makes it now possible the access to all embryonic and postembryonic stages ^10^, overcoming past challenges to study paleopterans which are generally not very amenable to rear in the lab.

Here we sequenced the genome and profiled a comprehensive set of stage- and organ-specific transcriptomes of *Cloeon dipterum*. Our analyses identify potential genomic signatures associated to the mayfly adaptations to the aquatic lifestyle, to innovations in its visual system and novel genetic players in the evolution of wings. The results from this work establish *C. dipterum* as a new platform to investigate insect genomics, development and evolution from a phylogenetic vantage point.

## Results

### *C. dipterum* genome assembly

We sequenced and assembled the genome of an inbred line of the mayfly species *C. dipterum* using both Illumina and Nanopore technologies (see Methods, Supplementary Figure 1, Supplementary Table 1-3). The assembled genome of *C. dipterum* is 180 Mb, which in comparison to other pterygote species ^11-14^, constitutes a relatively compact genome, probably due to the low fraction of Transposable Elements (TEs) (5%, in contrast to the median of 24% ± 12% found in other insects ^15^, Supplementary Figure 1). *C. dipterum* genome was assembled in 1395 scaffolds, with 0.461 Mb of N50. Protein coding gene annotation resulted in 16730 genes, which represented 96.77% and 98.2% of gene completeness using CEGMA v2.5 ^16^ and BUSCO v3 ^17^ databases, respectively. These annotated protein-coding genes were used to reconstruct orthologous gene families of *C. dipterum* and other animal species (Supplementary Table 4, Supplementary Table 5) allowing the study of gene gains and losses at the origin of winged insects (Figure 1a, Supplementary Figure 1). Along with the genomic DNA, we transcriptionally profiled a large number of time points along *C. dipterum’s* life cycle and a set of nymphal and adult organs. These datasets included four embryonic stages (4 days post fertilization (dpf) stage: “germ disc”; 6 dpf stage: “segmentation”; 10 dpf stage: “revolution” and 14 dpf stage: “pre-nymph”), heads of three different nymphal stages, nymphal gut, nymphal malpighian tubules, gills, wing pads, adult muscle, ovaries, testes, female adult brain and male and female adult heads (Figure 1c, Supplementary Table 6).

### Temporal gene expression profiles reflect life cycle adaptations

*C. dipterum* spends its life cycle in three very different environments: within the abdomen of the mother during embryonic stages (as *C. dipterum* is one of the few ovoviviparous mayfly species ^18^); freshwater streams and ponds as nymphs; and land/air as adults ^19^. To explore the expression of its genome during these three major phases, we performed soft clustering analysis of stage-specific transcriptomes using M-Fuzz ^20^ and focused on clusters expressed specifically in each of these phases (Figure 1d, Supplementary Figure 2, Supplementary Table 7). Clusters of genes transcribed preferentially during embryonic stages, such as cluster 21 and 8, showed enrichment in Gene Ontology (GO) categories that reflected the processes of embryogenesis and organogenesis happening during these stages (i.e. regulation of transcription, ectoderm development, dendrite morphogenesis, etc.). On the other hand, clusters with genes highly expressed during the aquatic phase (e.g., clusters 18 and 9) presented GO enrichment in categories that evidence the continued moulting process that mayfly nymphs undergo, such as chitin-based cuticle development. In addition, GO terms related to sensory perception of chemical stimulus or odorant binding were also enriched in these clusters (Figure 1d, Supplementary Figure 2, Supplementary Table 8). Finally, cluster 30, which contained genes with the highest expression during adulthood showed a striking enrichment of GO categories associated with visual perception. Therefore, the embryonic and postembryonic (aquatic and aerial) phases are characterized by very distinct gene expression profiles. Specifically, comparison between aquatic nymphs and aerial adults highlight expression differences that indicate very distinct sensory modalities in these two free-living phases. The aquatic phase is characterised by genes involved in the perception of chemical cues, whereas vision becomes the main sensory system during the terrestrial/aerial adult phase (Figure 1d).

### Expansion and aquatic role of Odorant Binding Proteins

Since temporal gene expression profiling indicated a prominent role of genes involved in perception of chemical cues during *C. dipterum* aquatic phase, we investigated in its genome the five main chemosensory (CS) gene families in arthropods. These include the odorant receptor/odorant receptor cofactor (OR/ORCO) and the odorant binding protein family (OBP), both of which have been suggested to play essential roles during the evolution of terrestrialization in hexapods and insects ^21-27^.

We identified 58 Gustatory Receptor (GR), 33 Ionotropic and ionotropic glutamate receptors (IR/IGluR), 15 Chemosensory Proteins (CSP), one copy of ORCO and 43 OR genes in the *C. dipterum* genome, similar to the number of genes found in the genome of the damselfly *Calopterix splendens* ^11^ for these CS gene families (Supplementary Figure 3, Supplementary Table 9). Importantly, we discovered 171 different OBP genes, which represents the largest repertoire of this family described until now, and a 57% increase with respect to the largest OBP gene complement previously described, that of the cockroach *Blattella germanica* (109 OBP genes)^22,28^. (Figure 2a, Supplementary Figure 3).

**Figure 2.**
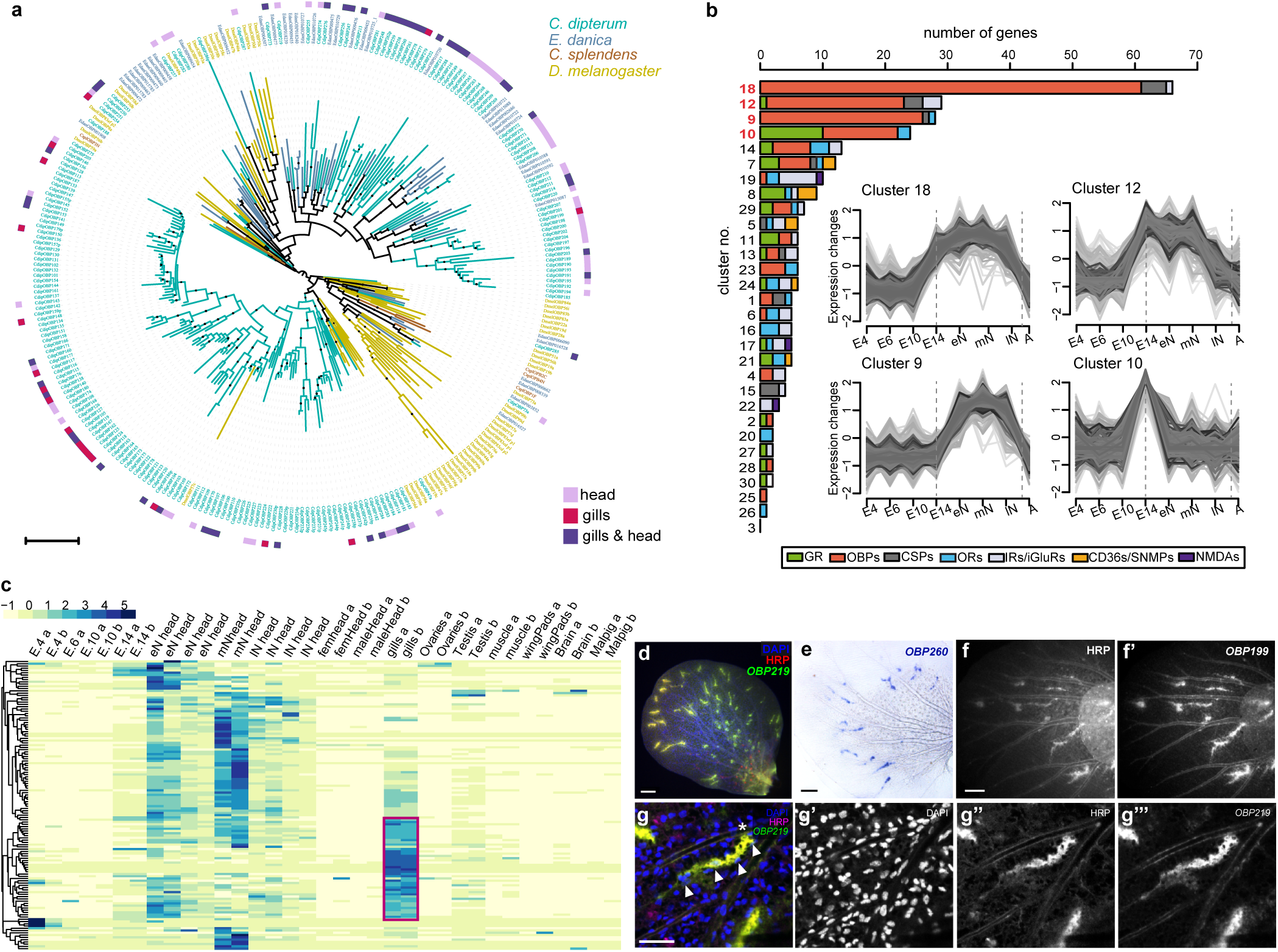
OBP gene family in *C. dipterum*. **a**, OBP phylogenetic reconstruction. *C*.*dipterum* genes are shown in green, *E. danica* in blue, *C. splendens* in brown and *D. melanogaster* in mustard. OBPs expressed specifically (according to tau values > 0,8) in nymphal heads are highlighted by pink squares, in gills in magenta and OBPs expressed both in heads and gills are shown in purple. **b**, *C. dipterum* chemosensory families distributed across M-Fuzz clusters. Clusters containing the largest amounts of chemoreceptors are no. 18, 12, 9 and 10. (E.4: 4 dpf embryo, E.6: 6 dpf embryo, E.10: 10 dpf embryo, E.14: 14 dpf embryo, eN: early nymph head, mN: mid nymph head, lN: late nymph head, A: adult head). **c**, Heat map showing enrichment of OBP expression across RNA-seq samples. Increased expression in nymphal head samples. Important fraction of OBPs are highly expressed in gills (red square). **d**, Spatial expression of *OBP219* (green) in neural structures, marked by HRP staining (in red) in abdominal gills. **e**, Spatial expression of *OBP260* in abdominal gills. **f-f’**, *OBP199* expression pattern in the gills (f’) co-localises with HRP-stained neural cells. **g**, Detail of a cluster of cells expressing *OBP219* (g’’’) and marked by HRP (g’’) in close contact with a trachea (highlighted by white asterisk). Nuclei are shown by DAPI staining (white arrowheads). Scale bars: 50 μm

Our previous GO analyses pointed to an important role of these CS genes during nymphal stages. To investigate this further, we asked in which of those M-fuzz clusters the different CS gene families were incorporated. We found that more than half of CS genes, and among these, nearly 80% of OBPs (147 out of 276 CS genes and 121 out of 152 OBPs included in the soft clustering analysis) were assigned to just four clusters (18, 12, 9 and 10), all of them related to nymphal or pre-nymphal stages. Cluster 10 contained genes that had a peak of expression at 14 dpf stage, just prior to the hatching of the first swimming nymph ^10^, and clusters 9, 12 and 18 are nymph-specific.

Next, we examined the expression of CS genes in specific tissues and organs and found that most of the CS genes were expressed in a highly tissue-specific manner, as indicated by high (> 0.8) tau values (an index used as a proxy for tissue specific expression ^29^). As expected, many CS genes were expressed in the head, where the antennae, the main olfactory organs in insects, are located. There was however, an additional major chemosensory tissue, the gills, where 34% of the CS genes were expressed, (5/13 CSP, 8/29 IR and 82/171 OBP genes, Figure 2c, Supplementary Figure 4). These included 25 gill specific OBPs, several of which (*OBP219, OBP199* and *OBP260*) were shown to be expressed in discrete cell clusters within the gills by *in situ* hybridization, often located at the branching points of their tracheal arborization (Figure 2d-g). To better characterise these clusters, we co-stained the gills with HRP, a marker of insect neurons ^30^. Remarkably, OBP-expressing cell clusters were HRP-positive, suggesting a neurosensory nature. Globally, these results strongly suggest that, beyond their respiratory role ^31^, the gills are a major chemosensory organ of the aquatic mayfly nymph (Figure 2f-g).

### Expansion of light sensing opsins in *C. dipterum*

Visual cues are essential during adulthood in mayflies (Figure 1d). During their short adult phase, they must be able to find mating partners while flying in big swarms, copulate and finally find a suitable place to deposit the eggs ^32^.

Similar to what has been observed in other diurnal insect lineages, such as Odonata ^33^, we found an expansion of Long Wave Sensitive (LWS) opsins in the *C. dipterum* genome, with four LWS opsin duplicates grouped in a genomic cluster (Figure 3a’, Supplementary Figure 5). This LWS cluster was also present within Ephemeroptera in the distantly related Ephemeridae, *Ephemera danica*, emphasizing the importance that these light-sensing molecules have had in the evolution and ecology of the entire Ephemeroptera group (Supplementary Figure 5). In addition, we also found that the blue-sensitive opsin underwent independent duplications in the Ephemeridae and Baetidae. *E. danica* has three different *blue-Ops*, while *Baetis* species with available transcriptomic data (*B. sp*. AD2013, *B. sp*. EP001, ^1^), and *C. dipterum* have two *blue-Ops*, which in the latter case are located together in tandem (Figure 3a, c).

**Figure 3.**
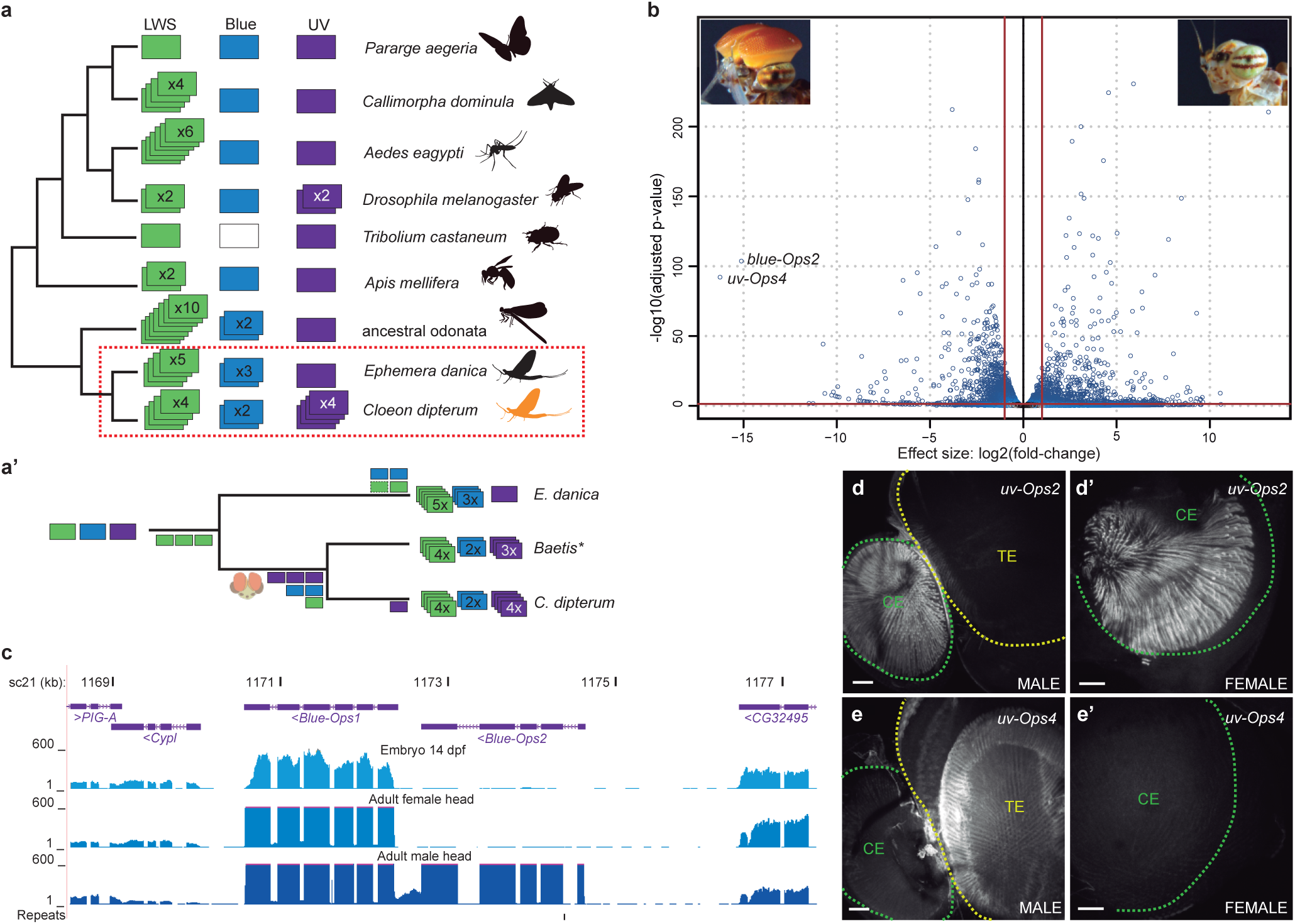
Opsin gene family in *C. dipterum*. **a**, opsin gene complement in insect phylogeny. **a’**, Evolutionary scenario of the expansions of three families of opsins in mayflies. **b**, Volcano plot showing differentially gene expression between male and female adult heads. **c**, Snapshot of Blue-Opsin cluster in *C. dipterum* genome browser. Upper and middle track show expression of *Blue-Ops1* in late embryos (st. E. 14 dpf) and adult female heads, while lower track shows high expression of both *Blue-Ops1*and *Blue-Ops2* in male adult heads. **d-d’**, Expression of *UV-Ops2* in the compound eye of a male (**d**) and a female (**d’**). **e-e’**, *UV-Ops4* high expression in the turbanate eye of a male (**e**) is not detectable in the female (**e’**).

Surprisingly, and in contrast to other insect lineages, where the UV sensing opsin (*UV-Ops*) has been usually kept as a single copy, we also found a large expansion of UV-Ops in the genome of *C. dipterum*. Duplicated UV-Ops genes were also present in different *Baetis* species (Figure 3a-a’ and Supplementary Figure 5), indicating that the baetid last common ancestor had at least three copies of UV-Ops genes, while *C. dipterum* acquired an extra duplicated *UV-Ops4*. This makes this species’ UV-Ops complement the largest one described thus far (Figure 3a).

The most salient feature of the Baetidae family is the presence of a sexually dimorphic pair of very large extra compound eyes on the dorsal part of the head, called the turbanate eyes, which develop during nymphal stages only in males ^34^. Indeed, the most highly upregulated gene in male heads compared to female heads was one of the UV-sensitive opsins, *UV-Ops4* followed by *blue-Ops2*, whose protein sequence is highly modified (Figure 3b, 3c, Supplementary Table 10). Moreover, *UV-Ops4* and *blue-Ops2* expression was only detectable from late nymph stages onwards, whereas most other opsins were expressed already in the pre-nymphal stage (14 dpf embryo, when the lateral compound eyes and ocelli, common to both males and females, start developing; Figure 3c). Their delayed expression onset and their sexual dimorphism strongly suggested that both *UV-Ops4* and *blue-Ops2* are turbanate eye-specific opsins, while the rest of *C. dipterum* opsins function in the lateral compound eyes and/or ocelli of male and females. Consistently, *in situ* hybridizations assays showed that the shared *UV-Ops2* was expressed in the compound eye of both sexes (Figure 3d), while *UV-Ops4* was predominantly expressed in the turbanate eyes of males but undetectable in females (Figure 3e). Thus, this sexually dimorphic opsin system may be of particular relevance for courtship and mating during flight.

### A conserved core set of wing genes in pterygote insects and the origin of flight

The terrestrial/aerial adult phase of *C. dipterum* is not only characterised by its visual system, but even more prominently by the key feature ancestral to all pterygote insects: the wings.

As paleopterans, mayflies can provide important insights into the origin of the genetic programs responsible for the evolution of this crucial morphological novelty. To investigate this, we first generated modules of co-regulated gene expression across several tissues (Supplementary Table 11), including developing wings, for *C. dipterum* and *D. melanogaster* (see Methods, Supplementary Data 1). To address transcriptomic conservation, we tested which modules showed significant orthologue overlap in pairwise comparisons between the two species. This analysis revealed deeply conserved co-regulated gene modules associated with muscle, gut, brain and Malpighian tubules (insect osmoregulatory organ), among others, indicating that the shared morphological features of the pterygote body plan are mirrored by deep transcriptomic conservation (Figure 4a). Remarkably, when we extend this comparison to include the centipede *Strigamia maritima*, the majority of these co-regulated modules, especially those corresponding to neural functions, muscle and gut, have been conserved as well, indicating that these genetic programs date back, at least, to the origin of Mandibulata (Supplementary Figure 6).

**Figure 4.**
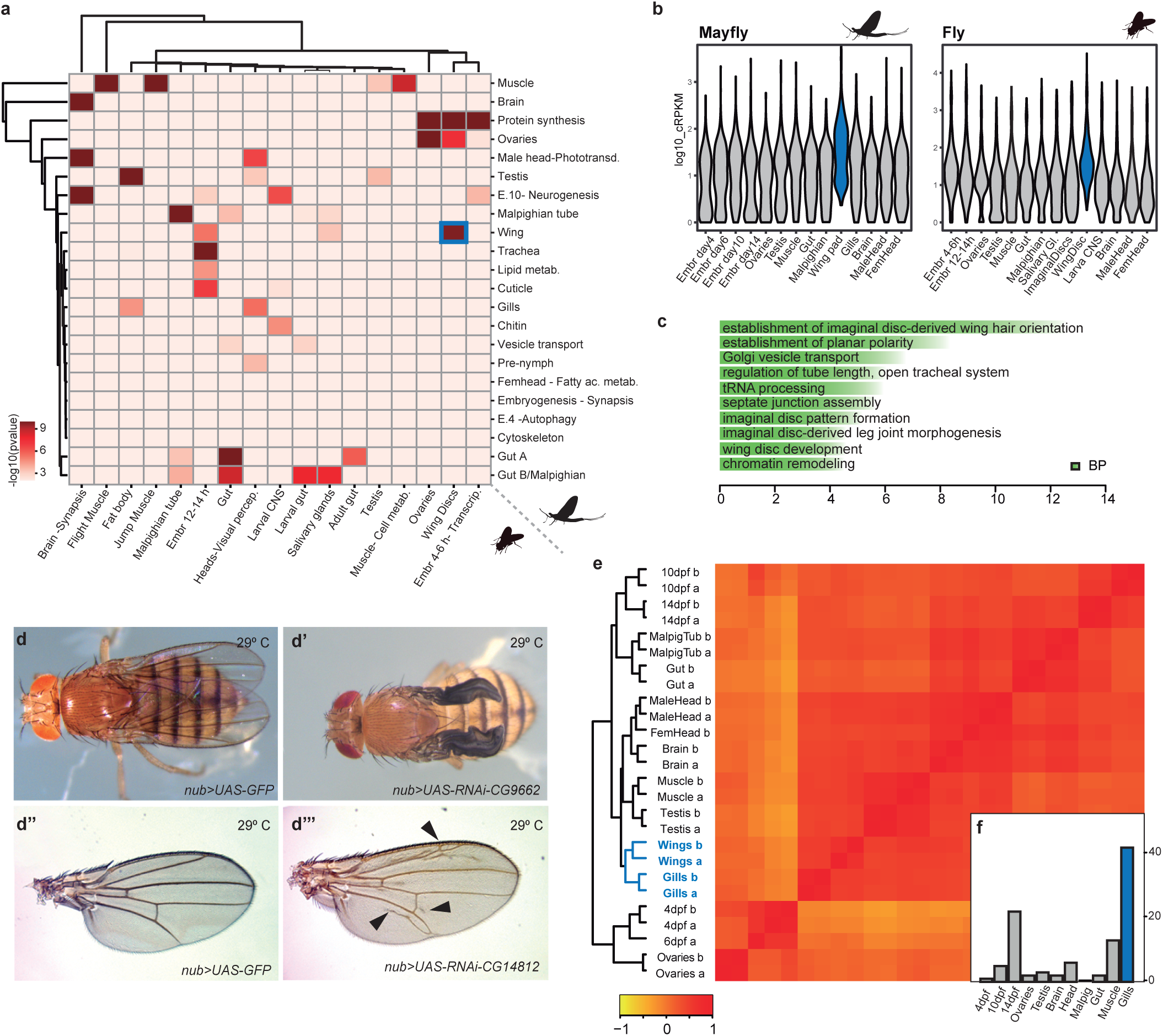
Transcriptomic conservation of wings and other insect tissues. **a**, Heat map showing the level of raw statistical significance, using upper-tail hypergeometric tests, of orthologous gene overlap between modules from *C. dipterum* (vertical) and *Drosophila* (horizontal) obtained by weighted correlation network analysis (WGCNA). **b**, Gene expression distributions (log10 cRPKMs) within the wing modules across each sample (a) for *C. dipterum* (left) and *D. melanogaster* (right). **c**, Enriched Gene Ontology terms (BP: Biological Process) of the orthologous genes shared between *C. dipterum* and *D. melanogaster* wing and wing disc modules. **d-d’’’**, Wing phenotypes resulted from knockdowns of orthologous genes shared between *C. dipterum* and *D. melanogaster* wing and wing disc modules using specific wing driver *nub-Gal4*. (d, d’’) *nub-Gal4; UAS-GFP* control fly (d) and control wing (d’’). (d’) *nub-Gal4; UAS-RNAi-CG9662* flies show severe wing phenotypes. (d’’’) *nub-Gal4; UAS-RNAi-CG14812* show extra vein tissue phenotypes. **e**, Heat map showing transcriptome datasets clustering. *C. dipterum* wing pads and gills samples cluster together. **f**, Bar plot displaying number of genes for second most expressed tissue (gills, in this case) in which the wing pad is the most expressed tissue.

Importantly, one of the highly correlated pairs of modules between mayfly and fruitfly corresponded to the wings (wing pad and wing imaginal disc modules, respectively). A total of 130 genes were shared between these modules, defining a core set of genes associated with this organ since the last common ancestor of pterygotes. This gene set exhibited an enrichment in GO terms that corresponded to wing development (Figure 4c), and in agreement with this, some of these orthologues have been shown to participate in wing development in *Drosophila* (e.g. *abnormal wing discs* (*awd*), *inturned* (*in*) or *wing blister* (*wb*), etc.). In addition, among them there were also numerous genes (96 genes out of 130, Supplementary Table 12) for which no previous wing function has been described so far. However, our comparative approach strongly indicated a putative function in wing development. We functionally tested this hypothesis for eight of these genes in *Drosophila* (Supplementary Table 13), by RNAi knockdown assays using the wing specific *nubbin-GAL4* driver ^35^. Indeed, these experiments resulted in wing phenotypes in all cases (8/8) and in particular, abnormal wing venation (7/8) (Figure 4d, Supplementary Figure 6).

We next asked if the origin of novel traits in particular lineages, including the wings in pterygotes, might have been coupled to the appearance of new genes. To this end, we classified orthologous genes according to their evolutionary origin and checked whether there was any correlation between their appearance and the tissues in which they were mainly expressed, based on our previous WGCNA modules (Figure 4a, Supplementary Data 1) and tau index analyses (Supplementary Figure 6). In general, our results were in agreement with previously described phylostratigraphic patterns, in which older genes exhibit broad expression patterns while novel genes are expressed in a tissue-specific manner ^36,37^. We did not observe any enrichment of new genes expressed in the wings in the Pterygota stratum that could be linked to the origin of this morphological novelty, despite the high number of novel genes that were gained in this lineage (Figure 1a, Supplementary Figure 1). These results suggested that the transcriptomic program responsible for the wings was assembled from genes that were already forming part of pre-existing gene networks in other tissues^38^. Therefore, we investigated the similarities between the transcriptomic programs of wings and other mayfly tissues (Figure 4e-f). Clustering *C. dipterum* genes based on their expression across the different tissues, organs and developmental stages revealed that gills were the most closely related organ to (developing) wings (Figure 4d). In fact, gills were the tissue that shared most specific genes with wing pads (42/98, 43%), as assessed by looking for genes with high expression only in wings and an additional tissue (see Methods, Figure 4e, Supplementary Figure 6d, e), confirming the transcriptional similarities between gills and wing pads in mayflies.

## Discussion

*C. dipterum* belongs to a critical lineage within insect phylogeny. Together with the possibility of culturing it continuously and its life cycle of about 45 days, *C. dipterum*’s genome sequence and transcriptome datasets presented here provide the foundations for the exploration of a number of critical evolutionary, developmental and physiological aspects of the biology of insects. When interrogated about the expression of the genome associated with the two very disparate environments that mayflies adapted to (aquatic and aerial), the data suggest a sensory specialization, with nymphs predominantly using chemical stimuli, whereas during the adult stages individuals would rely predominantly on their visual system. This sensory specialization has involved very specific genomic changes.

Arthropods perceive different chemical environmental cues, such as pheromones, food or the presence of predators, using different families of chemosensory proteins specially tuned to the structural and chemical characteristics of those cues (e.g. volatile, hydro soluble, etc.). Accordingly, CS families in arthropods have undergone multiple lineage-specific changes based on adaptations to the myriad of ecological niches. The possible co-option of OBPs by Hexapoda^24^ and the appearance of ORs in insects led to the idea that these two families evolved as an adaptation to terrestrial life and that their main function is to perceive smells in the aerial media. However, the role of some OBP genes is still unclear, and functions other than olfaction have been proposed for OBPs in terrestrial insects^39^. The finding of OBP-expressing organs in gills-apart from the heads-exhibiting neural markers, and associated with the trachea, together with the specific expression of some chemoreceptors in gills (Supplementary Figure 3) indicates that gills may be a prominent chemosensory organ in the mayfly. In fact, their large combined surface make them especially apt for this function. Further, the presence of these chemosensory structures in the gills challenges the classic idea of gills as exclusively respiratory organs^31^.

Living in two media (water and air) and the new ability to fly must have imposed a number of very specific requirements on the mayfly visual system, from vision in two refractive index media, or the use of novel visual cues for mating in the air, to an increase of the speed of visual information flow. For instance, insects evolved their visual systems and different types of light sensing opsins to navigate during sunlight or moonlight illumination or to use polarized light to obtain important information from the environment ^33,40^. Multiple insect lineages expanded the long-wavelength sensing (LWS) opsin complement. In a similar way, Ephemeroptera show an ancestral expansion of LWS opsins, located in a cluster. By contrast, most insect groups have kept blue and UV opsins as a single copy (Figure 3a). However, we found that the Baetidae family, where *C. dipterum* is included, has duplicated these two light sensing gene families, with the largest UV opsin expansion described so far. The expansion of this particular type of opsins in Baetidae could be related to the origin and evolution of the male-specific turbanate eyes in this family of mayflies, since we observed that *UV-Ops4* is exclusively expressed in this novel sexually dimorphic visual system (Figure 3d). UV opsins are usually more prominent in the dorsal part of insect eyes, due to their direct and frontal/upward position towards the sunlight^40-42^. Since it has been suggested that males use their dorsal turbanate eyes to locate females flying above their swarm, sexual selection might have played an important role in the expansion of UV and blue opsins and their specific expression in turbanate eyes of males.

Our transcriptomic comparisons between *C. dipterum, Drosophila* and *Strigamia* revealed for the first time high conservation of gene expression in multiple homologous arthropod tissues and organs, indicating that deep conservation of transcriptomic programs may be a common signature in different phyla^43,44^. We identified a core set of developing wing genes ancestral to all pterygote insects. Notably, when we tested functionally some of these genes with no previously described functions, all of them showed wing phenotypes, and most often defects in veins. Veins are thickenings of the wing epithelium that provide it with sufficient rigidity for flight and which are unique anatomical wing features, consistent with the specificity of these transcriptional programs. Overall, our results attest to the strong predictive power of our evolutionary comparative approach to infer function of conserved genes.

Several hypotheses to explain the origin of wings have been put forward since the last century^4,5,7-9,45^, including the pleural origin hypothesis in which the wings would be the thoracic serial homologues of the abdominal gills. Our results uncovered expression similarities between gills and wings, which would be consistent with both pleural and dual scenarios, in which ancestral thoracic gills would contribute to wing structures. Alternatively, in a scenario in which gills and the aquatic nymphs would not be the ancestral state to pterygotes^46,47^ these similarities could represent gene co-option between these two organs. Whatever the case the transcriptomic similarities observed between gills and wings suggest that they share a common genetic program.

## Methods

### Genome sequencing and assembly

Genomic DNA was extracted from *C. dipterum* adult males from an inbred line kept in the laboratory for seven generations^10^. Illumina and Nanopore technologies were used to sequence the genome (Supplementary Table 1-3). The assembly was generated by hybrid assembly using MaSuRCA v3.2.3 ^48,49^ with 95.9x short-read Illumina short-read and 36.3x ONT long-read coverage.

### RNA sequencing and assembly

35 RNA-seq datasets of multiple developmental stages (four embryonic stages) and dissected tissues and organs (nymphal and adult dissected organs and whole heads) were generated using the Illumina technology. Samples were processed immediately after dissection and RNA was extracted using RNeasy Mini Kit (Qiagen) or RNAqueous™-Micro Total RNA Isolation Kit (Ambion) following manufacturers instructions. Single-end and paired-end libraries were generated using Illumina (TruSeq) RNA-Seq kit.

### Genome annotation

Reads from all paired-end transcriptomes were assembled altogether using Trinity ^50^ and subsequently aligned to the genome using the PASA pipeline ^51^. High-quality transcripts were selected to build a Hidden-Markov profile for *de novo* gene prediction in Augustus ^52^.

Reads were aligned to the genome using STAR ^53^ and transcriptome assembled using Stringtie ^54^. The assemblies were merged using Taco ^55^. Consensus transcriptome assembly and splice-junctions were converted into hints and provided to Augustus gene prediction tool which yielded 16364 evidence-based gene models. These models contain 4308 non-redundant PFAM models as assigned using the PfamScan tool. RepeatModeller was used to identify repeats in *C. dipterum* assembled genome.

We integrated these RNA-seq data into a UCSC Genome Browser track hub together with two additional tracks, insect sequence conservation and annotation of repetitive elements.

### Comparative transcriptomics

Mfuzz software^20^ was used to perform soft clustering of genes according to developmental and life history expression dynamics using normalised read counts. We selected eight developmental and post-embryonic stages. We used DESeq2 R package^56^ to analyse differential gene expression between male and female adult heads.

We used the cRPKM metric (corrected-for-mappability Reads Per Kilobasepair of uniquely mappable positions per Million mapped reads^57^ to perform WGCNA gene expression analyses. We performed the analyses using as datasets genes that were present in a least two species in our family reconstructions and showed variance across samples (coef. var ≥ 1). Each module was designated with a tissue or a biological/molecular category. Finally, we analysed the overlap between homologous groups for each pair of modules for each of the species in a pairwise manner. To evaluate significance we performed hypergeometric tests.

### Gene orthology

To obtain phylogeny-based orthology relationships between different taxa, the predicted proteomes of 14 species (Supplementary Table 4) representing major arthropod lineages and outgroups were used as input for OrthoFinder2 ^58^.

### Chemosensory genes identification and phylogeny

We created a dataset containing well-annotated members of the chemosensory (CS) gene family (i.e., GR, OR, IR/iGluR, OBP and CSP) from a group of representative insects (Supplementary Table 9^11,26-28,59-64^). In addition, we constructed specific HMM profiles for each CS gene family based on their Pfam profiles (see Supplementary table 1 in ^65^). Briefly, we performed iterative rounds of BLASTP and HMMER searches against the annotated proteins of *C. dipterum*, curating incorrect gene models; and TBLASTN against the genomic sequence to identify CS proteins. Finally, we obtained a curated GTF containing the annotation for each CS gene family.

### Opsins identification and phylogeny

A total 1247 opsins from all the major metazoans groups were used as seed in the BLAST research the predicted protein sequences of *Cloeon dipterum, Ephemera danica* (https://www.hgsc.bcm.edu/arthropods/mayfly-genome-project) *and Ladona fulva* (https://www.hgsc.bcm.edu/arthropods/scarce-chaser-genome-project). To this set of sequences, additional mayfly LWS, UV and Blue opsin sequences from transcriptome assemblies for *Baetis sp. EP001* and *Epeorus sp. EP006* ^33,66^, for *Baetis sp. AD2013* from ^1^ and *Baetis rhodani* shotgun whole genome assembly ^67^ were obtained. Mayfly LWS, UV and Blue opsin sequences were carefully inspected and curated (Supplementary Table 15). To this dataset, we finally added the sequences for an additional 27 species covering seven orders were retrieved from ^68^. The alignment was performed using MAFFT with default parameters. Phylogenetic reconstruction was performed using Ultrafast bootstrap with 1000 replicates, aLRT Bootstrap and aBayes ^69,70^. In all the phylogenetic analyses the trees were rooted using the melatonin receptor which represent opsin closest outgroup ^71^.

### In situ hybridizations

Specific primers were designed to generate DIG-labelled probes against *OBP260, OBP219, OBP199, UV-Ops2* and *UV-Ops4* (Supplementary Table 14). After o.n. fixation of the gills or heads in FA 4% at 4° C, post-fixated gills and retinas were treated with Proteinase K for 15 min. The hybridization was carried out at 60° C o.n. Tissues were incubated with anti-digoxigenin-POD at 4° C o.n. and with 1:100 TSA in borate buffer for 1 hour. Leica SPE confocal microscope was used to acquire images that were processed with Fiji ^72^.

### RNAi assays in Drosophila wings

VDRC lines (see Supplementary Table 13) were crossed to *yw; nub-Gal4; +* line to expressed the RNAi constructs specifically in the wing. Crosses were kept at 25° C for 48 hours and then switched to 29° C. Wings were dissected from adult females and mounted in Hoyer’s/Lactic acid (1:1) medium prior to microscopic analysis and imaging.

## Acknowledgments

We thank Yamile Márquez, Juan J. Tena, Alejandro Gil and Rafael D. Acemel for help in bioinformatics analyses. This project has been mainly funded by the European Union’s Horizon 2020 research and innovation program under the Marie Sklodowska-Curie Grant Agreement 657732 to I.A., Grant BFU2015-66040-P to F.Ca. and institutional Grant MDM-2016-0687 (MINECO, Spain).

## Author contributions

T.A., M.Gu., A.dM., M.I., I.M., F.Ca. and I.A. contributed to concept and study design. J.V., A. dM., F.M., P.F., C.W., R.F., P. M., F.Cr., J.G.G., M. Gu., T.A., C. V-C., J.G., J.P., R.L., J.R., A.S-G., M.I., I.M., F.Ca., and I. A. performed computational analyses and data interpretation. K.D., S.A., B.M., F.Cr., J.G.G., M. Gu., T.A., M.I., F.Ca. and I.A. obtained biological material and generated next-generation sequencing data. G.M., A.A. and I.A. performed in situ hybridizations and knockdown assays. I.A. and F.Ca. coordinated the project and obtained funding. I.A. and F.Ca. wrote the main text with the help of I.M., and inputs from all authors. Arthropods illustrations by I.A., released under a Creative Commons Attribution (CC-BY) License.

## Competing interests

The authors declare no competing interests.

## Data Availability

The sequence reads and the genome assembly have been deposited in the European Nucleotide Archive (ENA) with the project accessions PRJEB34721 and PRJEB35103.

## Supplementary information

This file contains Supplementary text and data, supplementary references and supplementary Tables 1-15

**Supplementary Data 1**. WGCNA modules

**Supplementary Table 1**. Sequencing table statistics

**Supplementary Table 2**. Kraken contaminats in mayfly genome assembly

**Supplementary Table 3**. Contaminants

**Supplementary Table 4**. Orthofinder species

**Supplementary Table 5**. Orthofinder stats orthologues

**Supplementary Table 6**. RNA samples sequenced

**Supplementary Table 7**. M-fuzz stages

**Supplementary Table 8**. M-fuzz data

**Supplementary Table 9**. Chemosensory complement in insects

**Supplementary Table 10**. DESeq results female vs male heads

**Supplementary Table 11**. WGCNA genes per species

**Supplementary Table 12**. Shared genes between *C. dipterum* and *D. melanogaster* wing pad and wing disc WGCNA modules

**Supplementary Table 13**. RNAi lines used in *D. melanogaster*

**Supplementary Table 14**. Primers used to design RNA probes for in situ hybridization

**Supplementary Table 15**. Opsins identified in Ephemeroptera

## References

1 Misof, B. et al. Phylogenomics resolves the timing and pattern of insect evolution. Science 346, 763–767, doi: 10.1126/science.1257570 (2014).

2 Belles, X. The innovation of the final moult and the origin of insect metamorphosis. Philosophical Transactions of the Royal Society B: Biological Sciences 374, 20180415, doi: 10.1098/rstb.2018.0415 (2019).

3 Engel, M. S., Davis, S. R. & Prokop, J. in Arthropod Biology and Evolution: Molecules, Development, Morphology (eds Alessandro Minelli, Geoffrey Boxshall, & Giuseppe Fusco) 269–298 (Springer Berlin Heidelberg, 2013).

4 Averof, M. & Cohen, S. M. Evolutionary origin of insect wings from ancestral gills. Nature 385, 627–630 (1997).

5 Clark-Hachtel, C. M. & Tomoyasu, Y. Exploring the origin of insect wings from an evo-devo perspective. Current Opinion in Insect Science 13, 77–85, doi: http://doi.org/10.1016/j.cois.2015.12.005 (2016).

6 Hamilton, K. G. A. The insect wing. Part 1. Origin and development of wings from notal lobes. 1971, 44:421-433. Hamilton, K. G. A. J Kansas Entomol Soc 44:421–433. (1971).

7 Kukalova-Peck, J. Origin and evolution of insect wings and their relation to metamorphosis, as documented by the fossil record. Journal of Morphology 156, 53–125, doi: 10.1002/jmor.1051560104 (1978).

8 Ohde, T., Yaginuma, T. & Niimi, T. Insect Morphological Diversification Through the Modification of Wing Serial Homologs. Science 340, 495–498, doi: 10.1126/science.1234219 (2013).

9 Wigglesworth, V. B. Evolution of Insect Wings and Flight. Nature 246, 127–129, doi: 10.1038/246127a0 (1973).

10 Almudi, I. et al. Establishment of the mayfly Cloeon dipterum as a new model system to investigate insect evolution. Evodevo 10, 6, doi: 10.1186/s13227-019-0120-y (2019).

11 Ioannidis, P. et al. Genomic Features of the Damselfly Calopteryx splendens Representing a Sister Clade to Most Insect Orders. Genome Biol Evol 9, 415–430, doi: 10.1093/gbe/evx006 (2017).

12 Armisén, D. et al. The genome of the water strider Gerris buenoi reveals expansions of gene repertoires associated with adaptations to life on the water. BMC Genomics 19, 832, doi: 10.1186/s12864-018-5163-2 (2018).

13 Wu, C. et al. Analysis of the genome of the New Zealand giant collembolan (Holacanthella duospinosa) sheds light on hexapod evolution. BMC Genomics 18, 795, doi: 10.1186/s12864-017-4197-1 (2017).

14 Panfilio, K. A. et al. Molecular evolutionary trends and feeding ecology diversification in the Hemiptera, anchored by the milkweed bug genome. Genome Biology 20, 64, doi: 10.1186/s13059-019-1660-0 (2019).

15 Petersen, M. et al. Diversity and evolution of the transposable element repertoire in arthropods with particular reference to insects. BMC Evolutionary Biology 19, 11, doi: 10.1186/s12862-018-1324-9 (2019).

16 Parra, G., Bradnam, K. & Korf, I. CEGMA: a pipeline to accurately annotate core genes in eukaryotic genomes. Bioinformatics 23, 1061–1067, doi: 10.1093/bioinformatics/btm071 (2007).

17 Simao, F. A., Waterhouse, R. M., Ioannidis, P., Kriventseva, E. V. & Zdobnov, E. M. BUSCO: assessing genome assembly and annotation completeness with single-copy orthologs. Bioinformatics 31, 3210–3212, doi: 10.1093/bioinformatics/btv351 (2015).

18 Berner, L. Ovoviviparous Mayflies in Florida. The Florida Entomologist 24, 32–34, doi: 10.2307/3491942 (1941).

19 Clifford, H. F. Life Cycles of mayflies (Ephemeroptera), with special reference to voltinism. Quaestiones Entomologicae 18, 15–90 (1982).

20 Kumar, L. & E Futschik, M. Mfuzz: a software package for soft clustering of microarray data. Bioinformation 2, 5–7, doi: 10.6026/97320630002005 (2007).

21 Eyun, S.-i. et al. Evolutionary History of Chemosensory-Related Gene Families across the Arthropoda. Molecular biology and evolution 34, 1838–1862, doi: 10.1093/molbev/msx147 (2017).

22 Robertson, H. M., Baits, R. L., Walden, K. K. O., Wada-Katsumata, A. & Schal, C. Enormous expansion of the chemosensory gene repertoire in the omnivorous German cockroach Blattella germanica. Journal of Experimental Zoology Part B: Molecular and Developmental Evolution 330, 265–278, doi: 10.1002/jez.b.22797 (2018).

23 Robertson, H. M., Warr, C. G. & Carlson, J. R. Molecular evolution of the insect chemoreceptor gene superfamily in Drosophila melanogaster. Proceedings of the National Academy of Sciences of the United States of America 100 Suppl 2, 14537–14542, doi: 10.1073/pnas.2335847100 (2003).

24 Vizueta, J., Rozas, J. & Sánchez-Gracia, A. Comparative Genomics Reveals Thousands of Novel Chemosensory Genes and Massive Changes in Chemoreceptor Repertories across Chelicerates. Genome Biology and Evolution 10, 1221–1236, doi: 10.1093/gbe/evy081 (2018).

25 Missbach, C., Vogel, H., Hansson, B. S. & Große-Wilde, E. Identification of Odorant Binding Proteins and Chemosensory Proteins in Antennal Transcriptomes of the Jumping Bristletail Lepismachilis y-signata and the Firebrat Thermobia domestica: Evidence for an Independent OBP–OR Origin. Chemical Senses 40, 615–626, doi: 10.1093/chemse/bjv050 (2015).

26 Brand, P. et al. The origin of the odorant receptor gene family in insects. eLife 7, e38340, doi: 10.7554/eLife.38340 (2018).

27 Missbach, C. et al. Evolution of insect olfactory receptors. Elife 3, e02115, doi: 10.7554/eLife.02115 (2014).

28 Harrison, M. C. et al. Hemimetabolous genomes reveal molecular basis of termite eusociality. Nature Ecology & Evolution 2, 557–566, doi: 10.1038/s41559-017-0459-1 (2018).

29 Kryuchkova-Mostacci, N. & Robinson-Rechavi, M. Tissue-Specificity of Gene Expression Diverges Slowly between Orthologs, and Rapidly between Paralogs. PLOS Computational Biology 12, e1005274, doi: 10.1371/journal.pcbi.1005274 (2016).

30 Jan, L. Y. & Jan, Y. N. Antibodies to horseradish peroxidase as specific neuronal markers in Drosophila and in grasshopper embryos. Proceedings of the National Academy of Sciences of the United States of America 79, 2700–2704, doi: 10.1073/pnas.79.8.2700 (1982).

31 Wingfield, C. A. Function of the Gills of the Mayfly Nymph, Cloeon dipterum. Nature 140, 27–27, doi: 10.1038/140027a0 (1937).

32 Allan, J. D. & Flecker, A. S. The mating biology of a mass-swarming mayfly. Animal Behaviour 37, 361–371, doi: https://doi.org/10.1016/0003-3472(89)90084-5 (1989).

33 Futahashi, R. et al. Extraordinary diversity of visual opsin genes in dragonflies. Proceedings of the National Academy of Sciences 112, E1247–E1256, doi: 10.1073/pnas.1424670112 (2015).

34 Zimmer, C. Die Facettenaugen der Ephemeriden. Z. Wiss. Zool. 63, 236–261 (1898).

35 Calleja, M., Moreno, E., Pelaz, S. & Morata, G. Visualization of Gene Expression in Living Adult *Drosophila*. Science 274, 252–255, doi: 10.1126/science.274.5285.252 (1996).

36 Sebé-Pedrós, A. et al. High-Throughput Proteomics Reveals the Unicellular Roots of Animal Phosphosignaling and Cell Differentiation. Developmental Cell 39, 186–197, doi: 10.1016/j.devcel.2016.09.019 (2016).

37 Tautz, D. & Domazet-Lošo, T. The evolutionary origin of orphan genes. Nature Reviews Genetics 12, 692–702, doi: 10.1038/nrg3053 (2011).

38 Almudí, I. & Pascual-Anaya, J. in Old Questions and Young Approaches to Animal Evolution (eds José M. Martín-Durán & Bruno C. Vellutini) 107–132 (Springer International Publishing, 2019).

39 Pelosi, P., Iovinella, I., Felicioli, A. & Dani, F. R. Soluble proteins of chemical communication: an overview across arthropods. Front Physiol 5, 320, doi: 10.3389/fphys.2014.00320 (2014).

40 Heinloth, T., Uhlhorn, J. & Wernet, M. F. Insect Responses to Linearly Polarized Reflections: Orphan Behaviors Without Neural Circuits. Frontiers in Cellular Neuroscience 12, doi: 10.3389/fncel.2018.00050 (2018).

41 Sancer, G. et al. Modality-Specific Circuits for Skylight Orientation in the Fly Visual System. Current Biology 29, 2812-2825.e2814, doi: https://doi.org/10.1016/j.cub.2019.07.020 (2019).

42 Hilbrant, M. et al. Sexual dimorphism and natural variation within and among species in the Drosophilaretinal mosaic. BMC Evolutionary Biology 14, 240, doi: 10.1186/s12862-014-0240-x (2014).

43 Barbosa-Morais, N. L. et al. The Evolutionary Landscape of Alternative Splicing in Vertebrate Species. Science 338, 1587–1593, doi: 10.1126/science.1230612 (2012).

44 Marlétaz, F. et al. Amphioxus functional genomics and the origins of vertebrate gene regulation. Nature 564, 64–70, doi: 10.1038/s41586-018-0734-6 (2018).

45 Niwa, N. et al. Evolutionary origin of the insect wing via integration of two developmental modules. Evolution & Development 12, 168–176, doi: 10.1111/j.1525-142X.2010.00402.x (2010).

46 Lozano-Fernandez, J. et al. Pancrustacean Evolution Illuminated by Taxon-Rich Genomic-Scale Data Sets with an Expanded Remipede Sampling. Genome Biology and Evolution 11, 2055–2070, doi: 10.1093/gbe/evz097 (2019).

47 Wipfler, B. et al. Evolutionary history of Polyneoptera and its implications for our understanding of early winged insects. Proceedings of the National Academy of Sciences 116, 3024–3029, doi: 10.1073/pnas.1817794116 (2019).

48 Zimin, A. V. et al. Hybrid assembly of the large and highly repetitive genome of Aegilops tauschii, a progenitor of bread wheat, with the mega-reads algorithm. bioRxiv, doi: 10.1101/066100 (2016).

49 Zimin, A. V. et al. The MaSuRCA genome assembler. Bioinformatics 29, 2669–2677, doi: 10.1093/bioinformatics/btt476 (2013).

50 Grabherr, M. G. et al. Full-length transcriptome assembly from RNA-Seq data without a reference genome. Nat Biotechnol 29, 644–652, doi: 10.1038/nbt.1883 (2011).

51 Hass, J., Blaschke, S., Rammsayer, T. & Herrmann, J. M. A neurocomputational model for optimal temporal processing. J Comput Neurosci 25, 449–464, doi: 10.1007/s10827-008-0088-4 (2008).

52 Stanke, M., Tzvetkova, A. & Morgenstern, B. AUGUSTUS at EGASP: using EST, protein and genomic alignments for improved gene prediction in the human genome. Genome Biol 7 Suppl 1, S11 11–18, doi: 10.1186/gb-2006-7-s1-s11 (2006).

53 Dobin, A. et al. STAR: ultrafast universal RNA-seq aligner. Bioinformatics 29, 15–21, doi: 10.1093/bioinformatics/bts635 (2013).

54 Pertea, M. et al. StringTie enables improved reconstruction of a transcriptome from RNA-seq reads. Nat Biotechnol 33, 290–295, doi: 10.1038/nbt.3122 (2015).

55 Niknafs, Y. S., Pandian, B., Iyer, H. K., Chinnaiyan, A. M. & Iyer, M. K. TACO produces robust multisample transcriptome assemblies from RNA-seq. Nat Methods 14, 68–70, doi: 10.1038/nmeth.4078 (2017).

56 Love, M. I., Huber, W. & Anders, S. Moderated estimation of fold change and dispersion for RNA-seq data with DESeq2. Genome Biol 15, 550, doi: 10.1186/s13059-014-0550-8 (2014).

57 Labbé, R. M. et al. A Comparative Transcriptomic Analysis Reveals Conserved Features of Stem Cell Pluripotency in Planarians and Mammals. STEM CELLS 30, 1734–1745, doi: 10.1002/stem.1144 (2012).

58 Emms, D. M. & Kelly, S. OrthoFinder: phylogenetic orthology inference for comparative genomics. bioRxiv, 466201, doi: 10.1101/466201 (2019).

59 Benton, R., Vannice, K. S., Gomez-Diaz, C. & Vosshall, L. B. Variant ionotropic glutamate receptors as chemosensory receptors in Drosophila. Cell 136, 149–162, doi: 10.1016/j.cell.2008.12.001 (2009).

60 Vogt, R. G. et al. The insect SNMP gene family. Insect Biochem Mol Biol 39, 448–456, doi: 10.1016/j.ibmb.2009.03.007 (2009).

61 Vieira, F. G. & Rozas, J. Comparative genomics of the odorant-binding and chemosensory protein gene families across the Arthropoda: origin and evolutionary history of the chemosensory system. Genome Biol Evol 3, 476–490, doi: 10.1093/gbe/evr033 (2011).

62 Kirkness, E. F. et al. Genome sequences of the human body louse and its primary endosymbiont provide insights into the permanent parasitic lifestyle. Proc Natl Acad Sci U S A 107, 12168–12173, doi: 10.1073/pnas.1003379107 (2010).

63 Terrapon, N. et al. Molecular traces of alternative social organization in a termite genome. Nat Commun 5, 3636, doi: 10.1038/ncomms4636 (2014).

64 Wu, C. et al. Analysis of the genome of the New Zealand giant collembolan (Holacanthella duospinosa) sheds light on hexapod evolution. BMC Genomics 18, 795–795, doi: 10.1186/s12864-017-4197-1 (2017).

65 Frias-Lopez, C. et al. Comparative analysis of tissue-specific transcriptomes in the funnel-web spider Macrothele calpeiana (Araneae, Hexathelidae). PeerJ 3, e1064, doi: 10.7717/peerj.1064 (2015).

66 Suvorov, A. et al. Opsins have evolved under the permanent heterozygote model: insights from phylotranscriptomics of Odonata. Mol Ecol 26, 1306–1322, doi: 10.1111/mec.13884 (2017).

67 Macdonald, H. C., Ormerod, S. J. & Bruford, M. W. Enhancing capacity for freshwater conservation at the genetic level: a demonstration using three stream macroinvertebrates. Aquatic Conservation: Marine and Freshwater Ecosystems 27, 452–461, doi: 10.1002/aqc.2691 (2017).

68 Feuda, R., Marletaz, F., Bentley, M. A. & Holland, P. W. Conservation, Duplication, and Divergence of Five Opsin Genes in Insect Evolution. Genome Biol Evol 8, 579–587, doi: 10.1093/gbe/evw015 (2016).

69 Hoang, D. T., Chernomor, O., von Haeseler, A., Minh, B. Q. & Vinh, L. S. UFBoot2: Improving the Ultrafast Bootstrap Approximation. Molecular biology and evolution 35, 518–522, doi: 10.1093/molbev/msx281 (2017).

70 Anisimova, M., Gil, M., Dufayard, J.-F., Dessimoz, C. & Gascuel, O. Survey of branch support methods demonstrates accuracy, power, and robustness of fast likelihood-based approximation schemes. Systematic biology 60, 685–699, doi: 10.1093/sysbio/syr041 (2011).

71 Feuda, R., Hamilton, S. C., McInerney, J. O. & Pisani, D. Metazoan opsin evolution reveals a simple route to animal vision. Proceedings of the National Academy of Sciences 109, 18868–18872, doi: 10.1073/pnas.1204609109 (2012).

72 Schindelin, J. et al. Fiji: an open-source platform for biological-image analysis. Nature Methods 9, 676–682, doi: 10.1038/nmeth.2019 (2012).

