## Supplementary figures and images for "Genomic adaptations to aquatic and aerial life in mayflies and the origin of wings in insects"

### Supplementary Figure 1

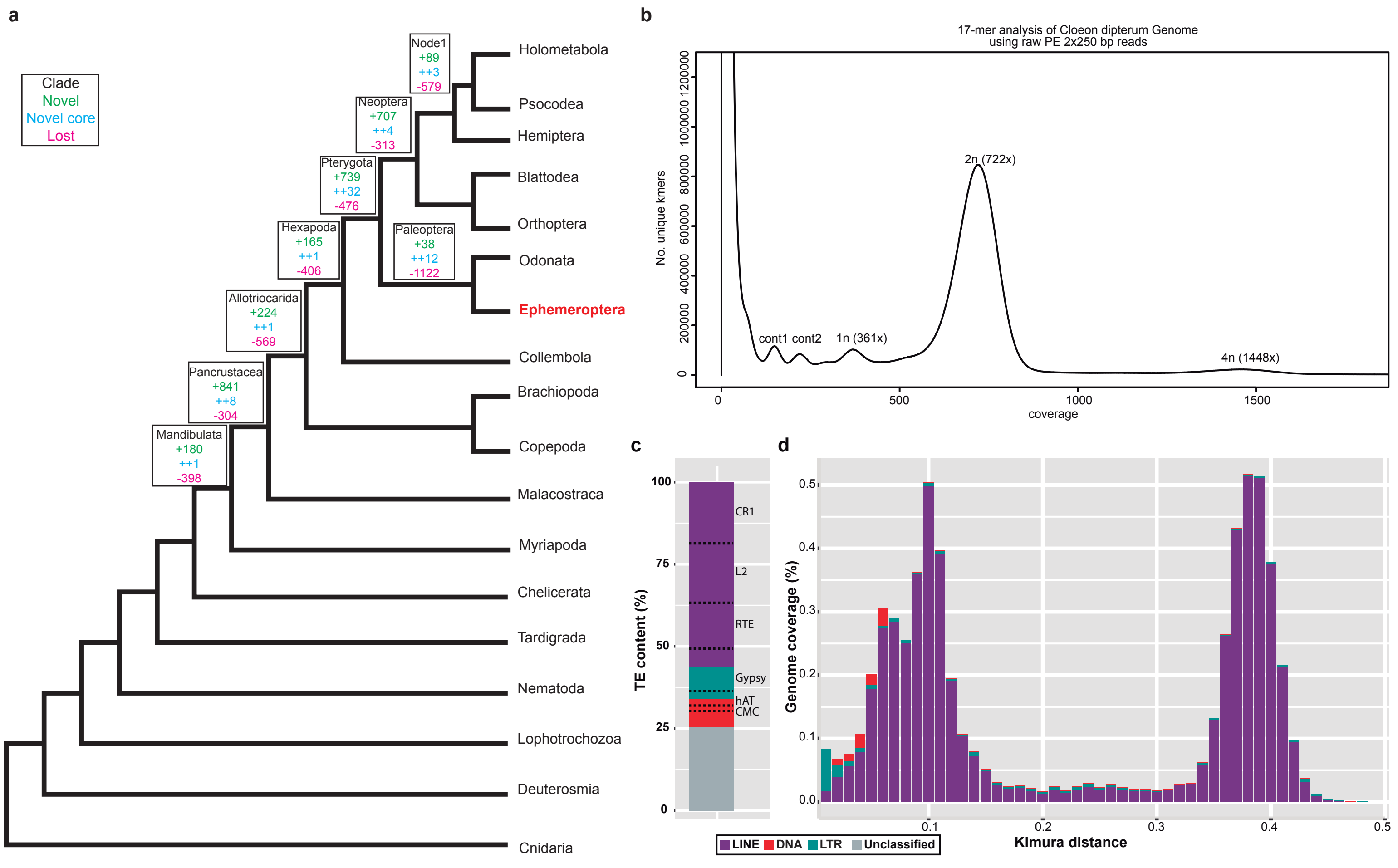
