## Supplementary Figure 2 for "Genomic adaptations to aquatic and aerial life in mayflies and the origin of wings in insects"

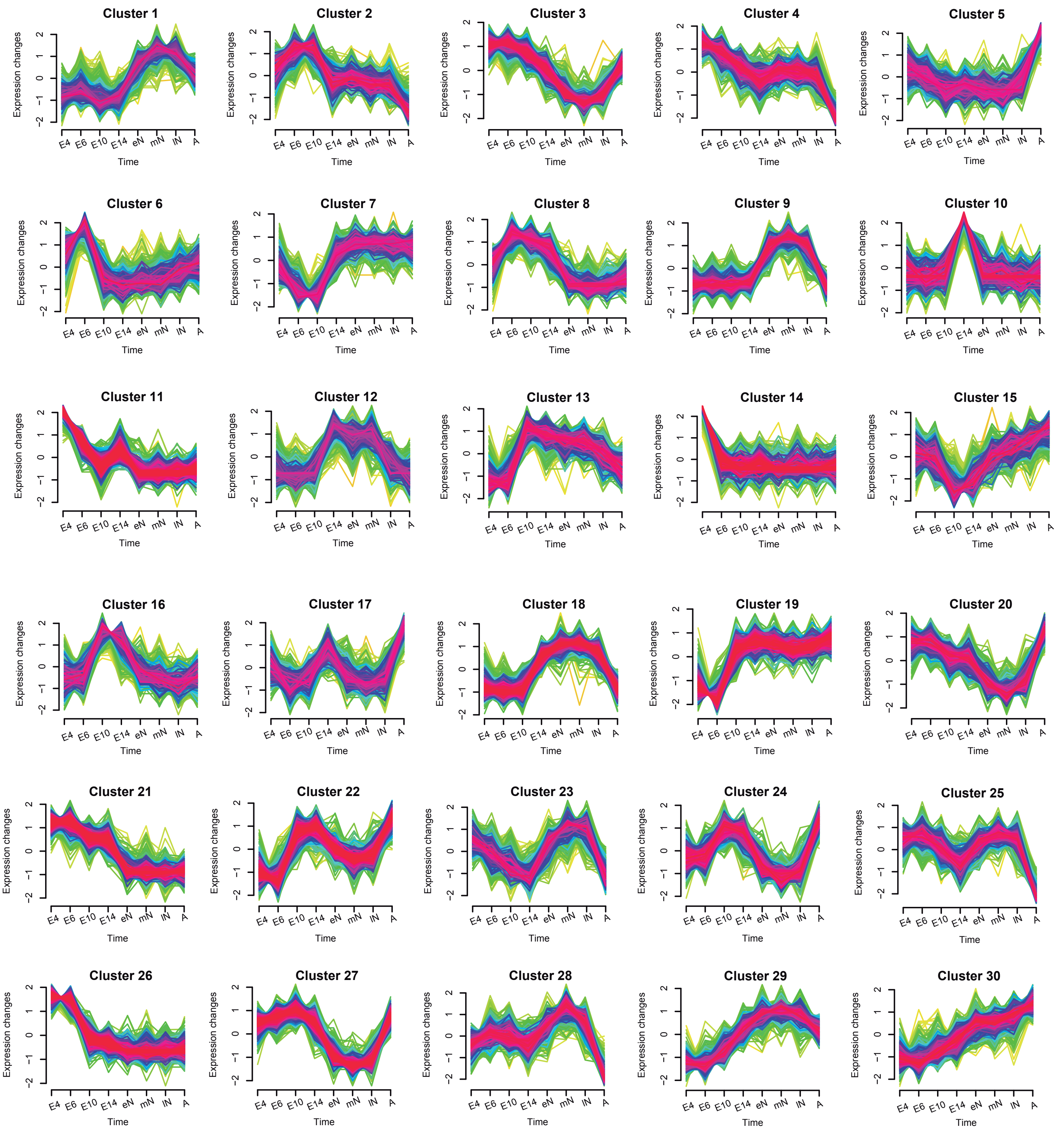


**Supplementary Figure 2.** Dynamics of gene expression along *C. dipterum* life cycle. Relative expression changes along RNA-seq samples, corresponding to different life cycle stages obtained with Mfuzz R package (E.4: 4 days post fertilization (dpf) embryo, E.6: 6 dpf embryo, E.10: 10 dpf embryo, E.14 14 dpf embryo, eN: early nymph head, mN: mid nymph head, lN: late nymph head, A: adult head).
