## Supplementary Figure 3 for "Genomic adaptations to aquatic and aerial life in mayflies and the origin of wings in insects"

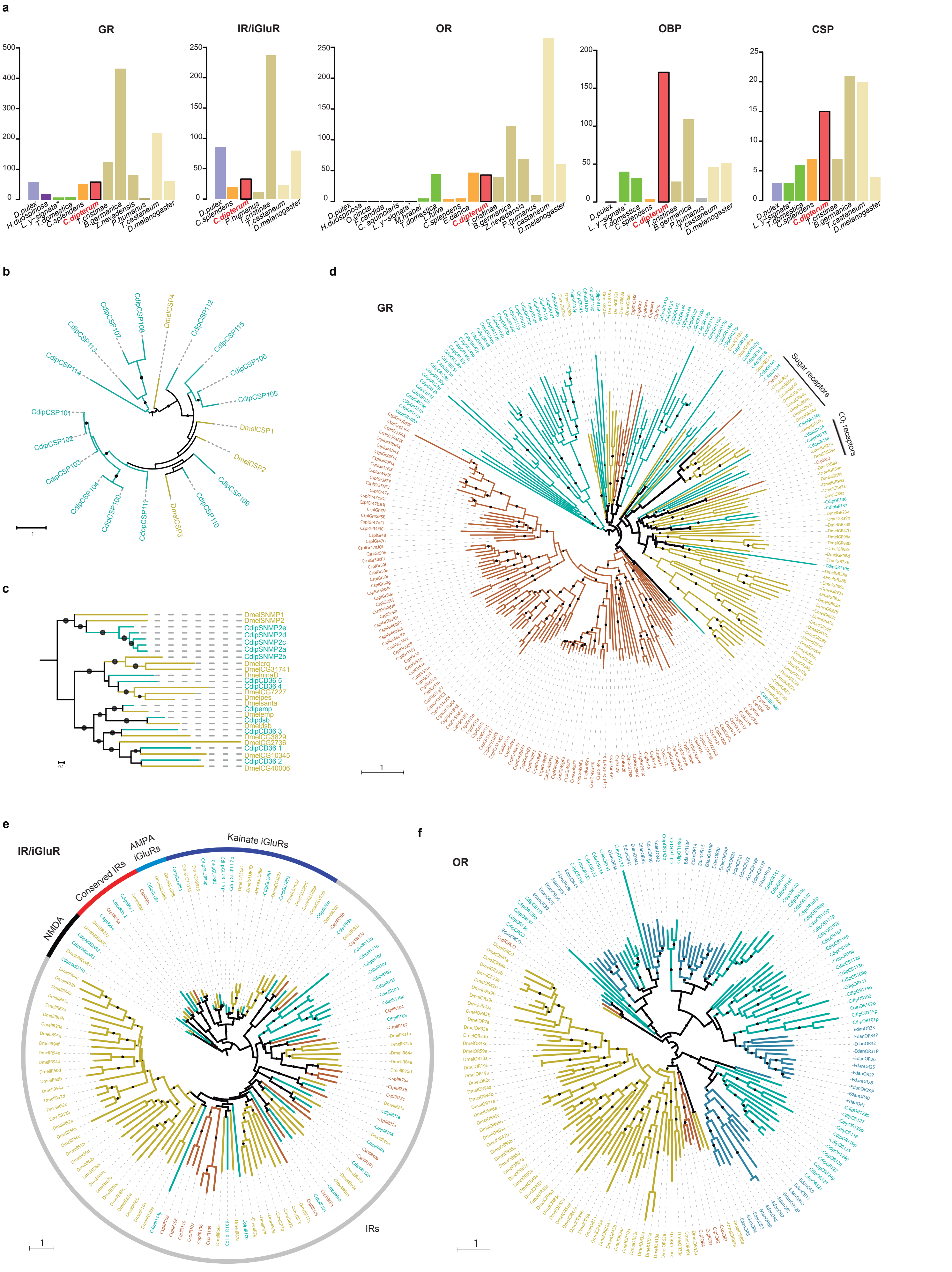


**Figure Supplementary Figure 3.** Chemosensory gene families in C. dipterum genome. a, Bar plots exhibiting the CS gene complement in several pancrustacea species (*D. pulex*, Crustacea; *H. duospinosa, O. cincta, F. candida*, Collembola; *C. aquinolaris*, Diplura; *M. hrabei, L. y-signata*, Archaeognatha; *T. domestica*, Zygentoma; *L. fulva, C. splendens*, Odonata; *E. danica, C. dipterum*, Ephemeroptera; *P. humanus*, Psocodea; *T. cristinae,* Phasmatodea; *B. germanica, Z. nevadensis*, Blattodea; *T. castaneum*, Coleoptera; *D. melanogaster*, Diptera). b, CSP gene family phylogeny. c, SNMP and CD36 gene families and phylogeny. d, GR phylogenetic tree. e, IR/iGluR phylogeny. f, OR gene complement and phylogeny. *C. dipterum* genes are shown in green, *D. melanogaster* shown in mustard, *E. danica* in blue and *C. splendens* genes are shown in brown
