## Supplementary Figure 4 for "Genomic adaptations to aquatic and aerial life in mayflies and the origin of wings in insects"

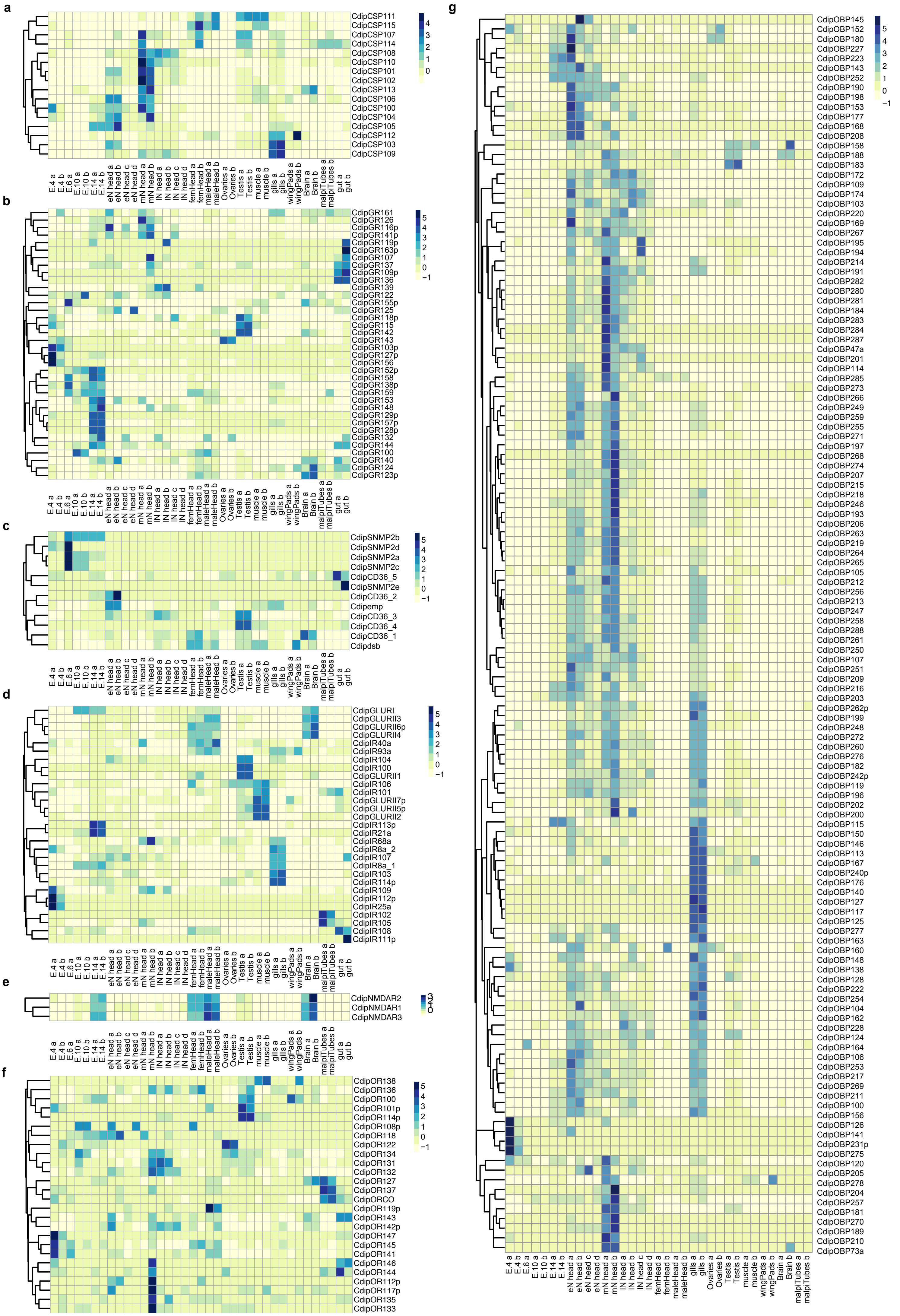


**Supplementary Figure 4.** Expression of chemosensory gene families across tissues and developmental stages. a, CSP gene family. b, GRs. c, SNMP and CD36 genes. d, IR/iGluR C. dipterum genes. e, NMDAR genes. f, OR genes. g, OBP genes. Blue corresponds to high expression; light yellow corresponds to low expression.
