## Supplementary Figure 5 for "Genomic adaptations to aquatic and aerial life in mayflies and the origin of wings in insects"

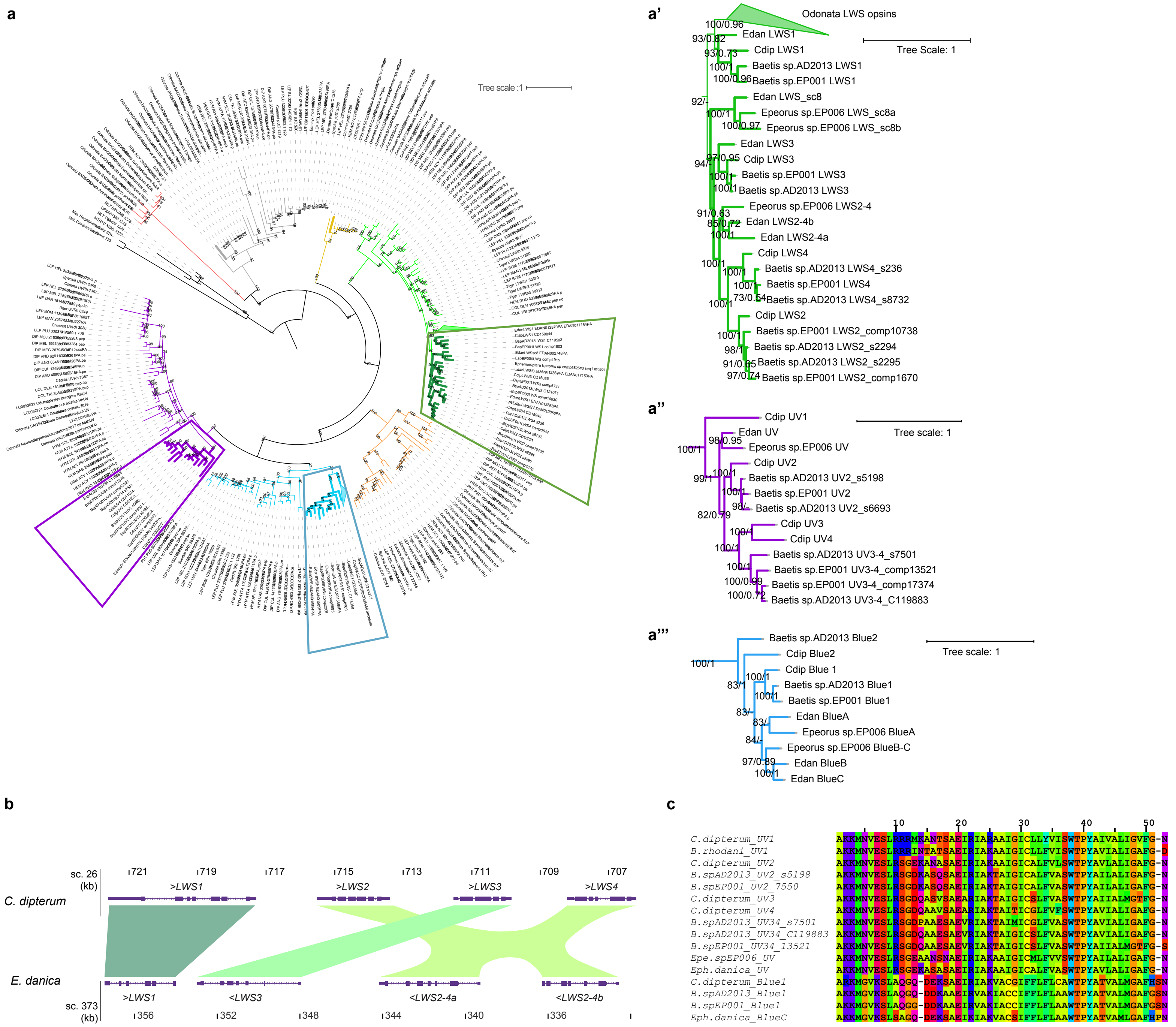


**Supplementary Figure 5.** Opsin complement in *C. dipterum*. a, ML-tree of opsin genes from different insects (DIP, Diptera; HYM, Hymenoptera; HEM, Hemiptera; COL, Coleoptera; Odonata and Ephemeroptera). Black branches represent melatonin receptor, which was chosen as outgroup; in red, RGR genes; in grey, pteropsins; arthropsin in mustard; in green, LWS opsin; in orange Rh7; in blue, Blue sensitive opsins and in violet, UV-sensitive opsins. a', Detail of Ephemeroptera LWS Opsin phylogeny. Numbers at each node represent boostrap and posterior probabilities values, respectively. a'', UV-sensitive Opsin. a''', Blue sensitive Opsins in Ephemeroptera. b, Ancestral expansion of LWS opsins in Ephemeroptera is located in a genomic cluster. Upper track shows *C. dipterum* genomic region and lower track shows *E. danica* region. c, Alignment of exon 6 of *C. dipterum* UV-Ops1 with its orthologous region from *B. rhodani* and homologous regions from other mayflies
