## Supplementary Figure 6 for "Genomic adaptations to aquatic and aerial life in mayflies and the origin of wings in insects"

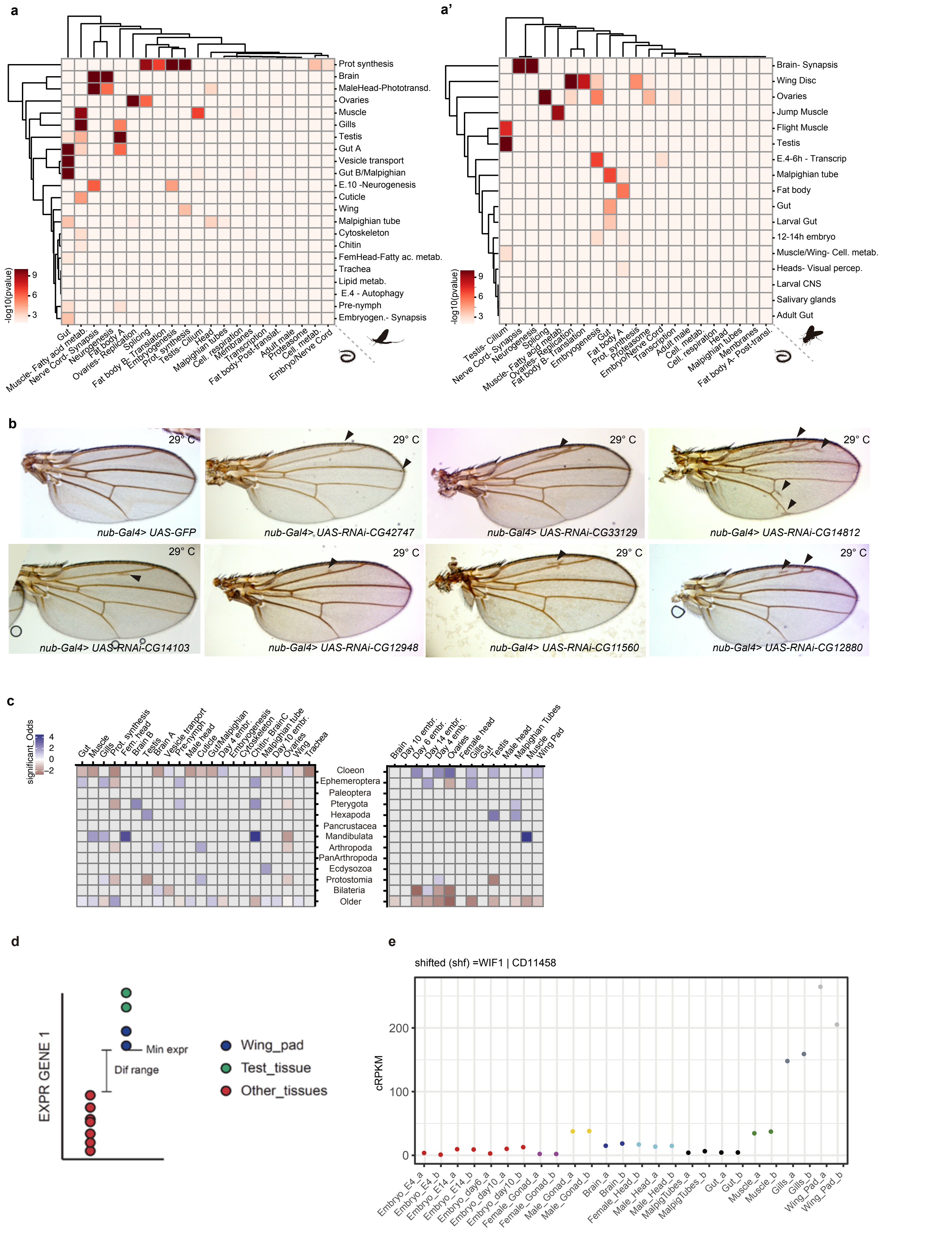


**Supplementary Figure 6.**Transcriptomic conservation of wings and other insect tissues. a, Orthologous gene overlap between modules from *C. dipterum* (vertical) and *S. maritima* (horizontal) obtained by weighted correlation network analysis (WGCNA) shown by levels of raw statistical significance. a’, Overlap of orthologues between modules from *D. melanogaster* (vertical) and *S. maritima* (horizontal) obtained by weighted correlation network analysis (WGCNA) shown by levels of raw statistical significance. b, Wing phenotypes observed when knocking down orthologous genes (*CG42747, CG33129, CG14812, CG14103, CG12948, CG11560 and CG12880*) shared between *C. dipterum* and *D. melanogaster* wing and wing disc modules. Control wings: *nub-Gal4; UAS-GFP.* Extra vein territories are highlighted by black arrowheads. c, Heat maps showing enrichment test on gene age/phylostrata in WGCNA modules (left heat map) and tissue-specific RNA datasets (right heat map) relative to the background proportion of ages of the species of interest (p-value < 0.01). d, Scheme showing the methodology used to assign transcriptomic similarities between tissues. e, *CD11458*, which *D. melanogaster* orthologue is *shifted*, is highly expressed in wings and gills, in comparison to the rest of the tissue-specific RNA samples
