## Supplementary Data 1 for "Genomic adaptations to aquatic and aerial life in mayflies and the origin of wings in insects"

#### WGCNA module annotation

1. *C. dipterum*, *D. melanogaster* and *S. maritima* module annotations
Module colour and annotated names……………………………………………………page 2

2. *C. dipterum* module RNA-seq expression and GO terms………………………………..page 3

3. *D. melanogaster* module RNA-seq expression and GO terms……………………….page 14

4. *S. maritima* module RNA-seq expression and GO terms……………………………..page 23

| *C. dipterum* WGCNA modules annotation | | *D. melanogaster* WGCNA modules annotation | | *S.maritima* WGCNA modules annotation | |
|---|---|---|---|---|---|
| Module Colour | Module annotated name | Module Colour | Module annotated name | Module Colour | Module annotated name |
| grey60 | Vesicle transport | darkgrey | Gut | lightcyan | Head |
| greenyellow | Brain | darkturquoise | Larval CNS | saddlebrown | Adult male |
| sienna3 | Cytoskeleton | salmon | Jump Muscle | greenyellow | Neurogenesis |
| lightgreen | Pre-nymph | red | Salivary glands | red | Fat body |
| yellow | Wing | blue | Brain- Synapsis | darkgreen | membrane |
| steelblue | Malpighian tube | greenyellow | Wing Disc | skyblue | Embryo/Nerve cord |
| brown | Gills | cyan | 12-14 hpf embryo | orange | Protein synthesis |
| orange | Cuticle | black | Flight muscle | black | Nerve cord- Synapsis |
| blue | Muscle | magenta | 4-6 h embryo- Transcription | midnightblue | Splicing |
| skyblue | Chitin | purple | Adult heads- visual perception | blue | Gut |
| yellowgreen | Trachea | green | Ovaries | grey60 | Malpighian tubes |
| black | Gut | lightcyan | Fat body | purple | Fat body B- Translation |
| paleturquoise | Gut/Malpighian | white | Muscle- Cell metabolism | white | Proteasome |
| cyan | Protein synthesis | brown | Testis | lightgreen | Cellular respiration |
| tan | 10 dpf embryo- Neurogenesis | pink | Malpighian tubule | magenta | Ovaries-Replication |
| darkolivegreen | Brain B- Lipid metabolism | skyblue | Adult gut | lightyellow | Fatty acid metabolism |
| magenta | Male Head- Phototransduction | grey60 | Larval gut | royalblue | Cellular metabolism |
| turquoise | Ovaries | | | darkorange | Fat body- Post-translation modification |
| darkmagenta | Female Head- Fatty acid metabolism | | | brown | Embryogenesis |
| red | 4 dpf embryo- Autophagy | | | turquoise | Testis-Cilium |
| saddlebrown | Embryogenesis- Synapsis | | | darkgrey | Transcription |
| green | Testis | | | | |

### *C. dipterum* modules

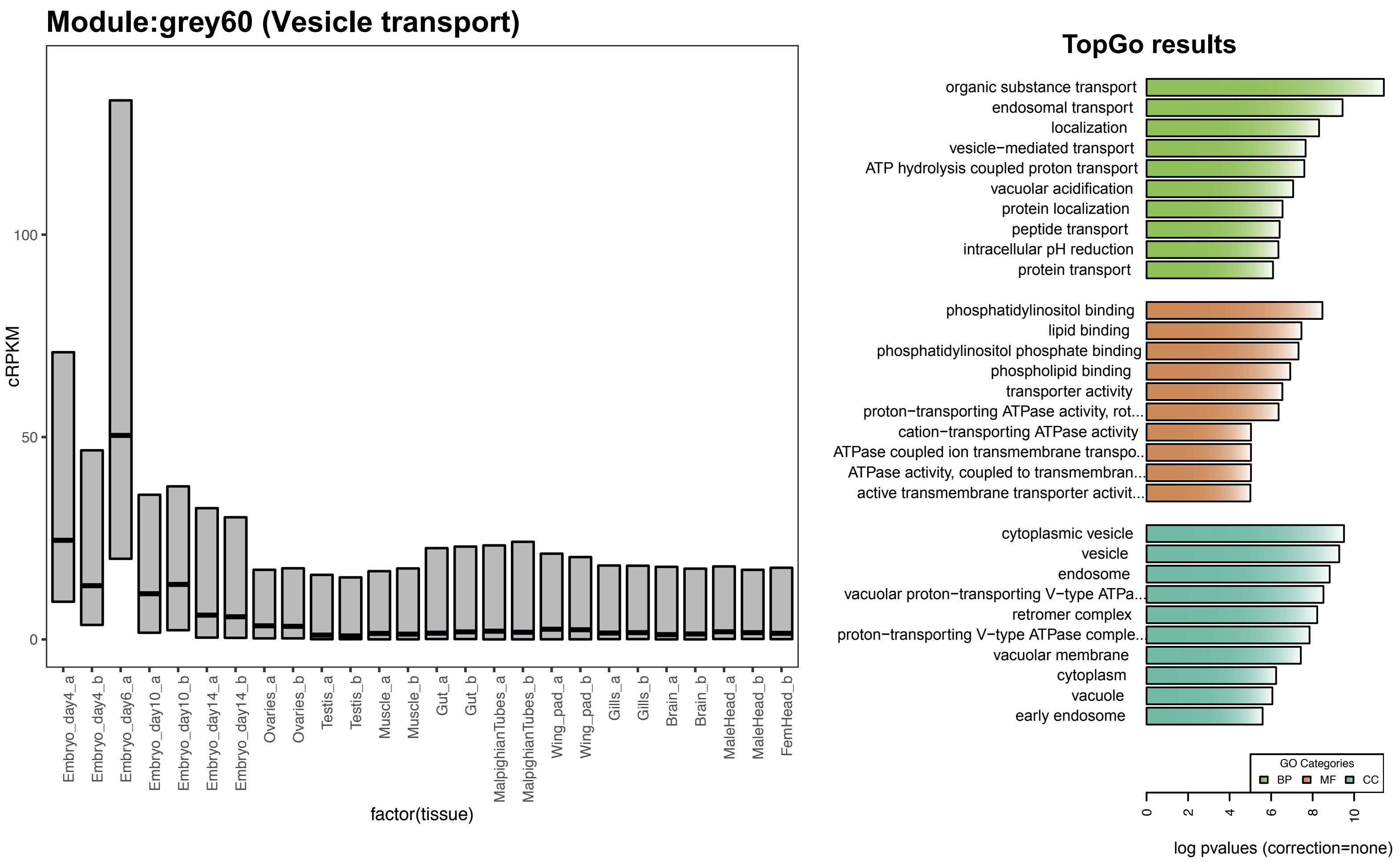


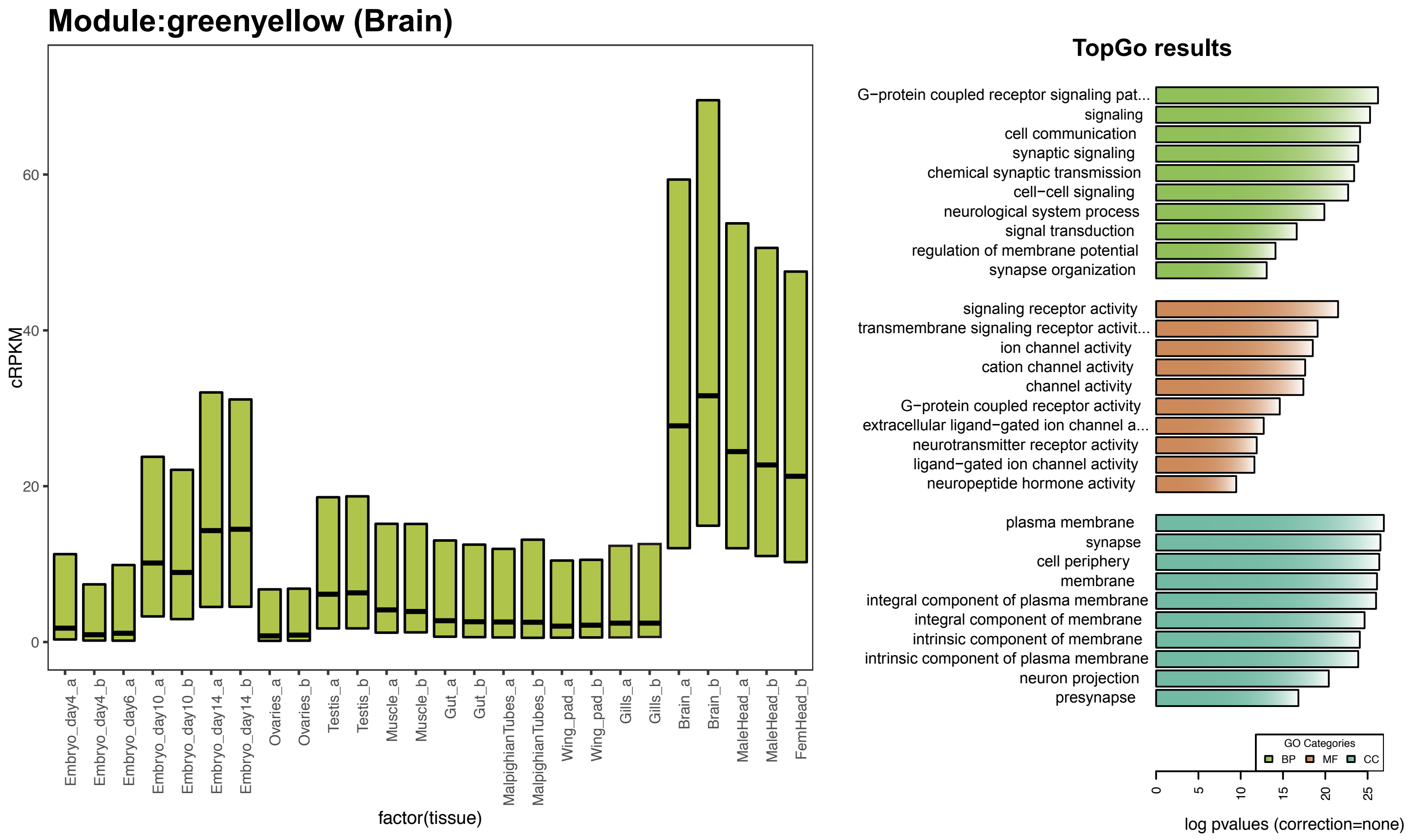

### *C. dipterum* modules

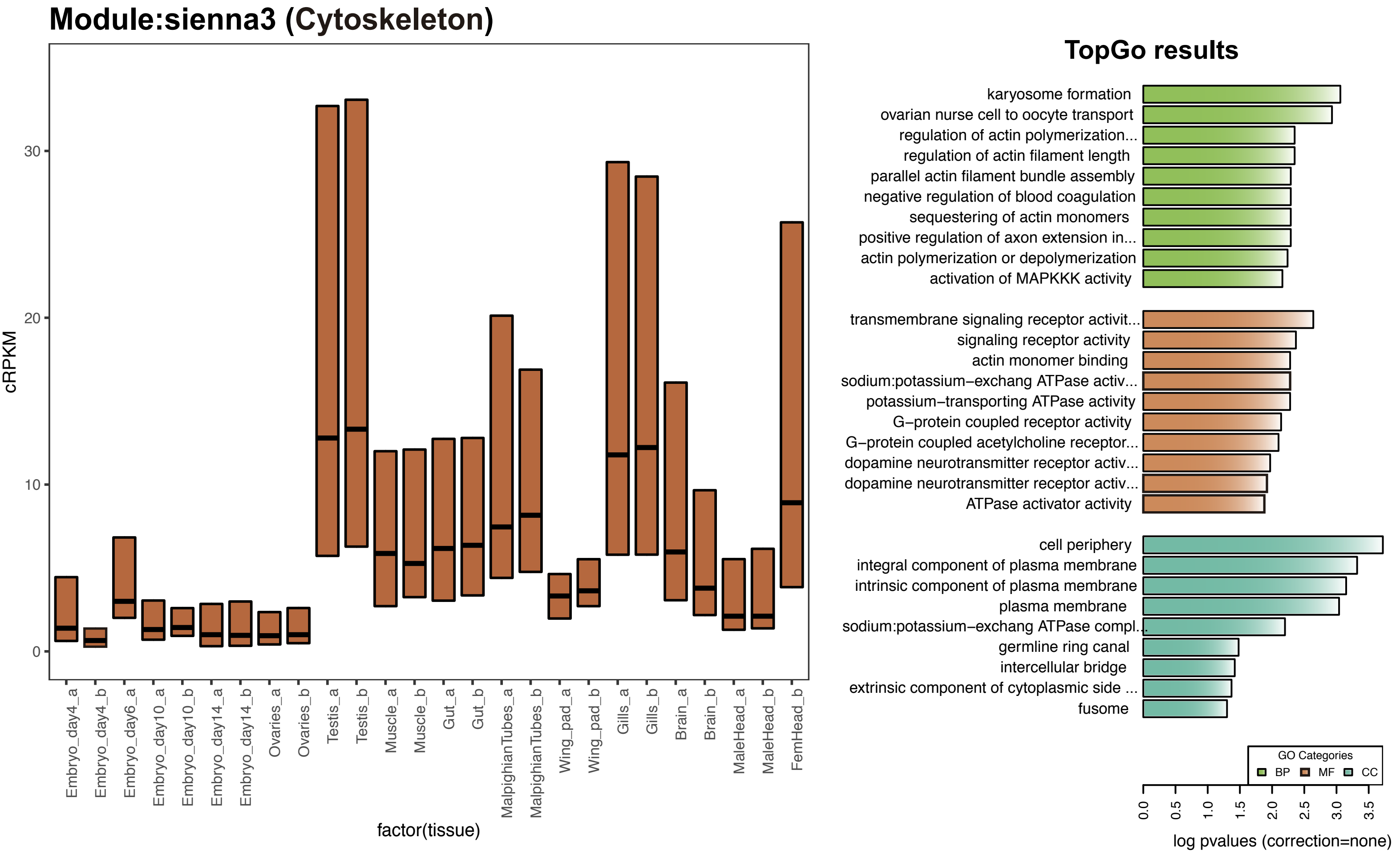


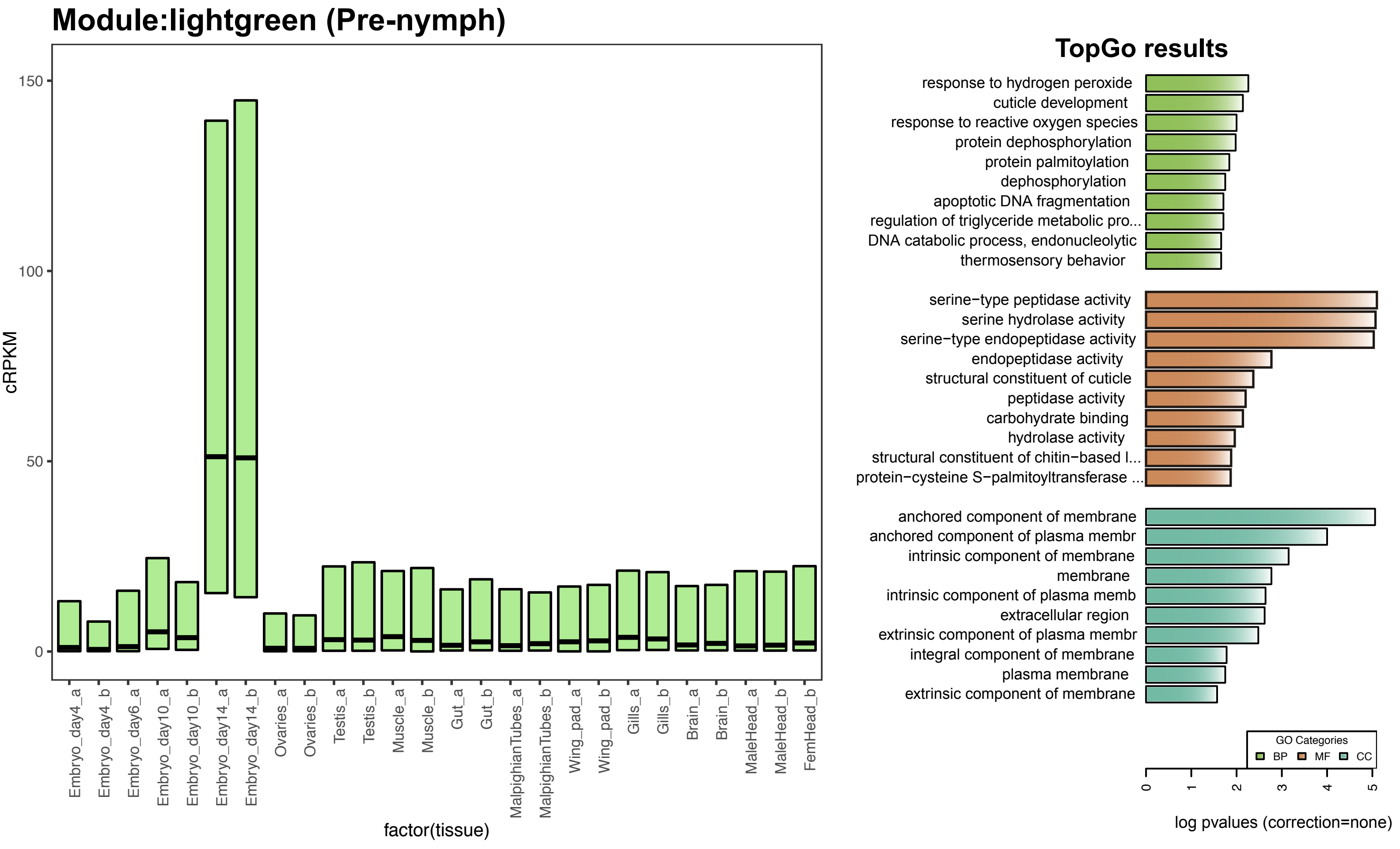

### *C. dipterum* modules

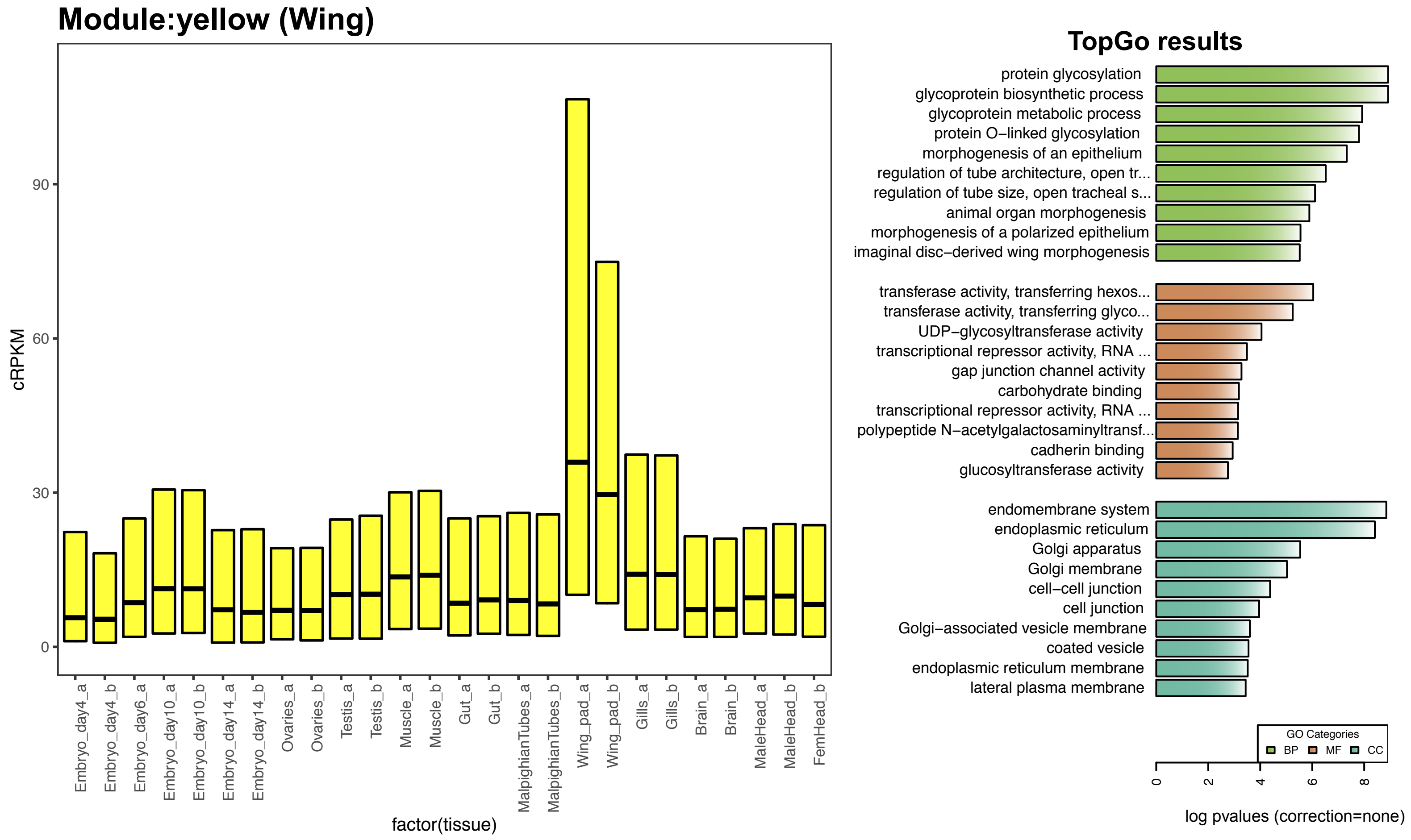


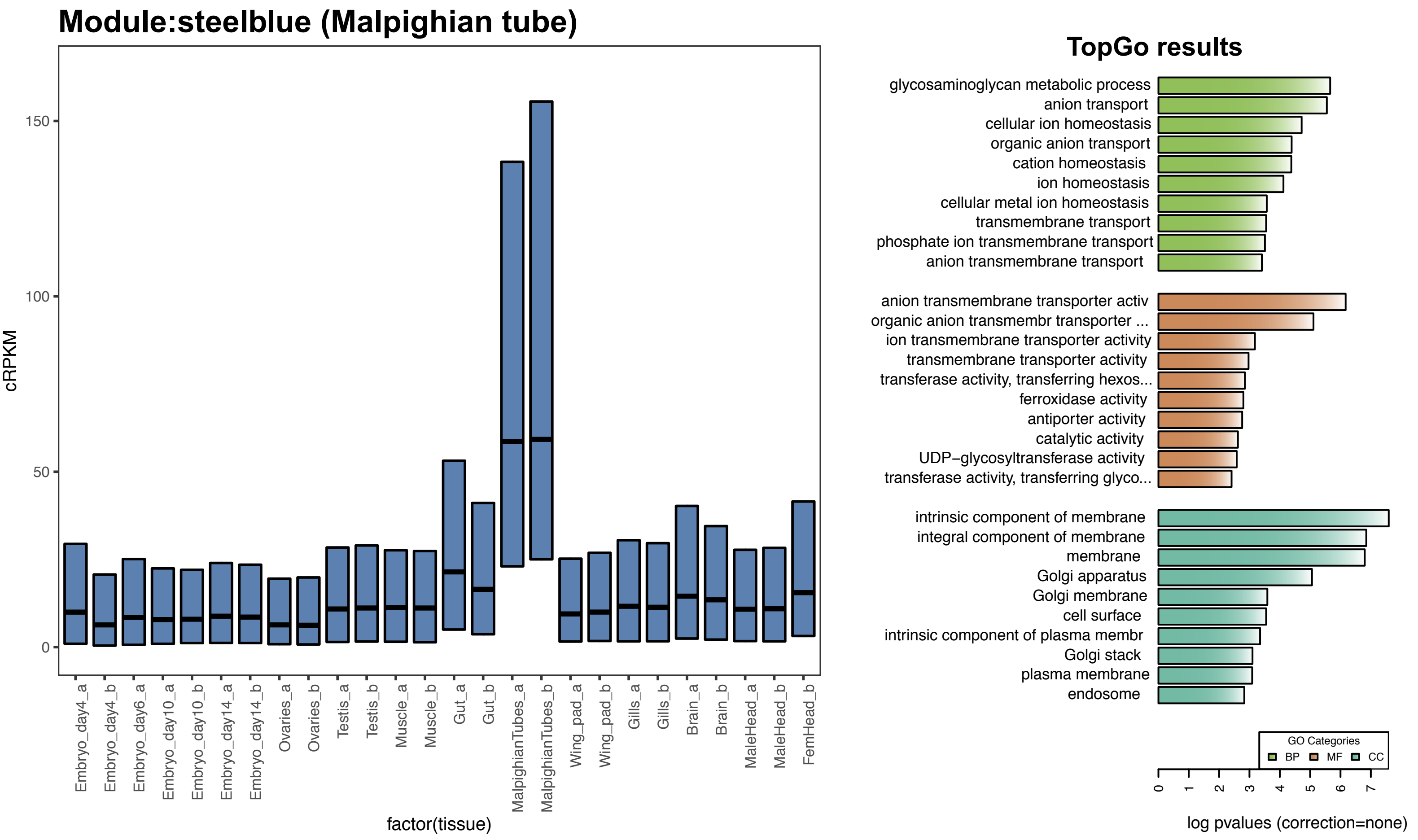

### *C. dipterum* modules

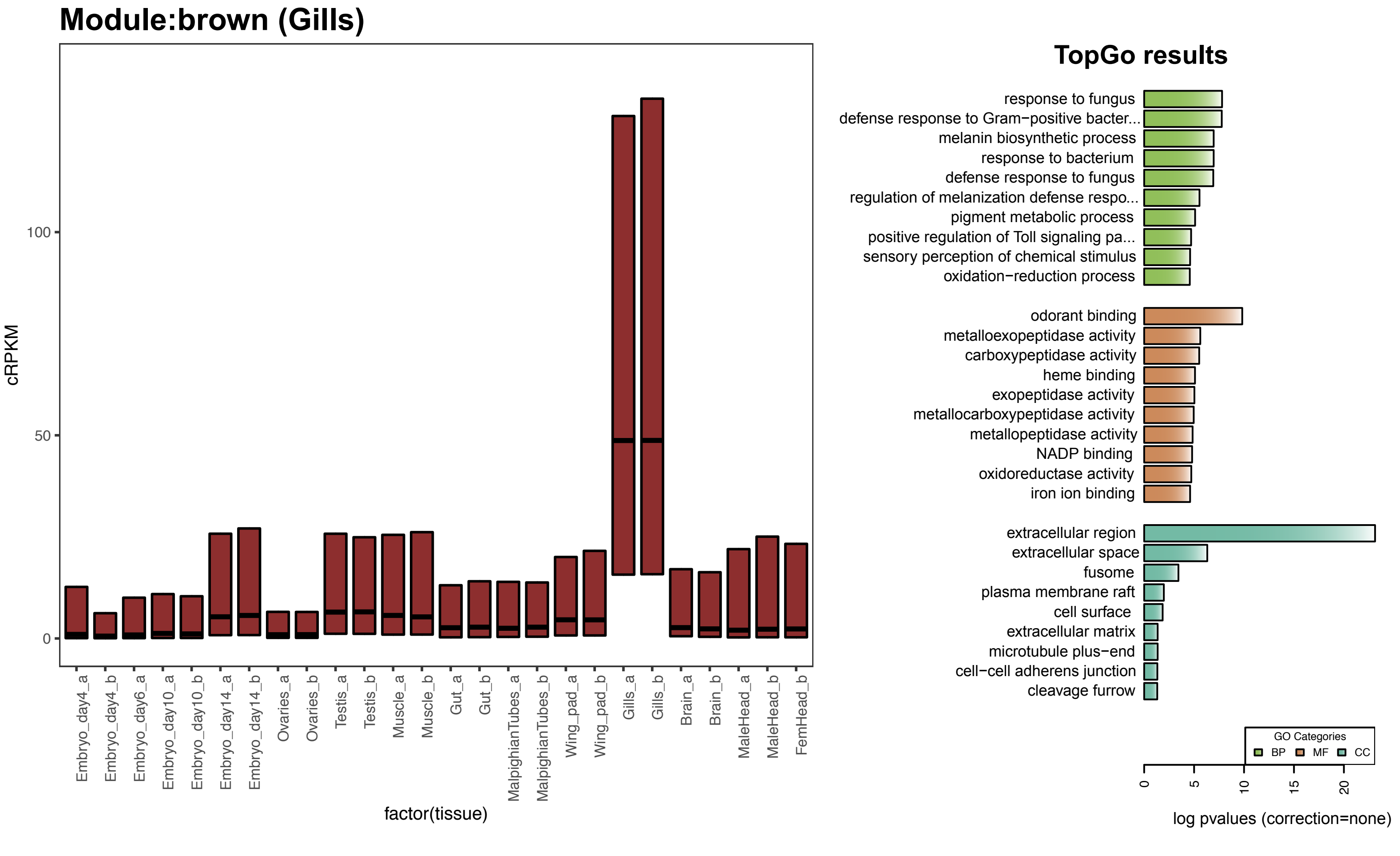


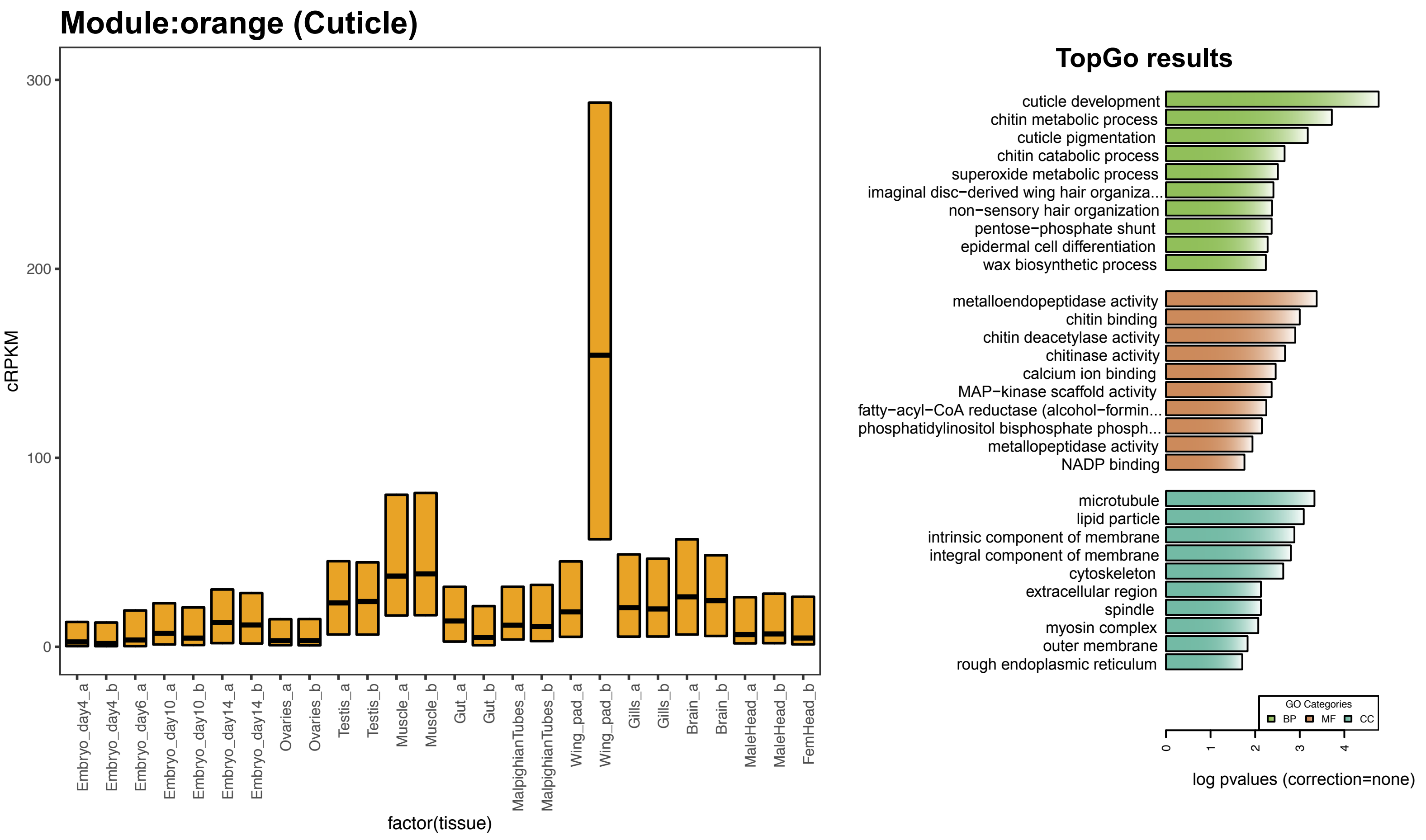

### *C. dipterum* modules

#### Module:blue (Muscle)

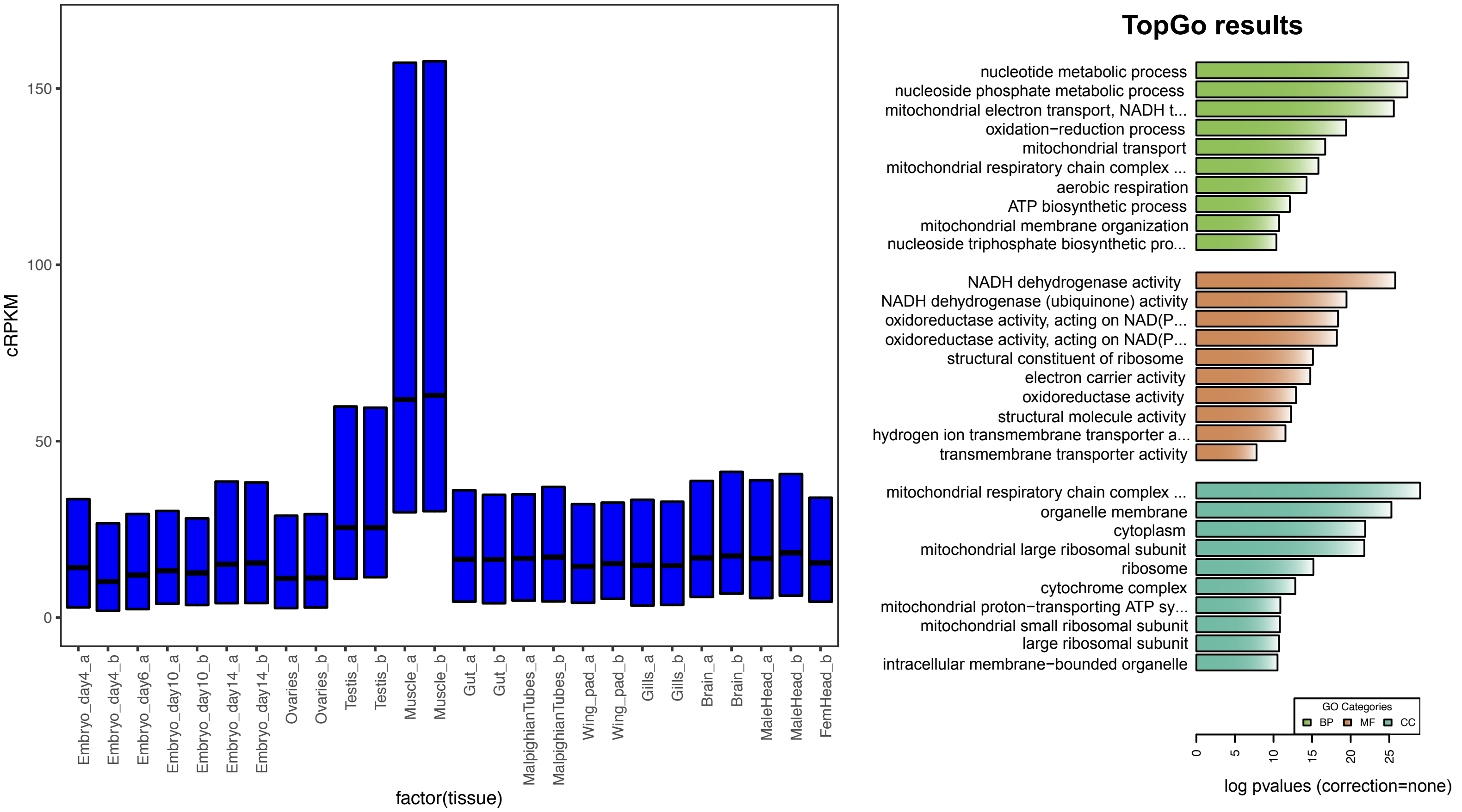


#### Module:skyblue (Chitin)

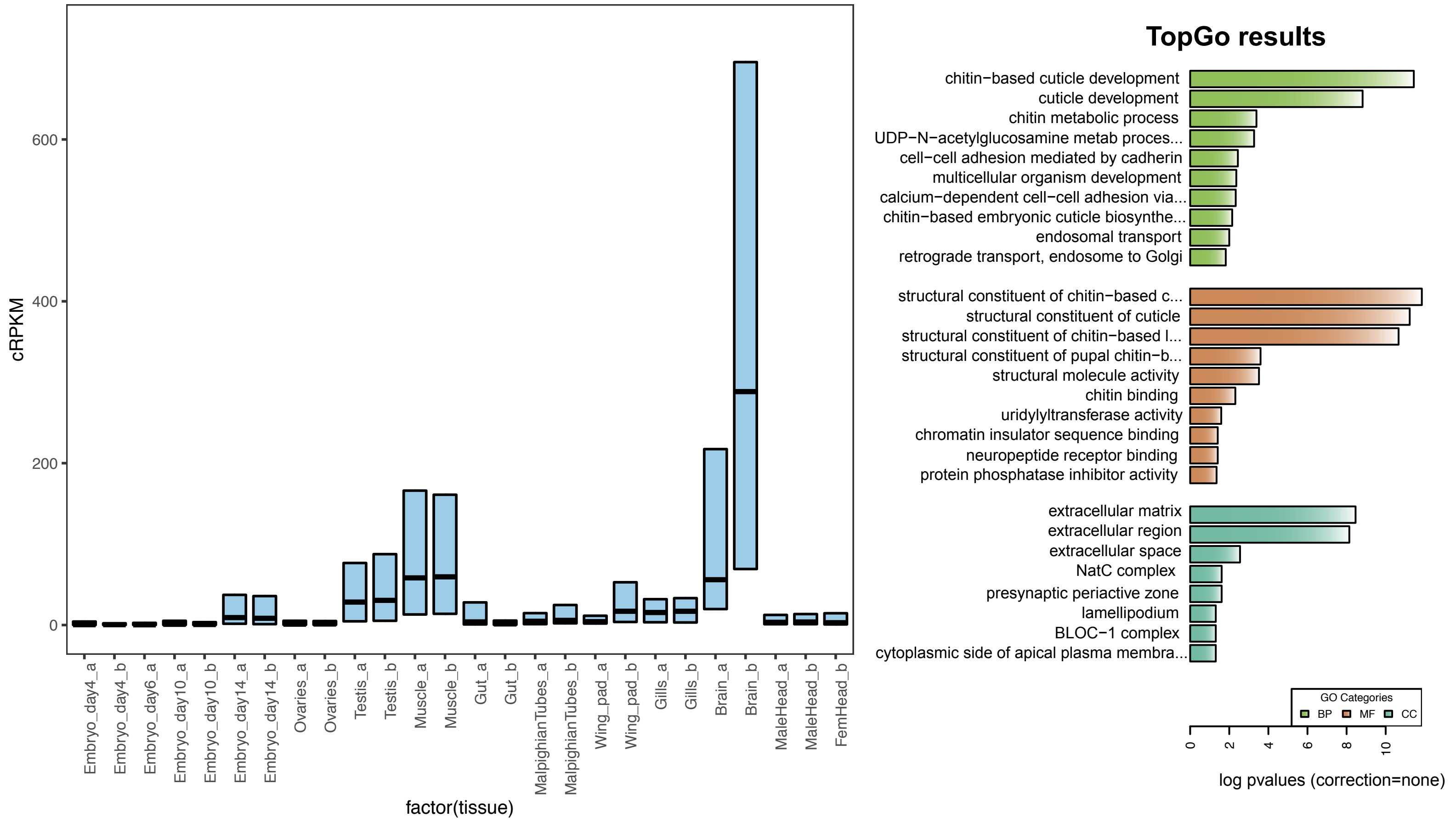

### *C. dipterum* modules

#### Module:yellowgreen (Trachea)

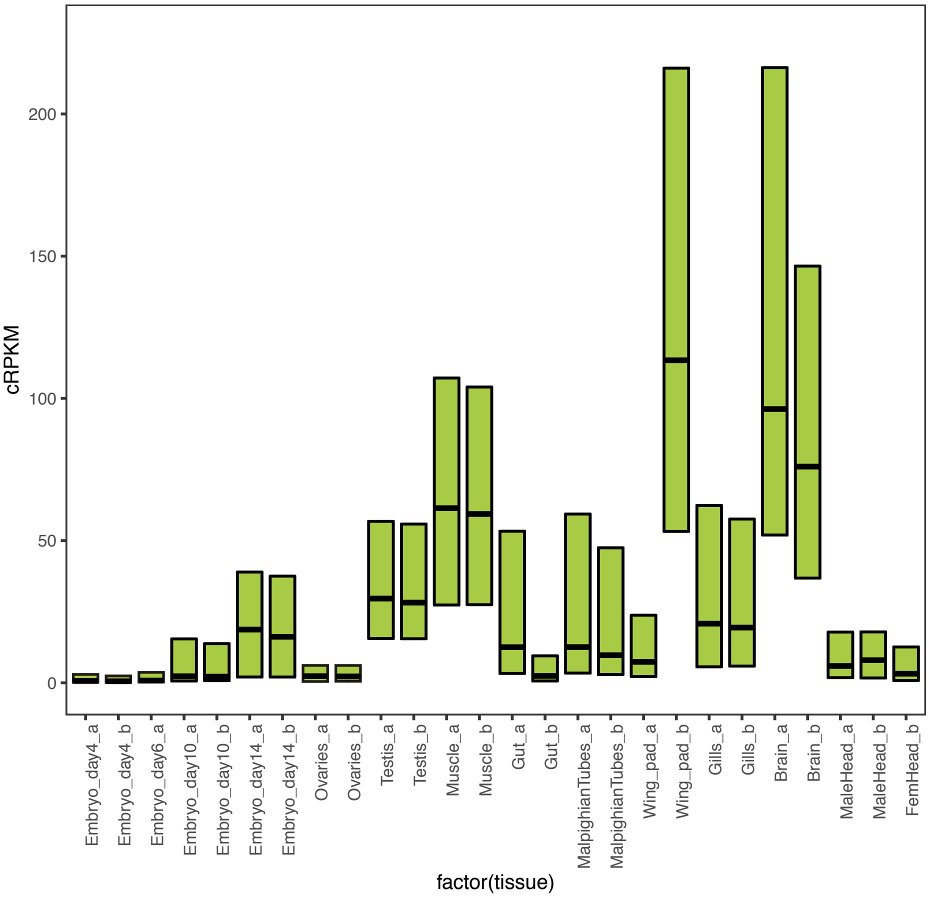


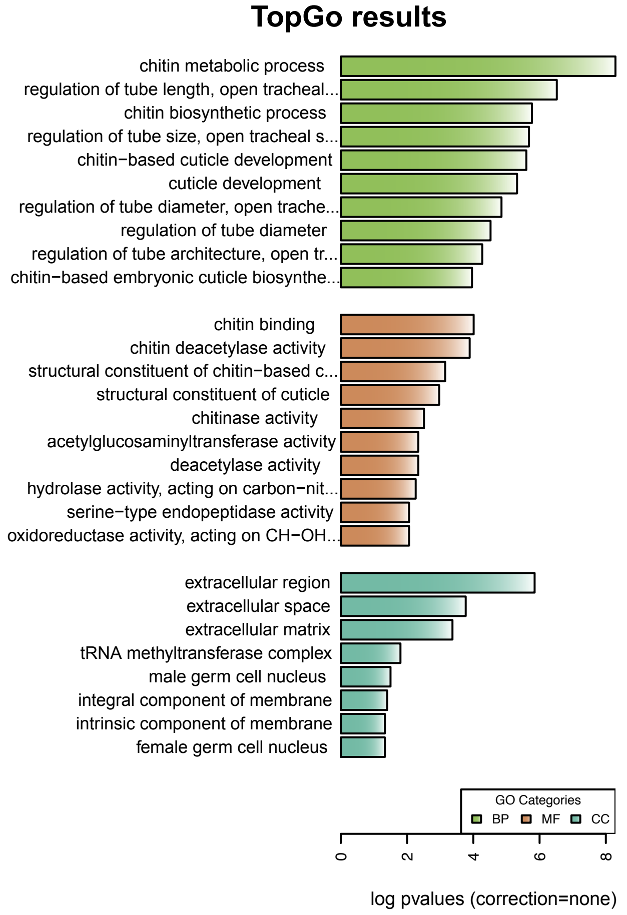


#### Module:black (Gut)

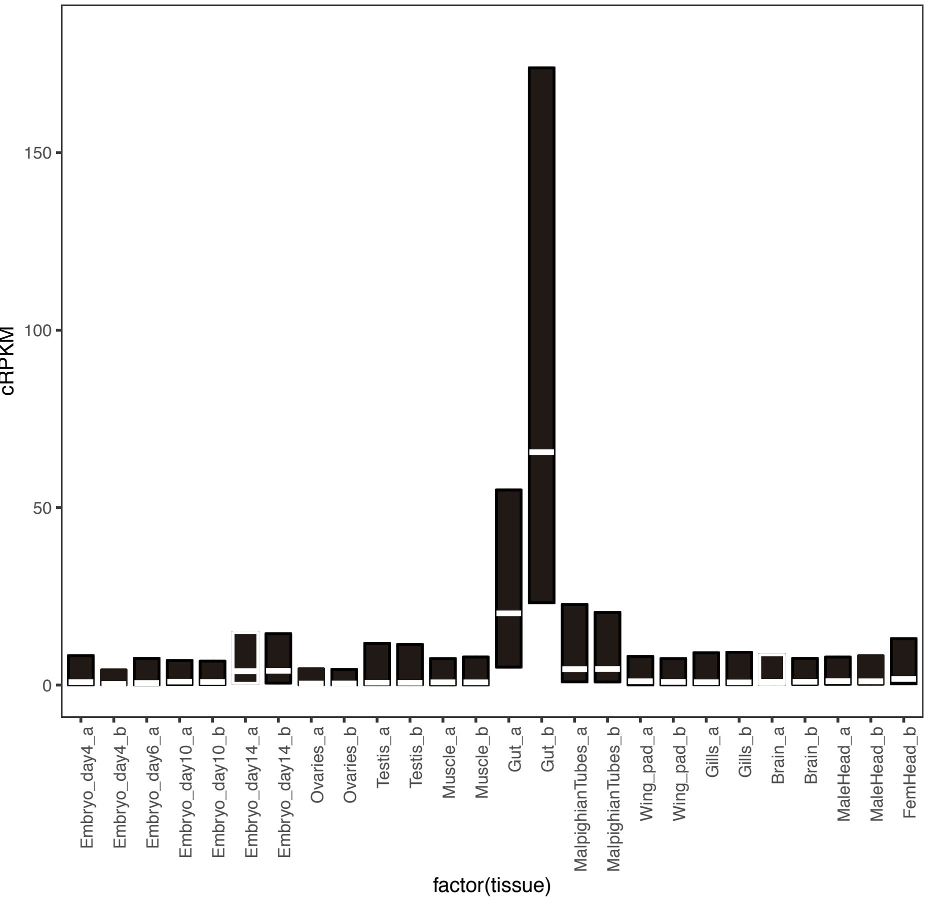


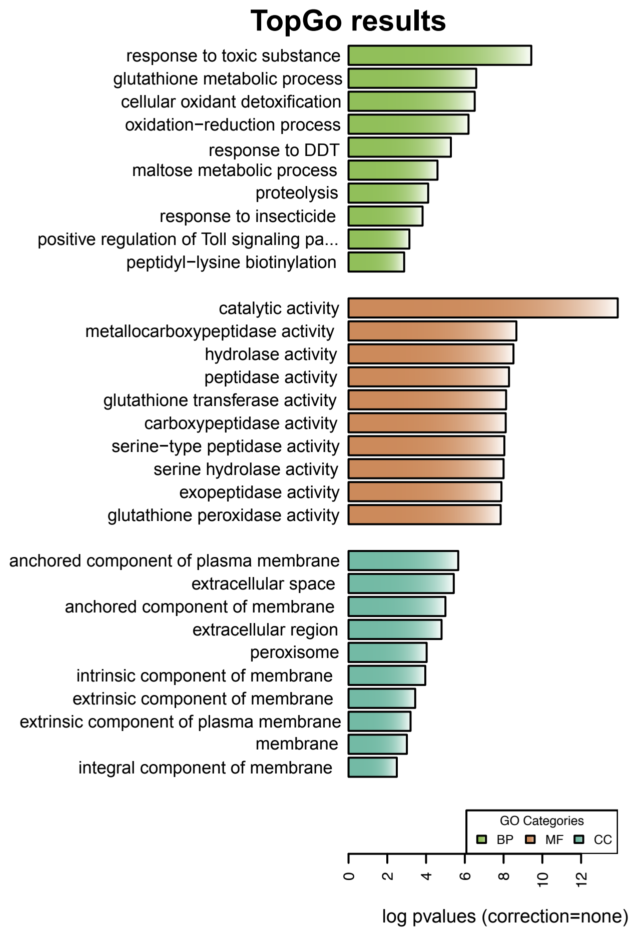

### *C. dipterum* modules

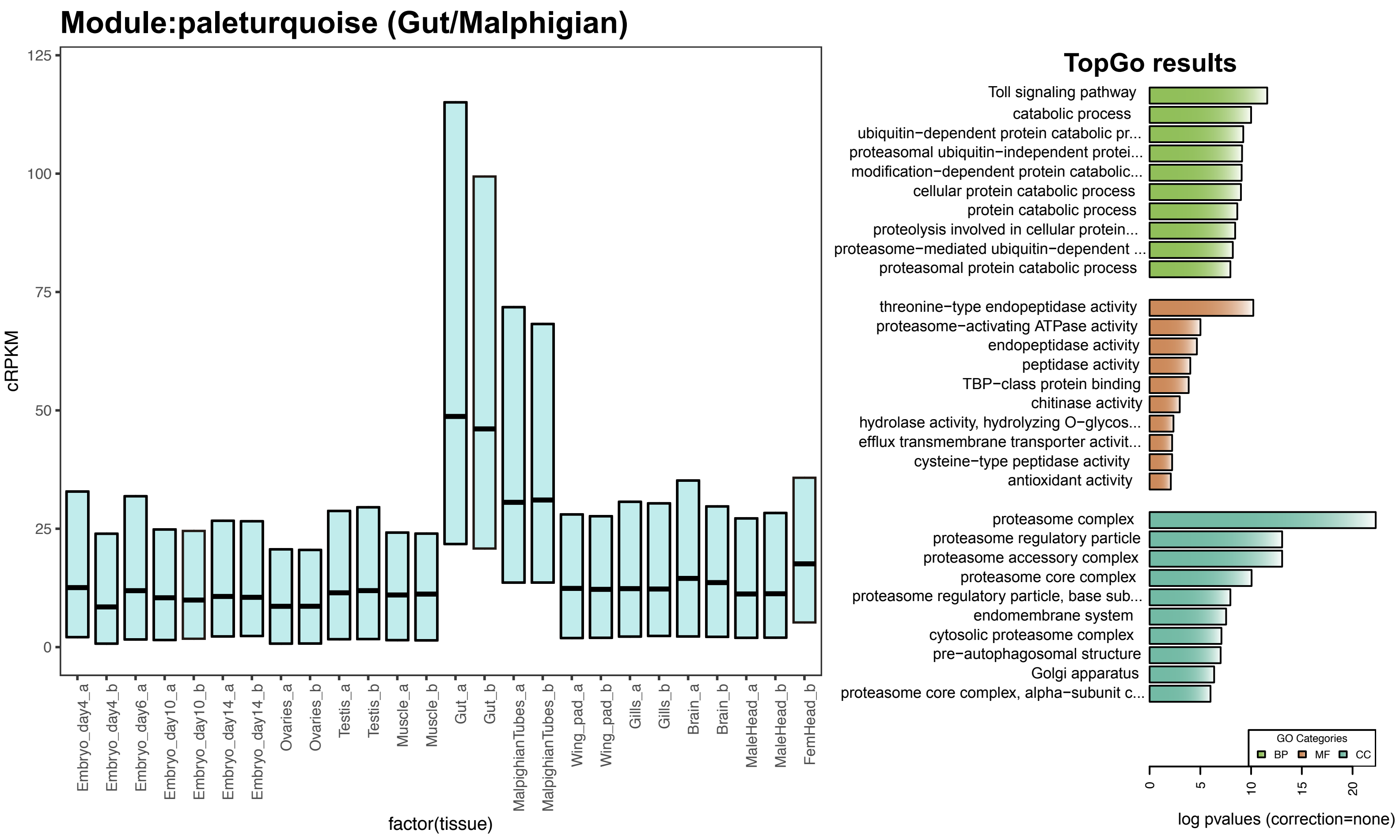


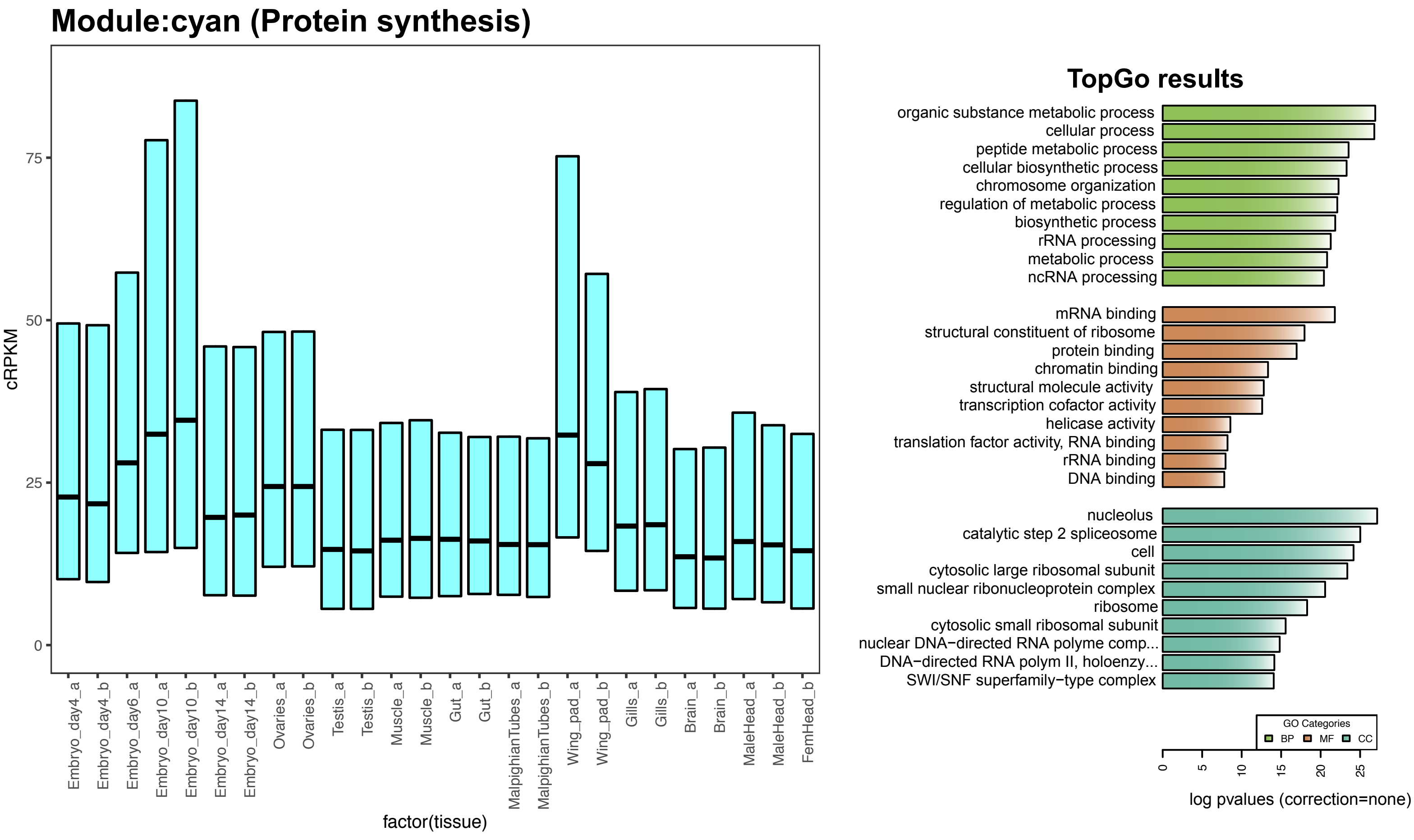

### *C. dipterum* modules

#### Module:tan (Day 10 embryo- Neurogenesis)

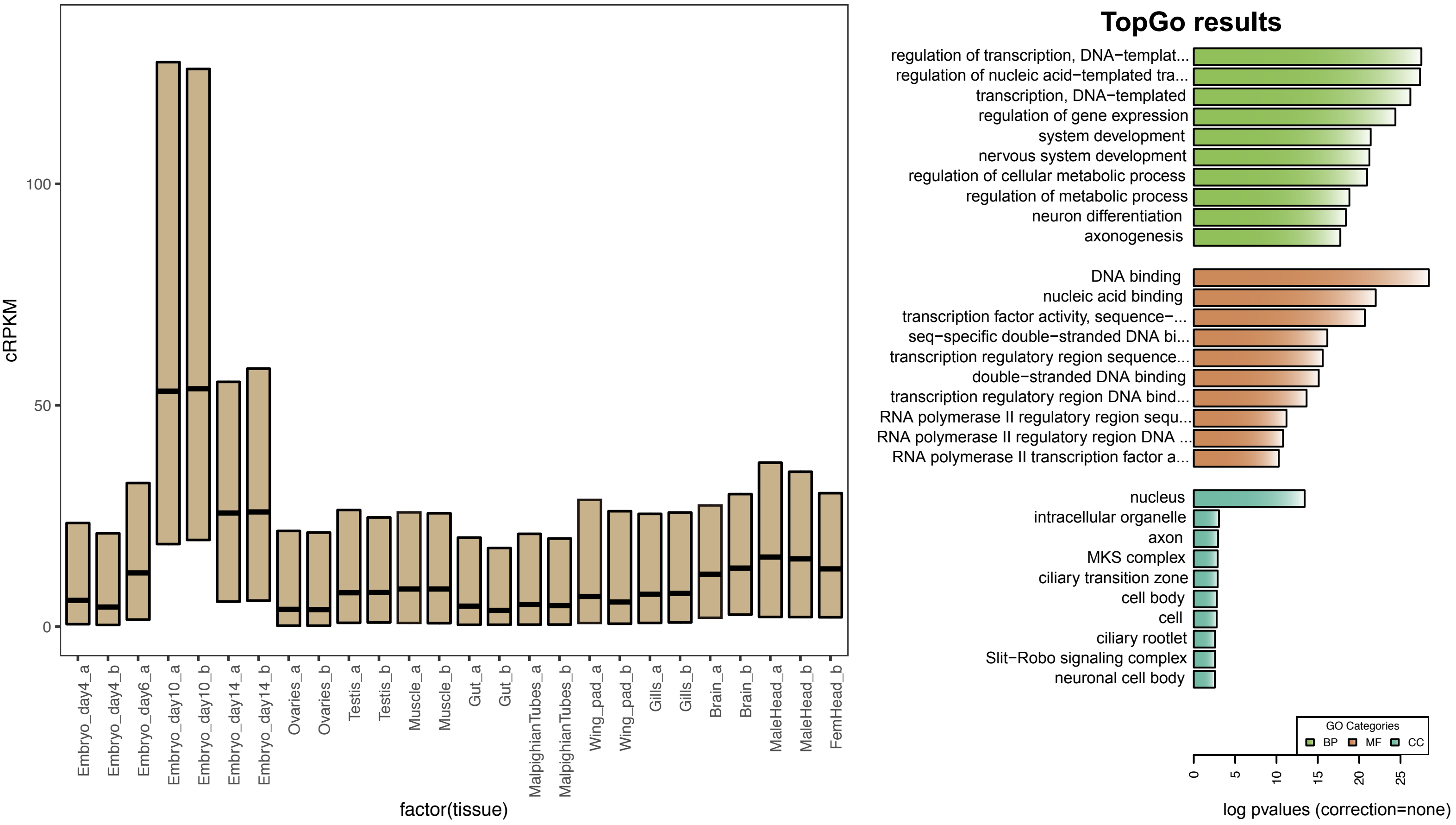


#### Module:darkolivegreen (Brain B- Lipid metabolism)

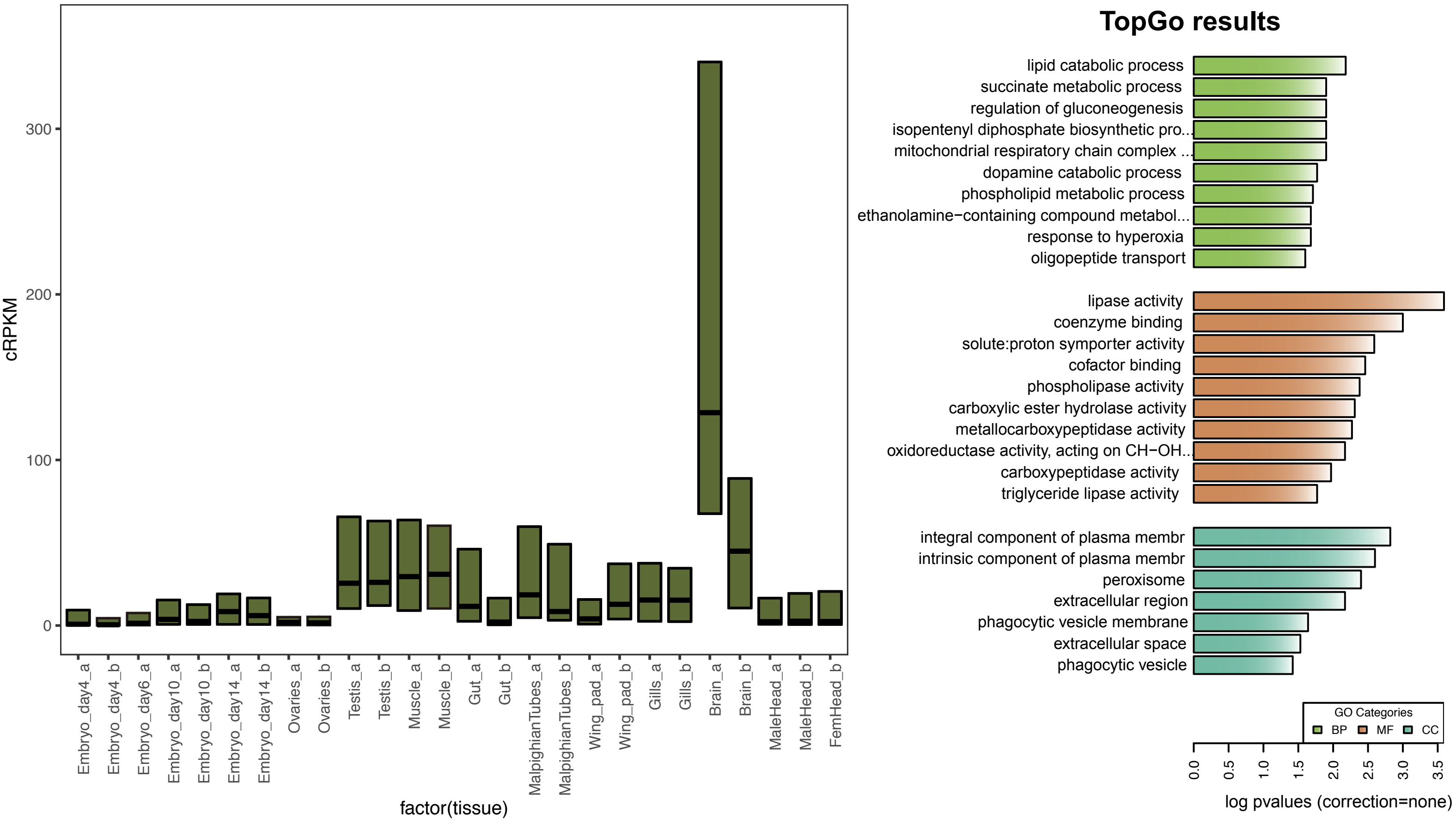

### *C. dipterum* modules

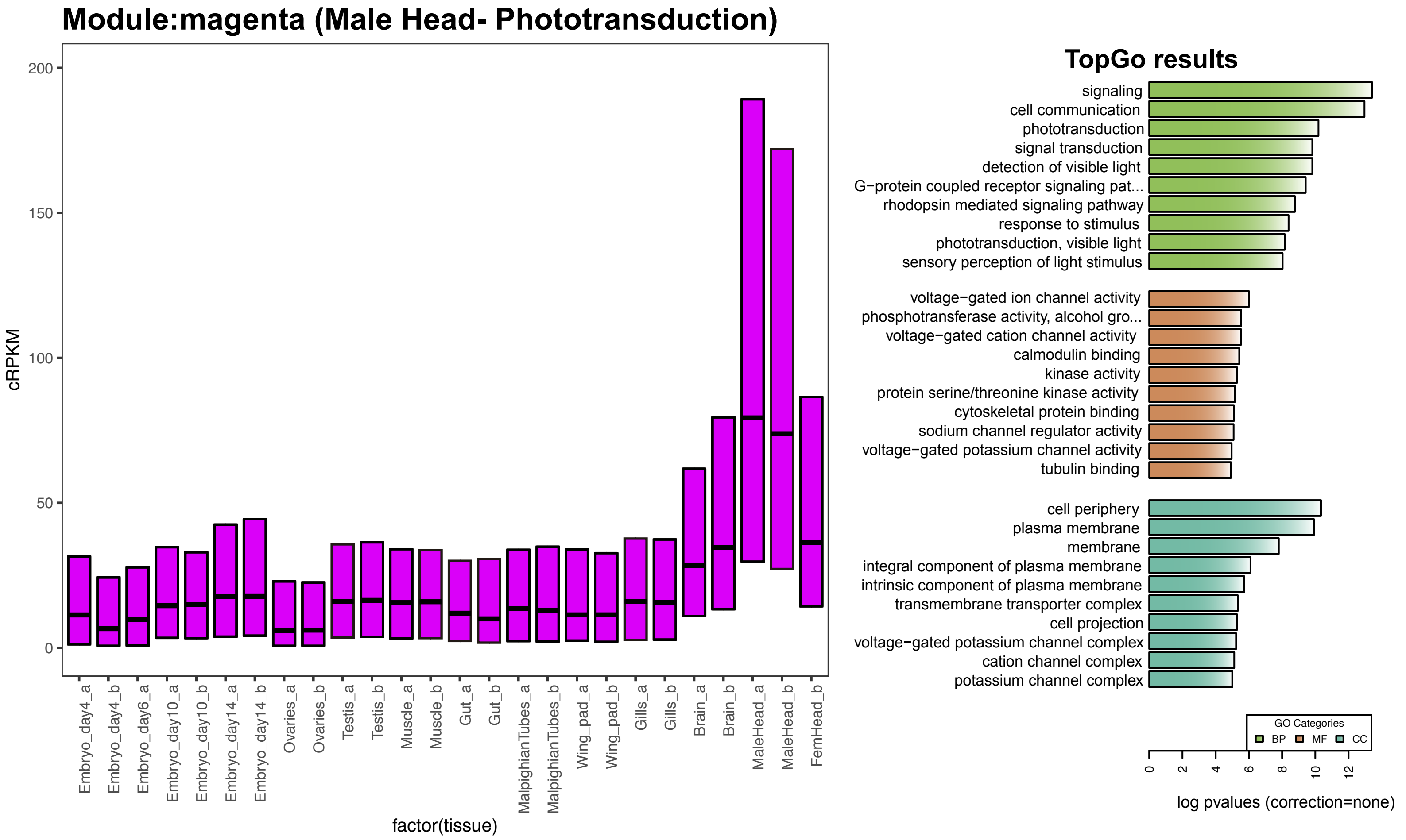


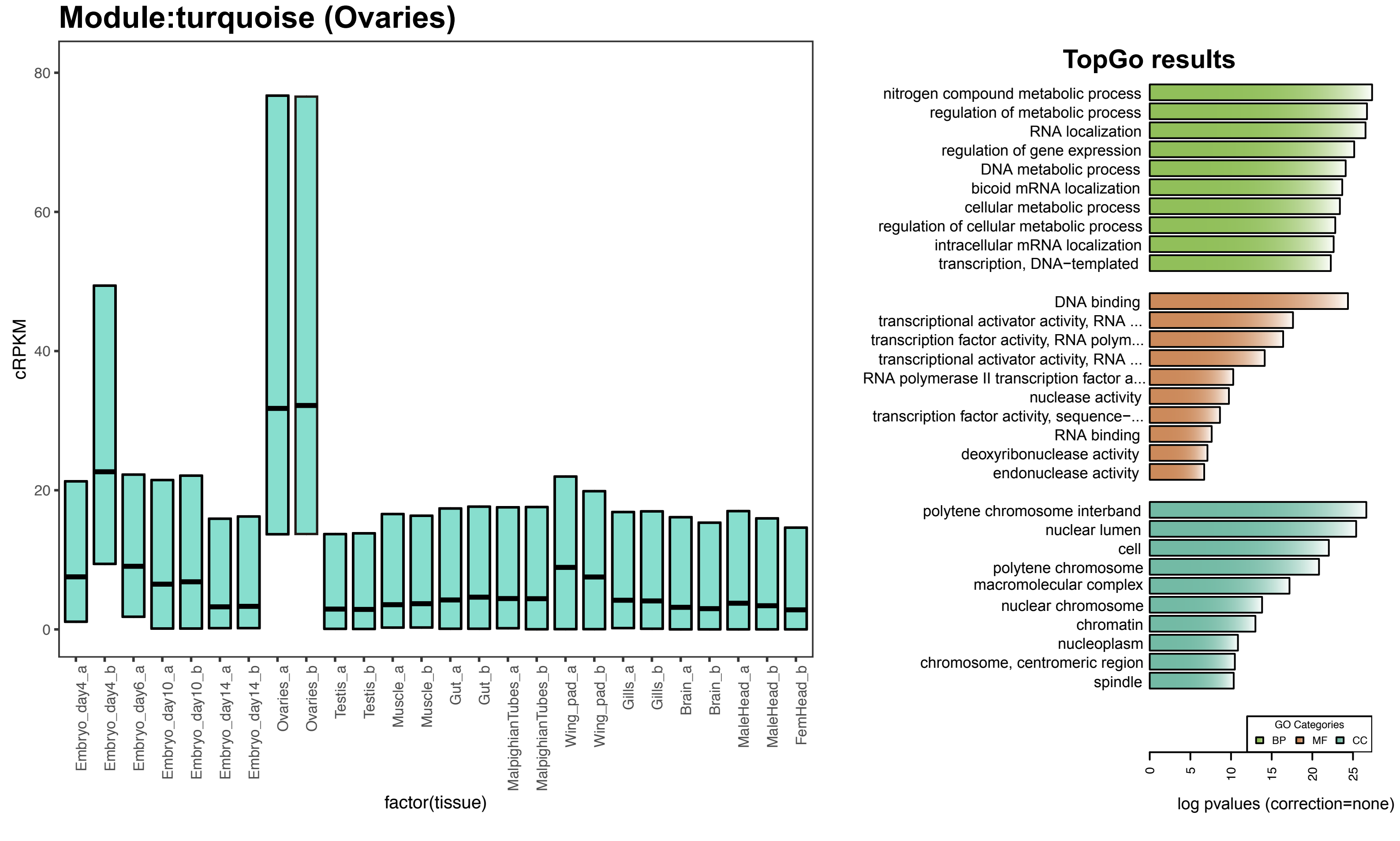

### *C. dipterum* modules

#### Module:darkmagenta (Female Head- Fatty acid metab)

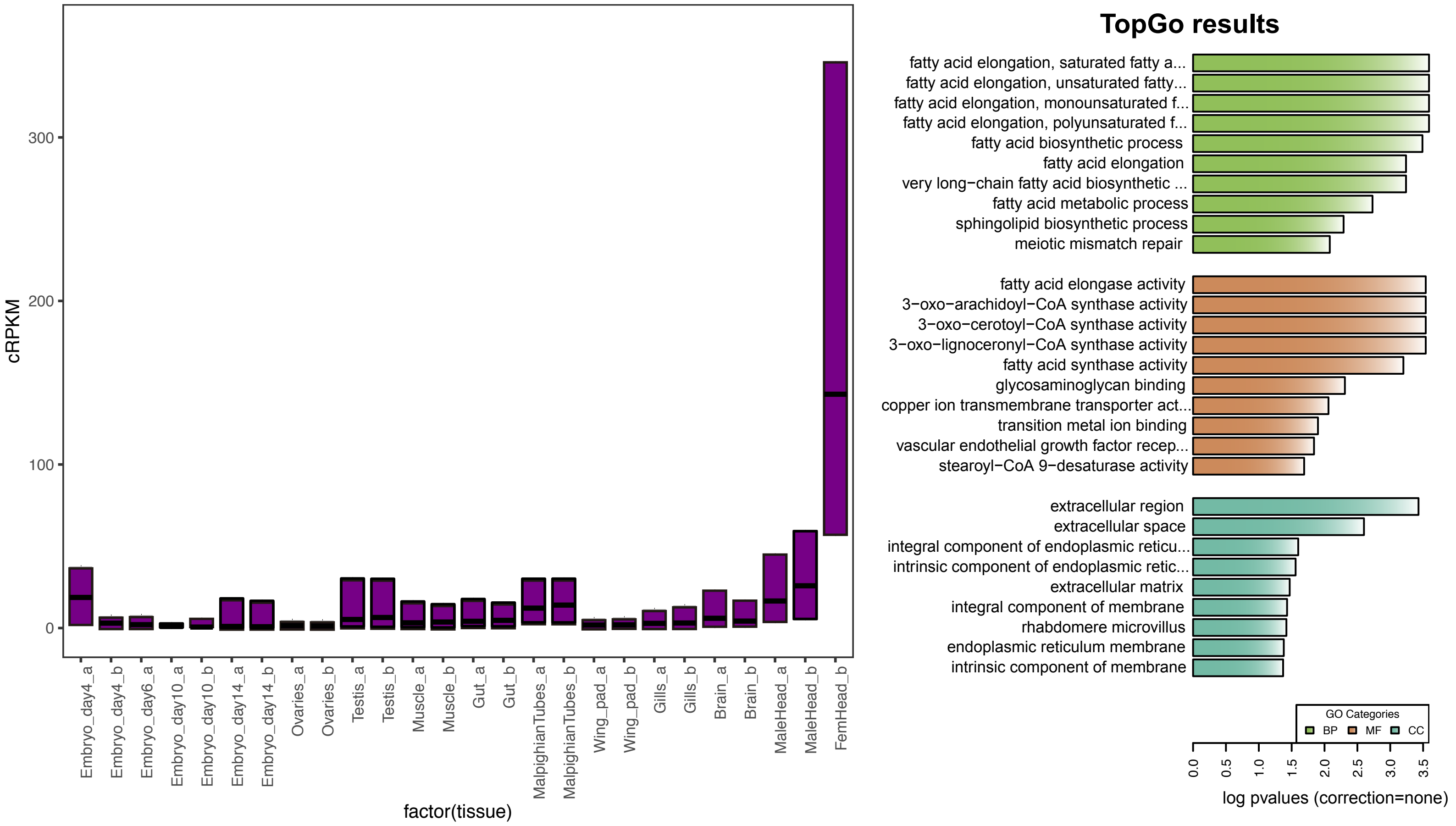


#### Module:red (Embryo day 4- Autophagy)

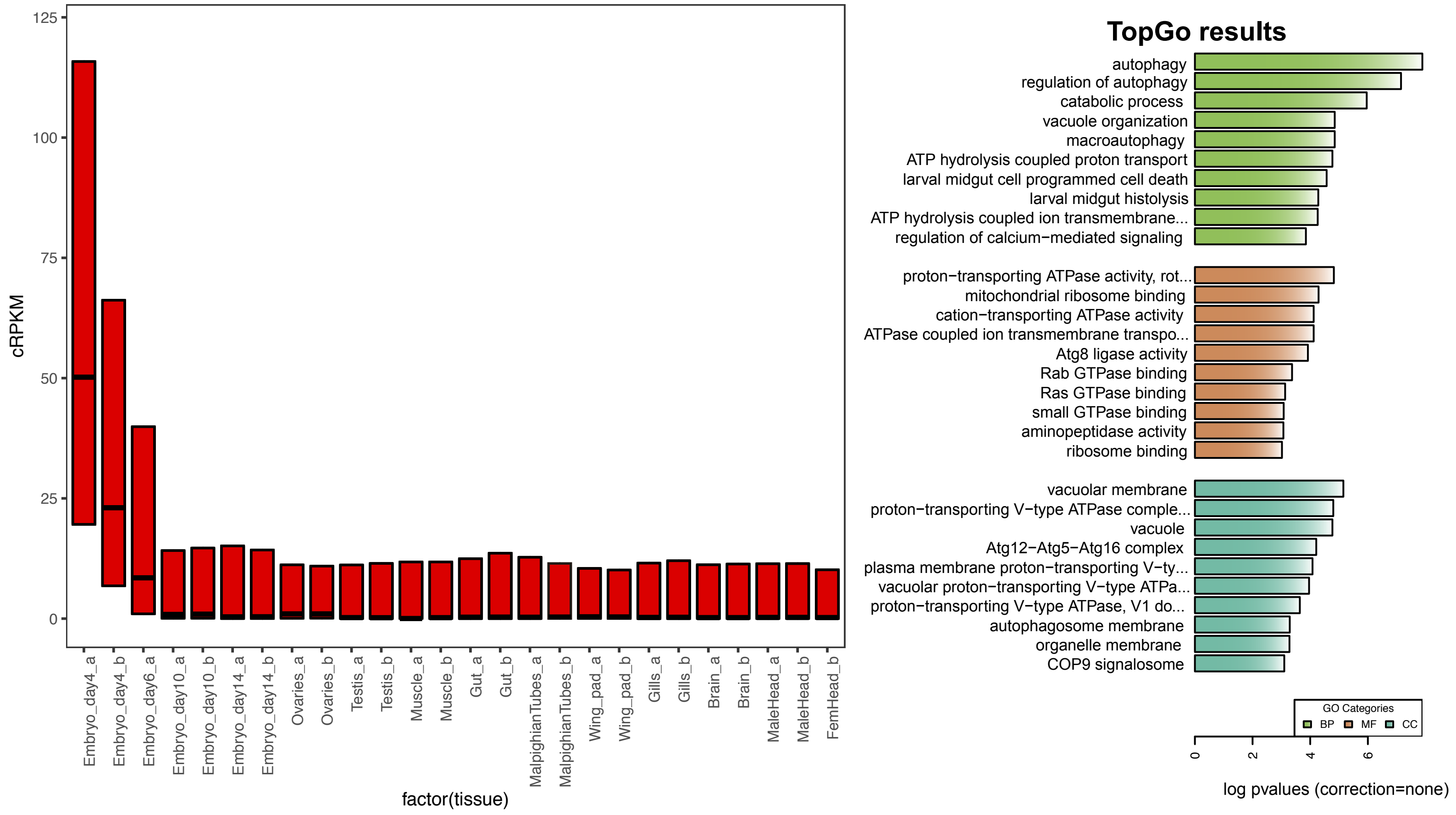

### *C. dipterum* modules

#### Module:saddlebrown (Embryogenesis- Synapsis)

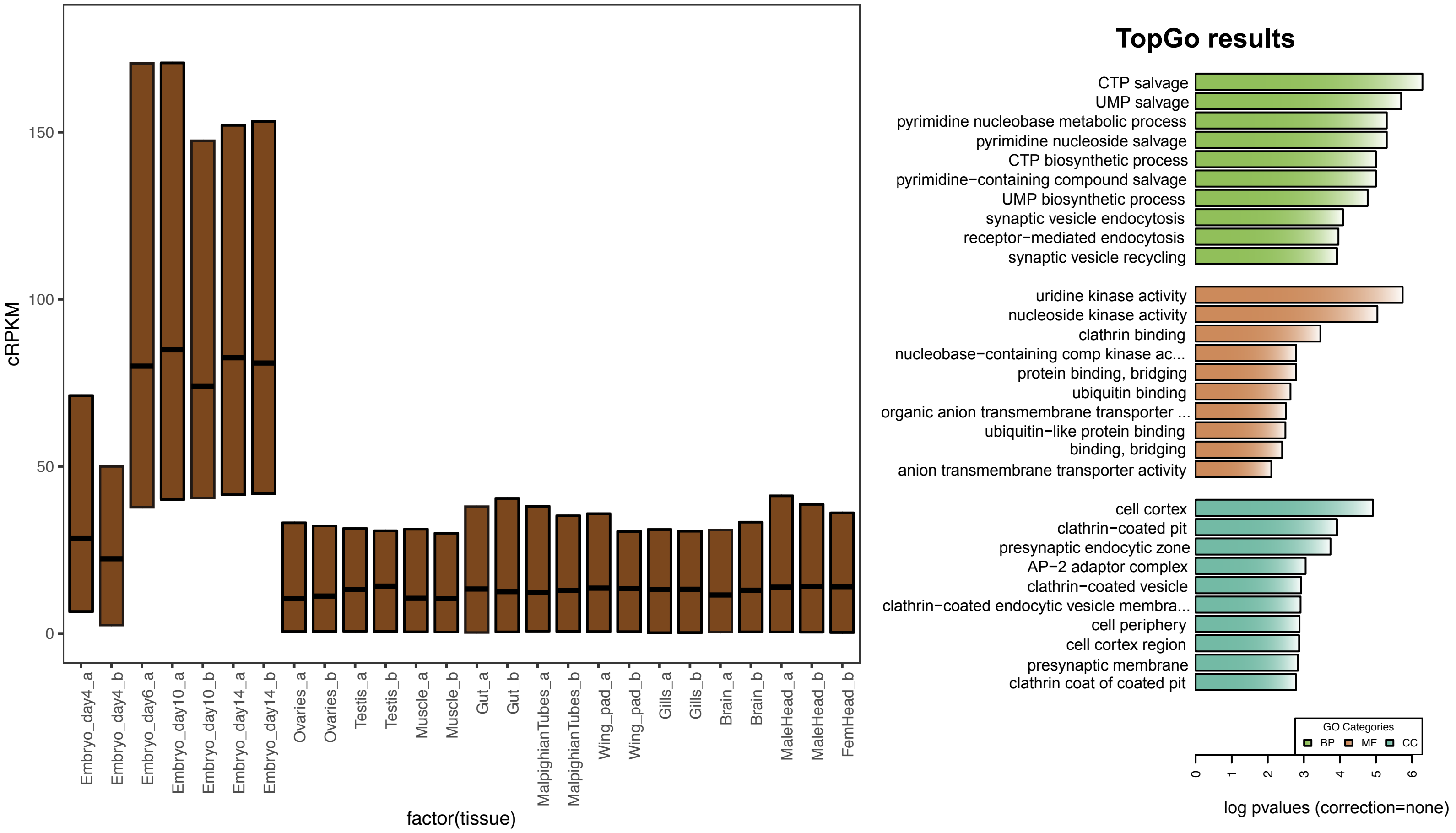


#### Module:green (Testis)

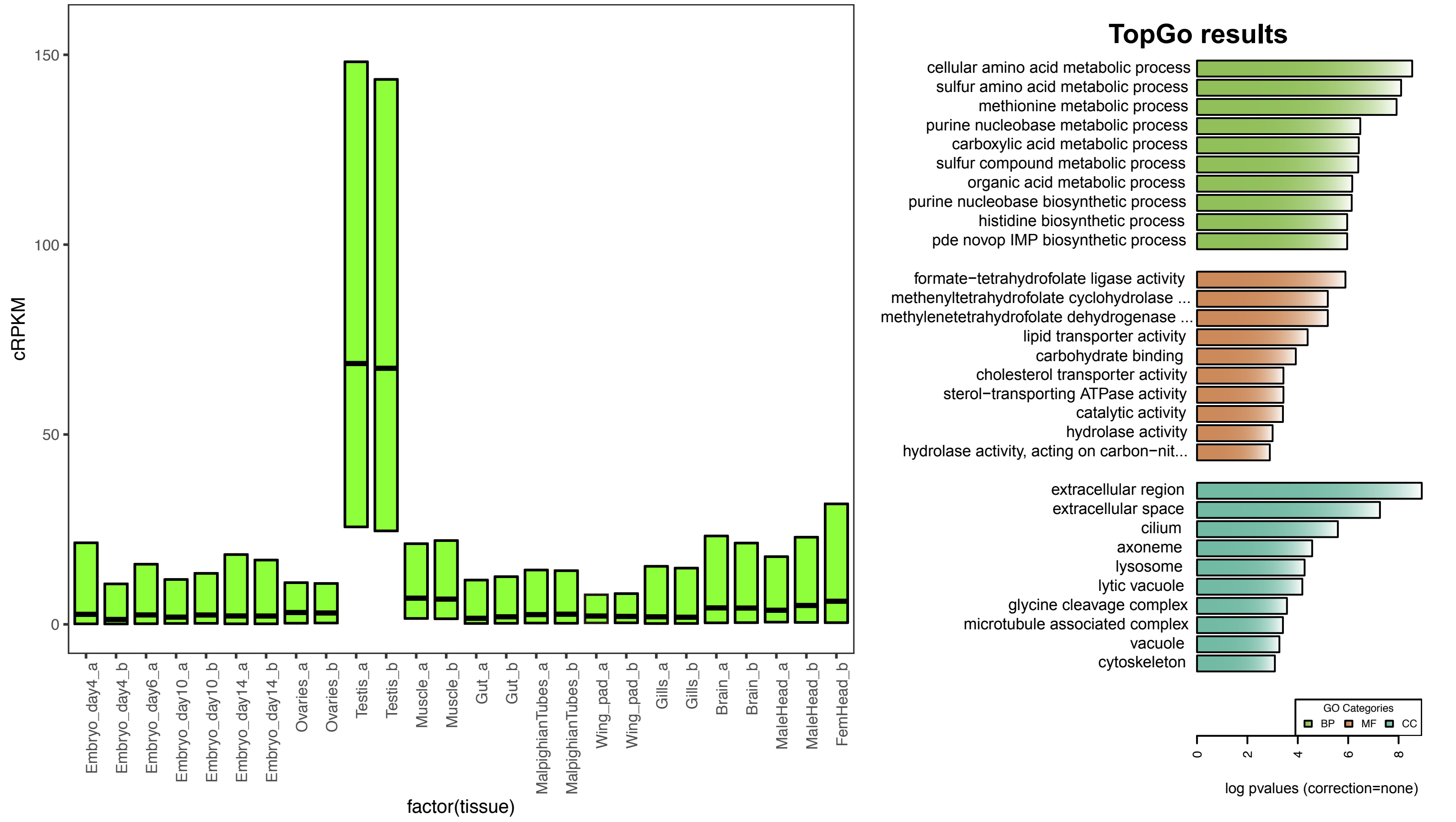

### *D. melanogaster* modules

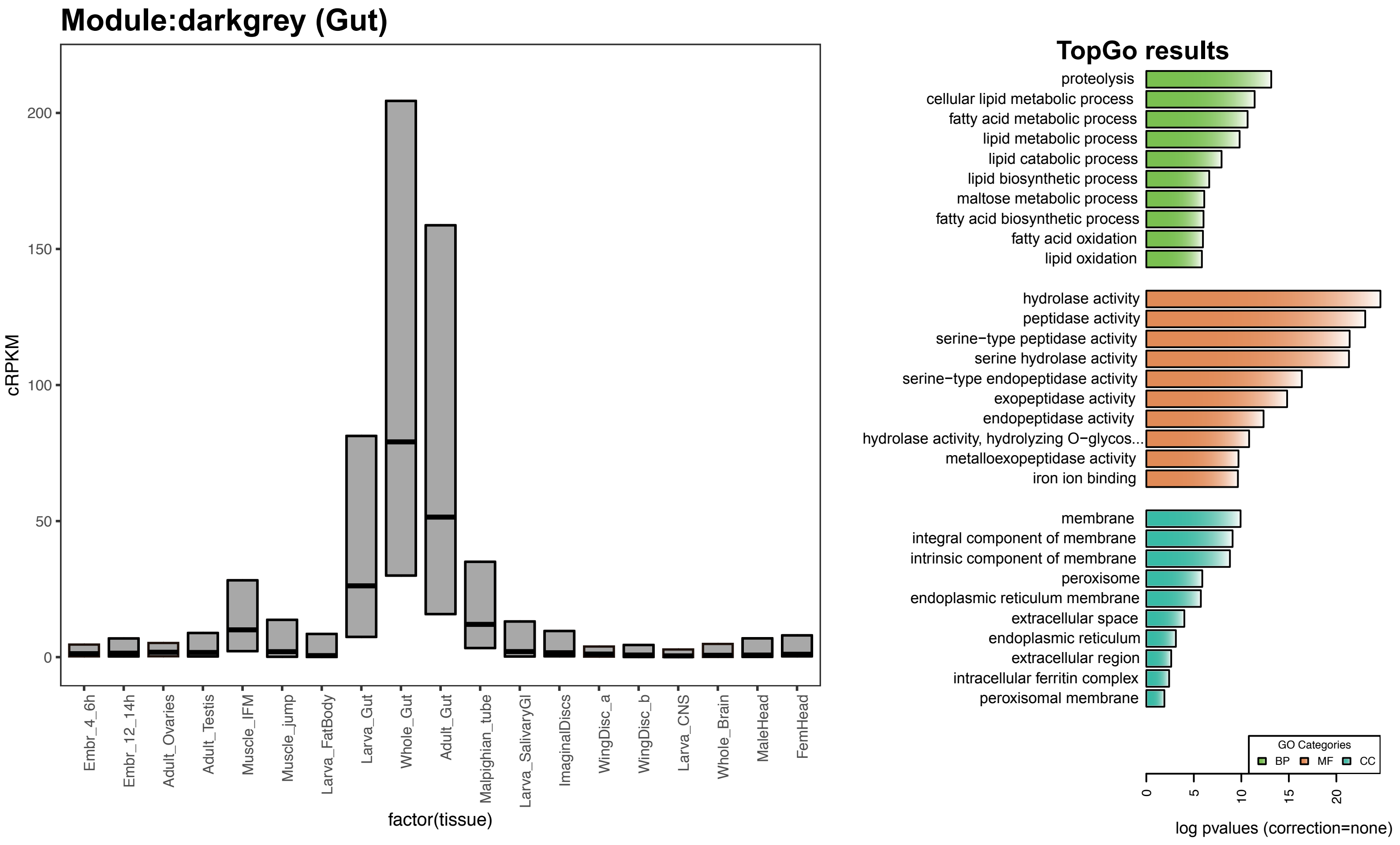


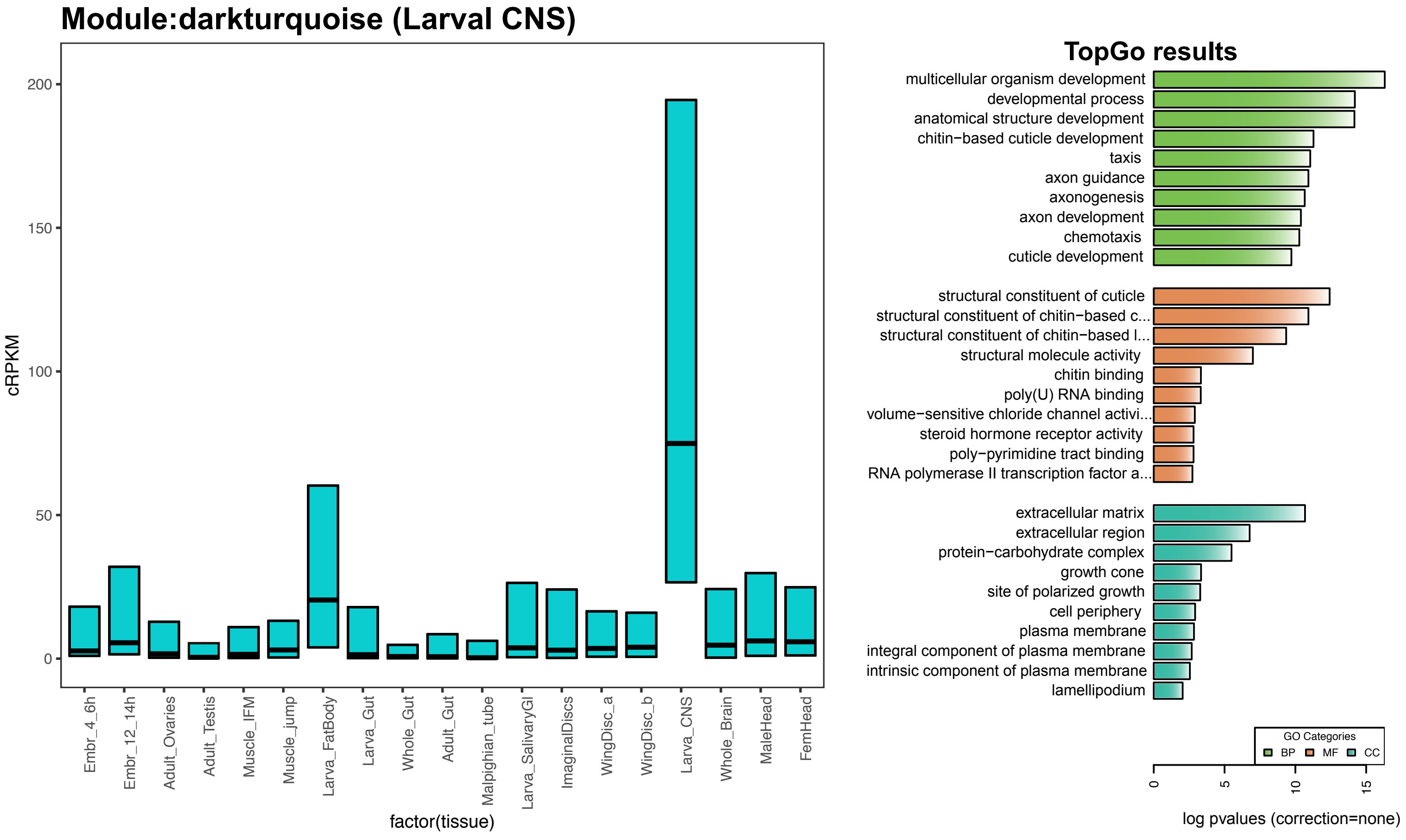

### *D. melanogaster* modules

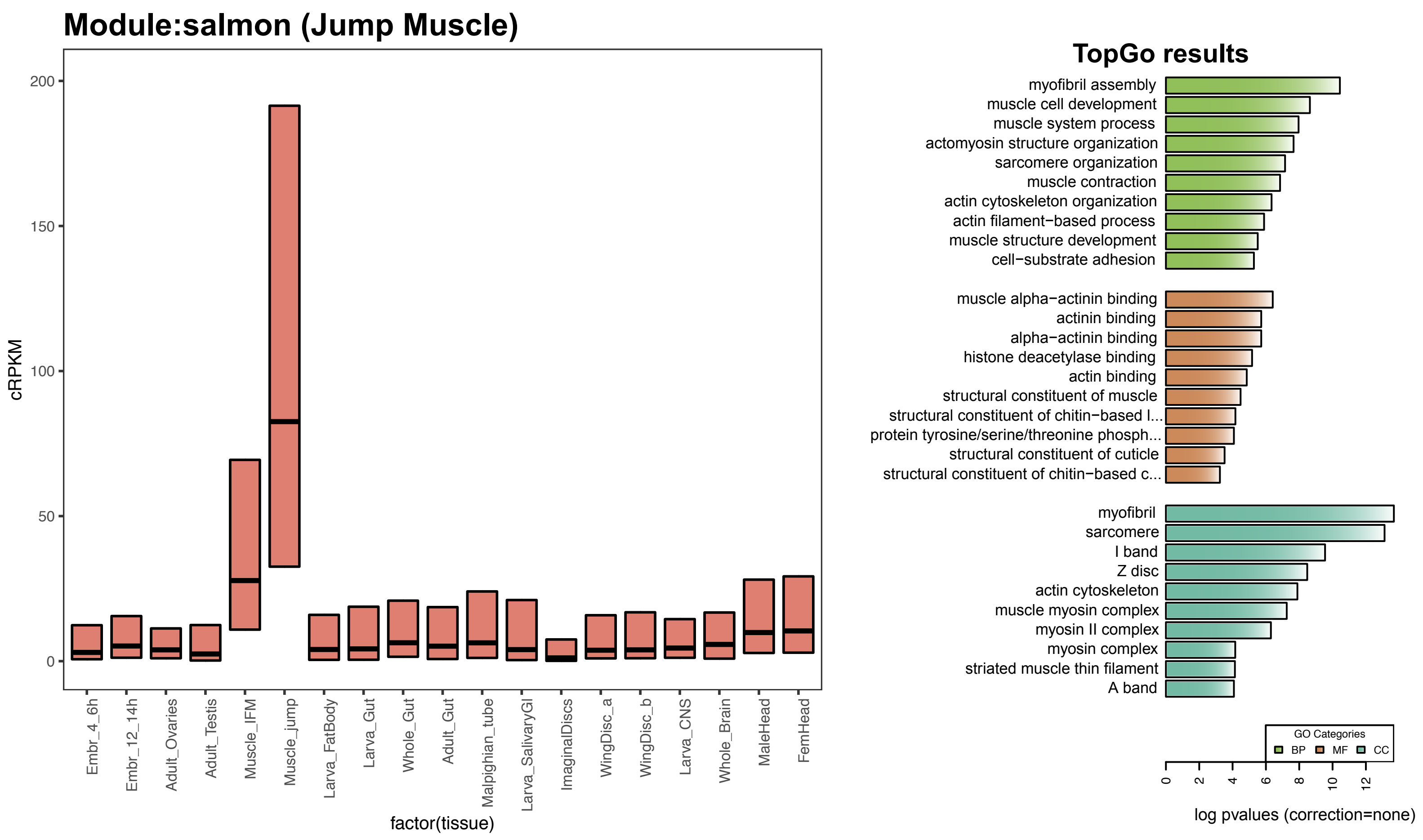


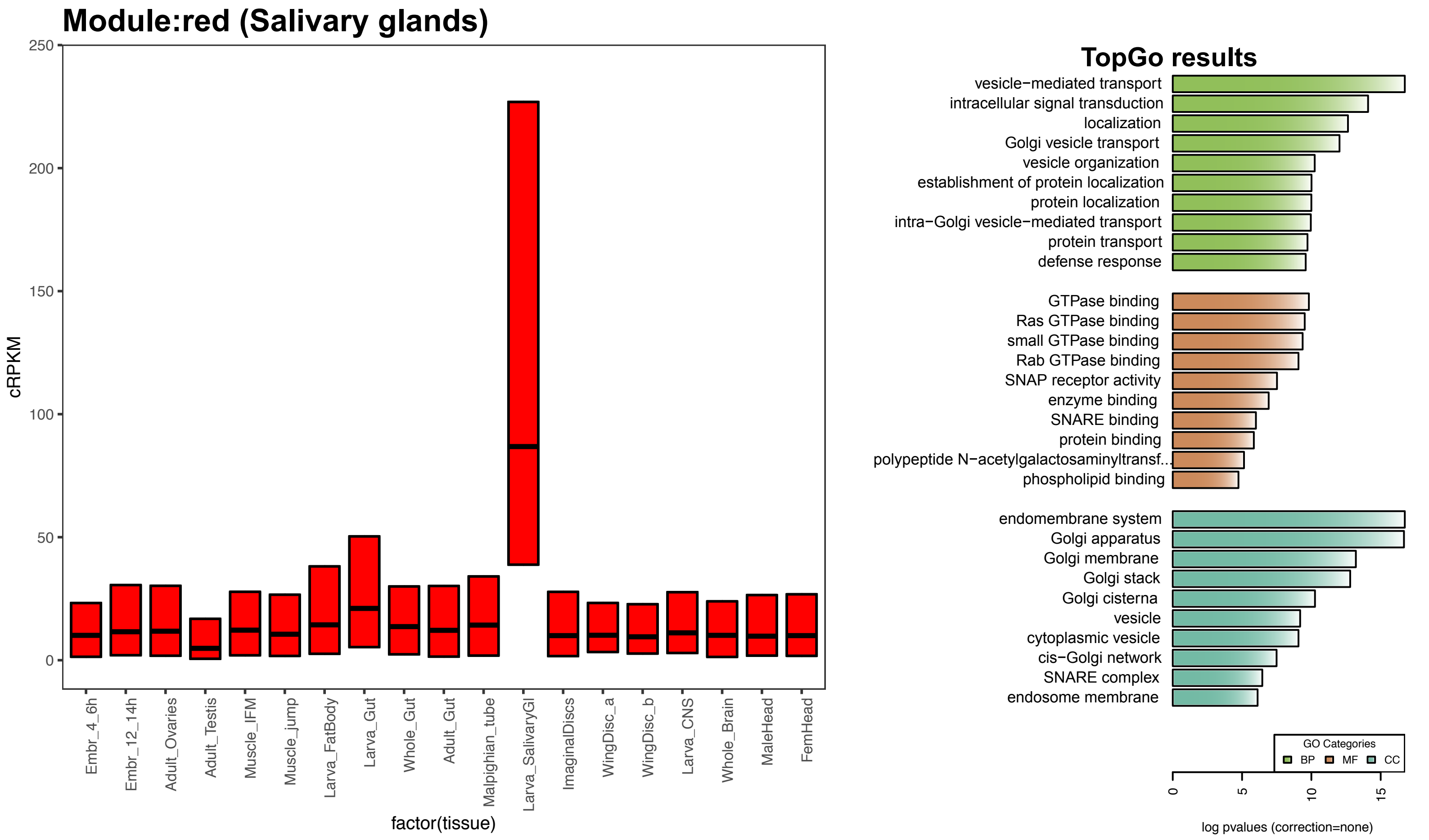

### *D. melanogaster* modules

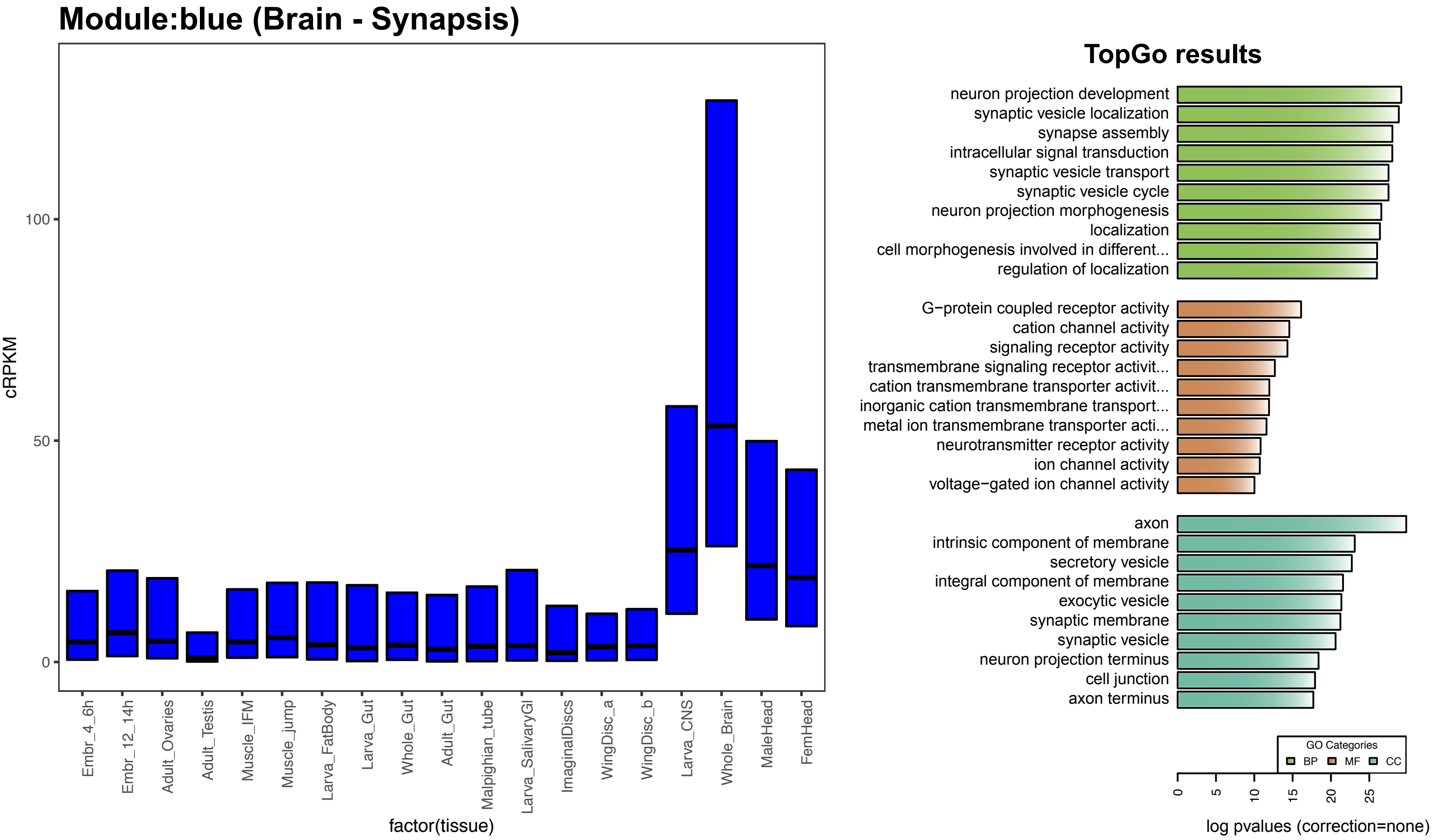


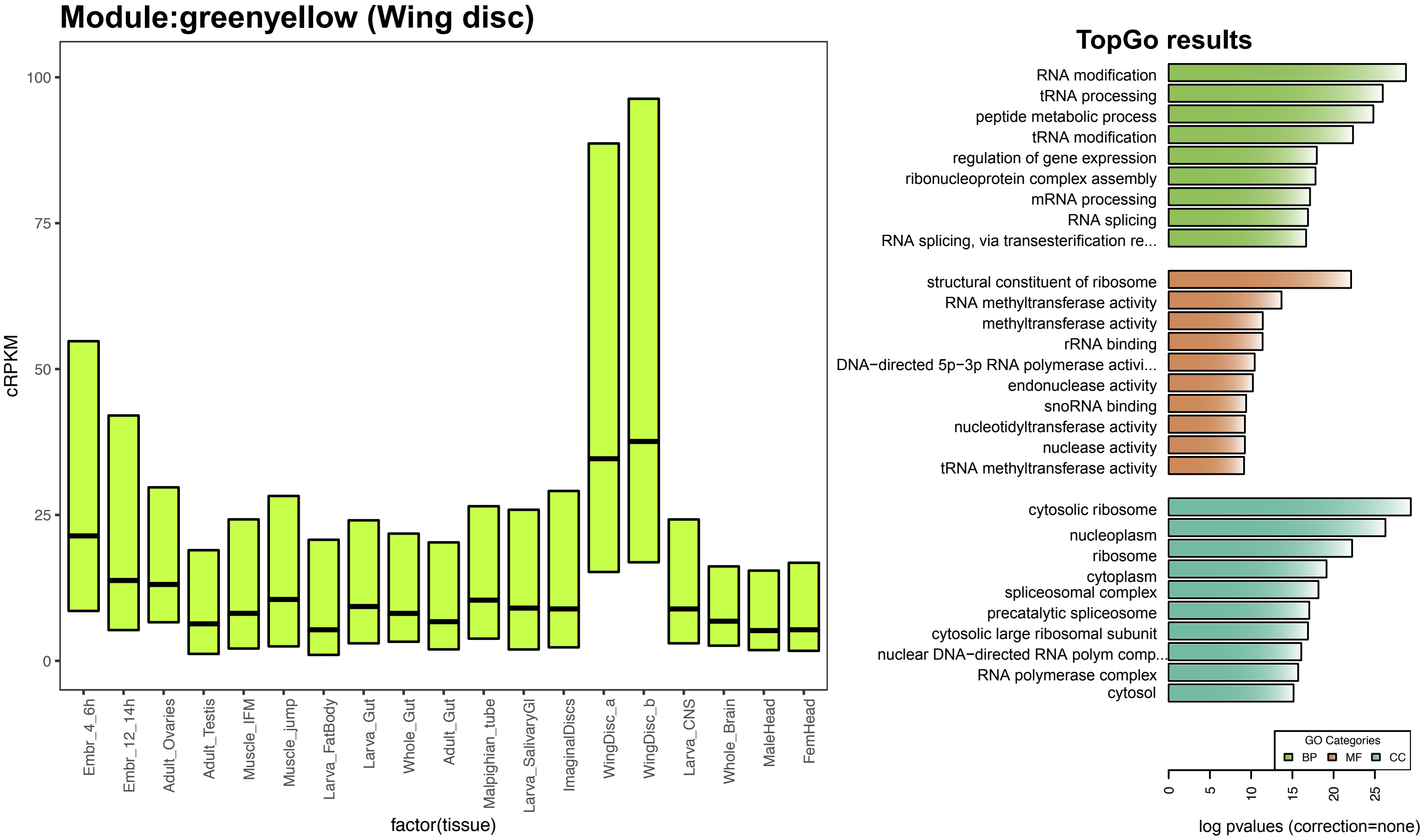

### *D. melanogaster* modules

#### Module:cyan (Embryo 12-14 hours)

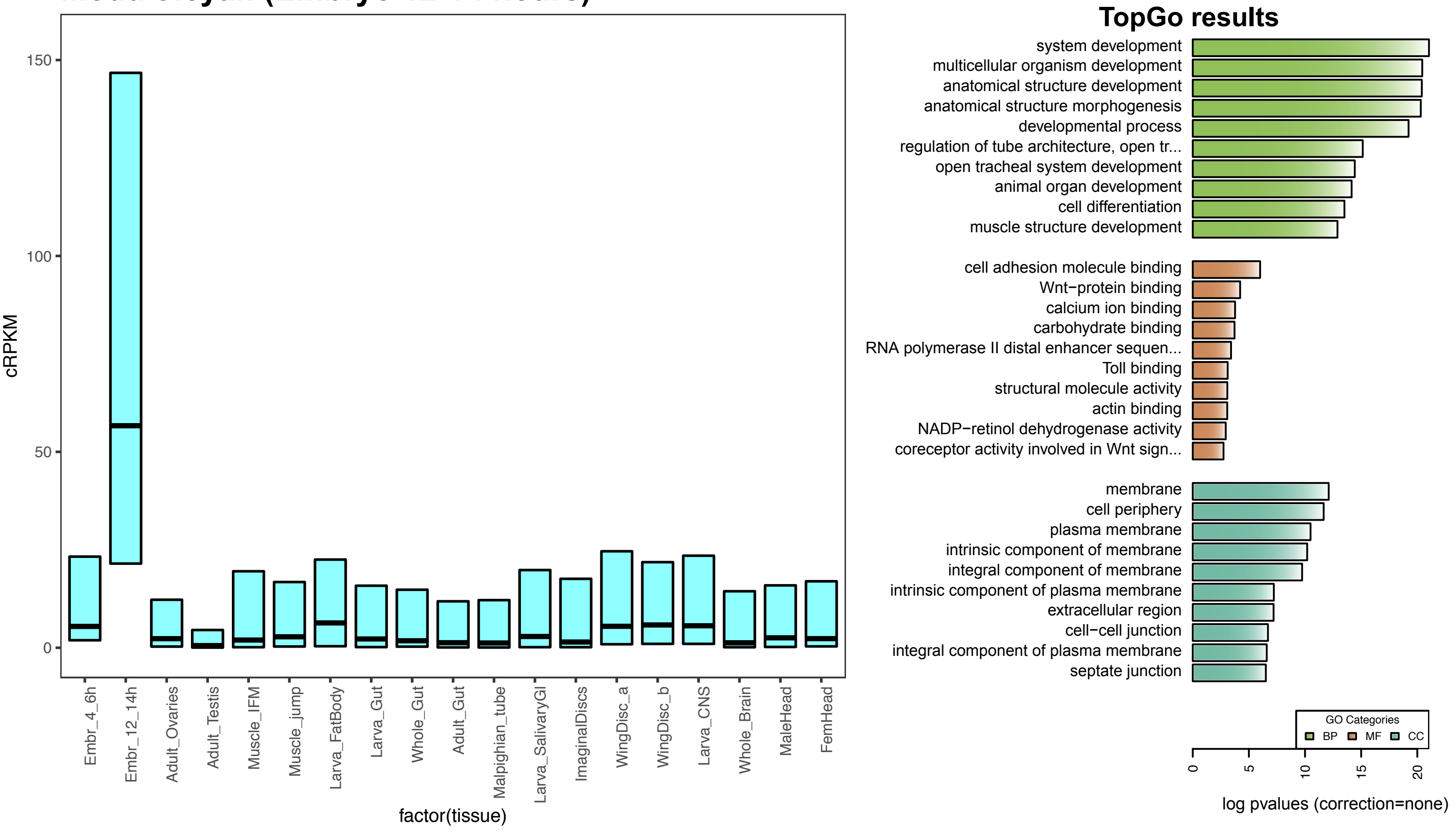


#### Module:black (Flight Muscle)

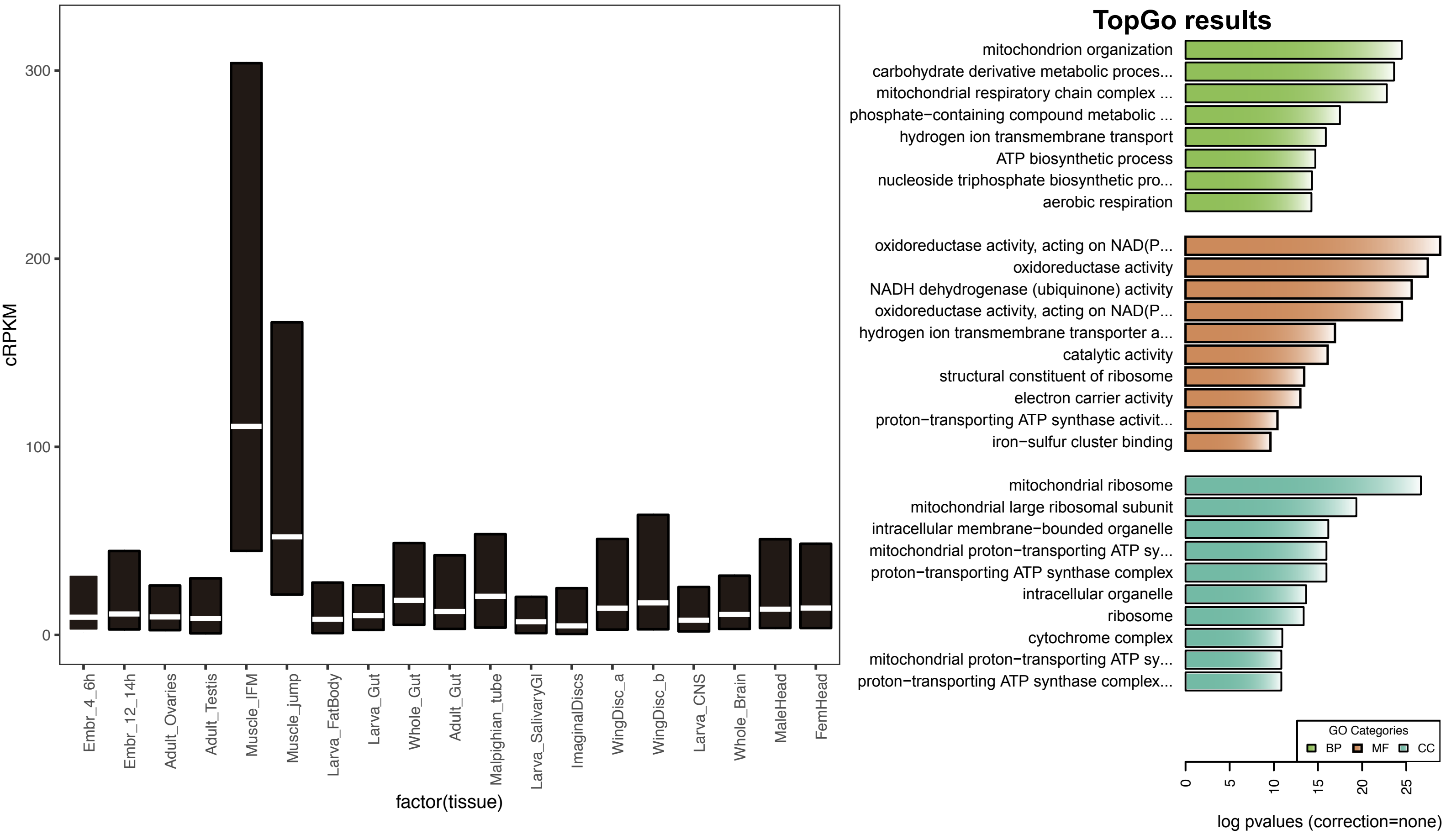

### *D. melanogaster* modules

#### Module:magenta (Embryo 4-6 hours- Transcription)

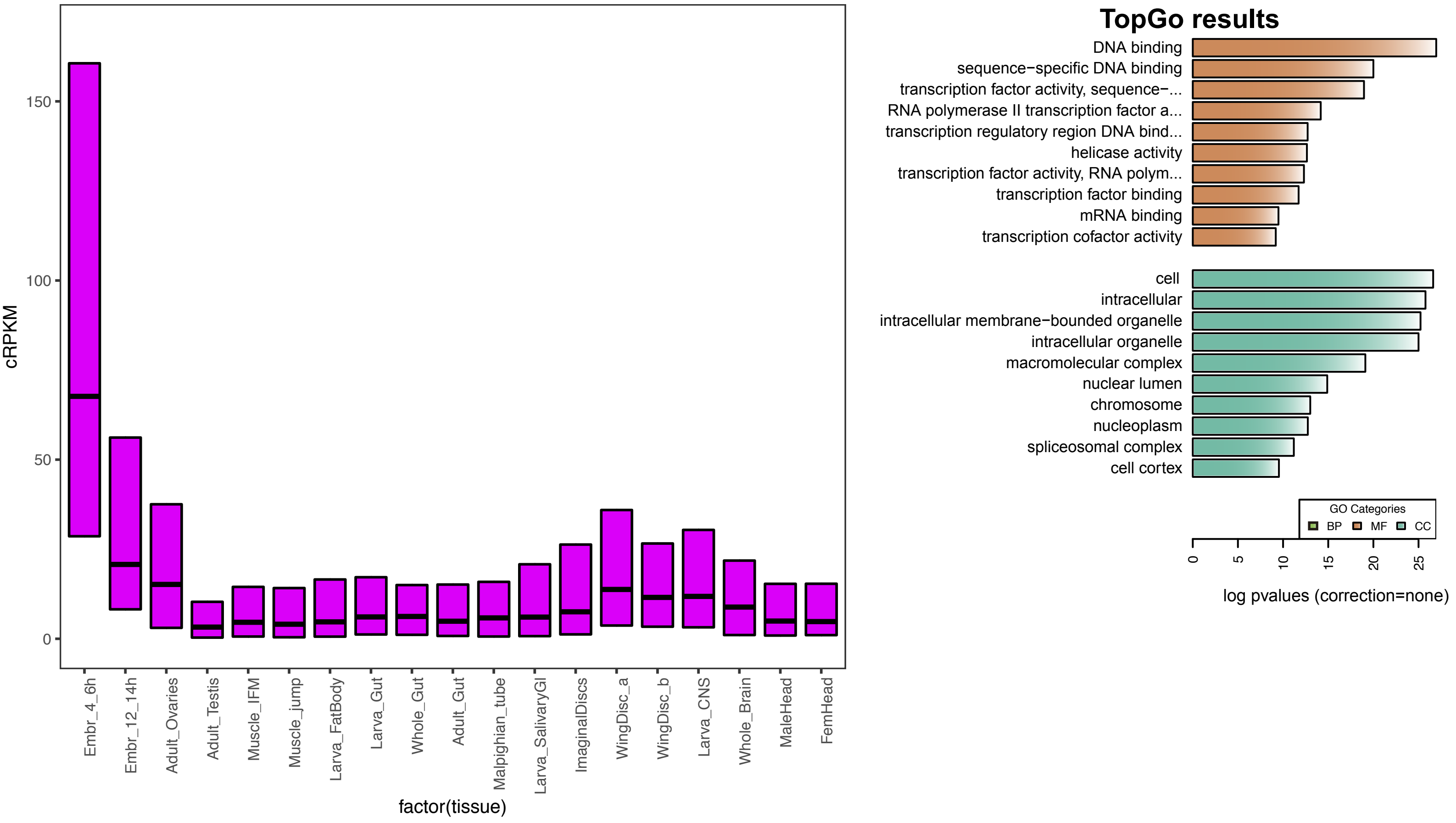


#### Module:purple (Adult heads- Visual perception)

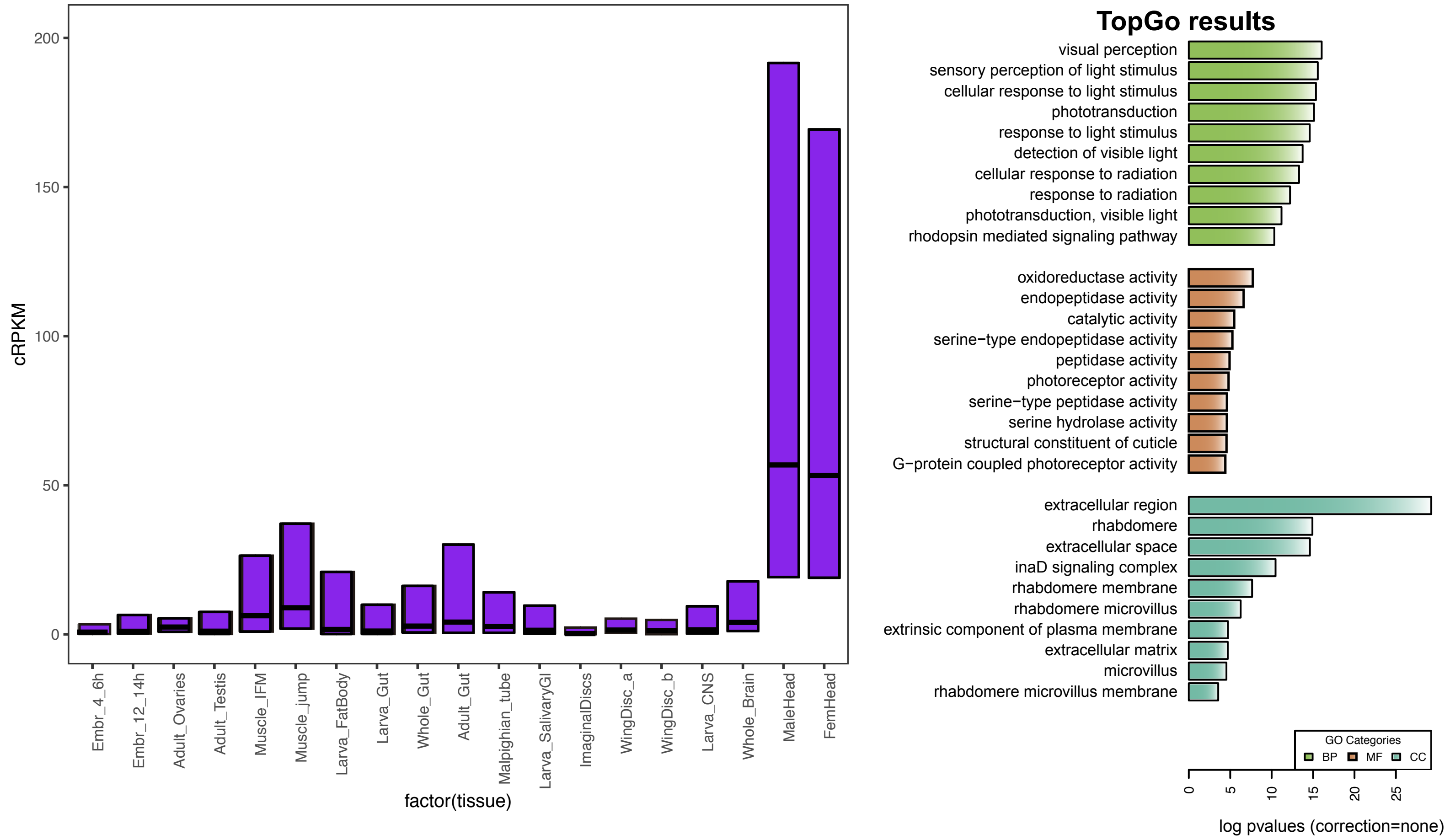

### *D. melanogaster* modules

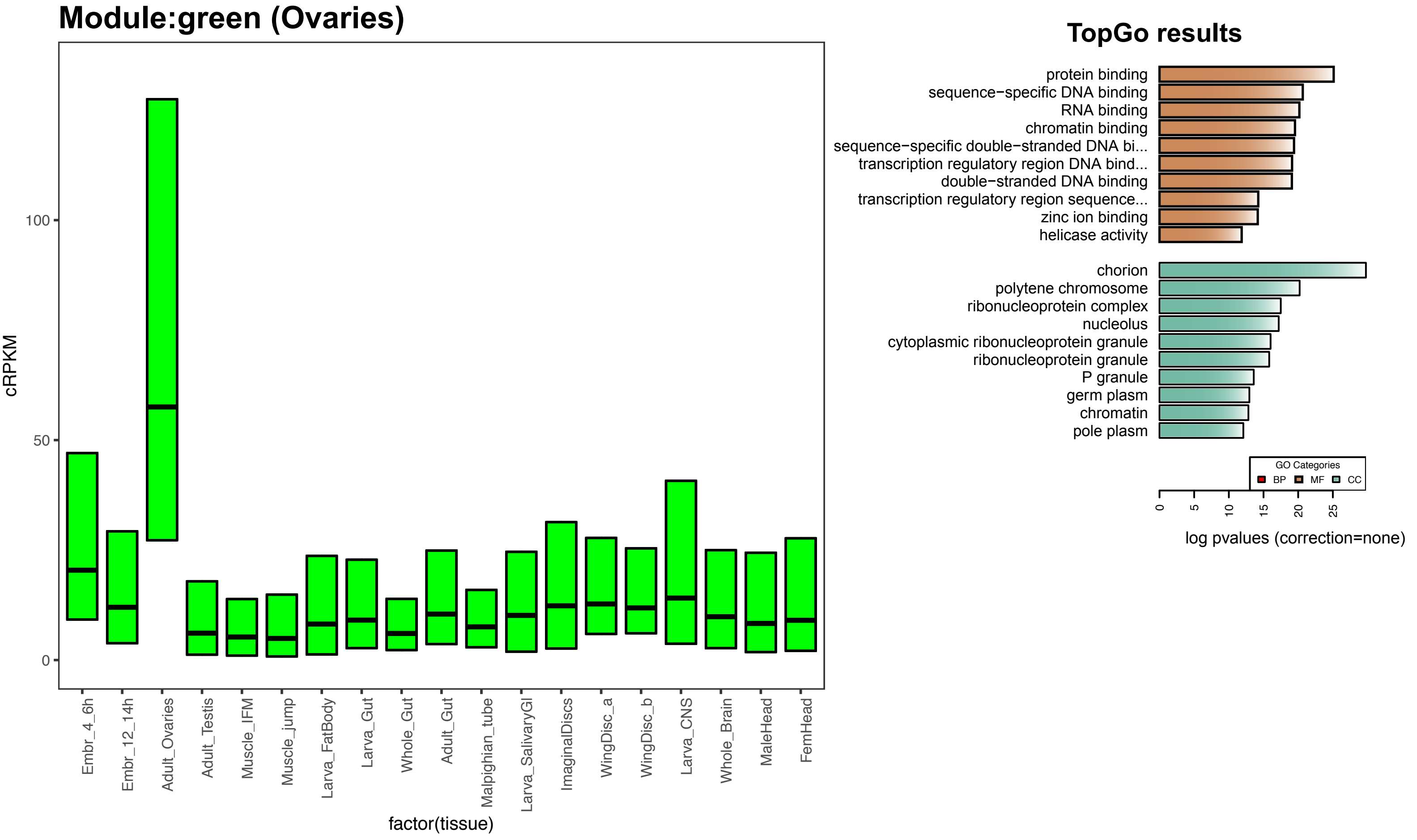


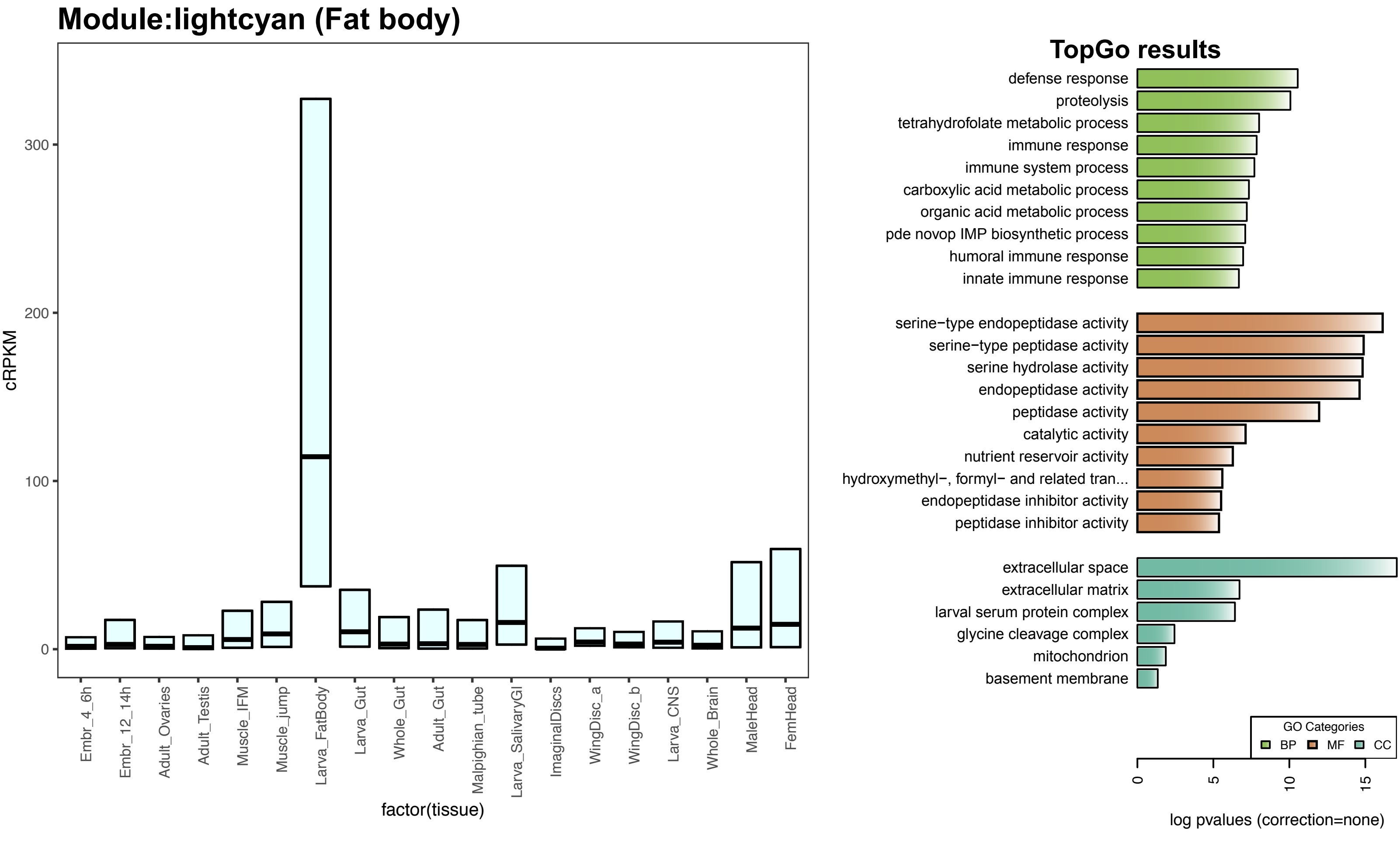

### *D. melanogaster* modules

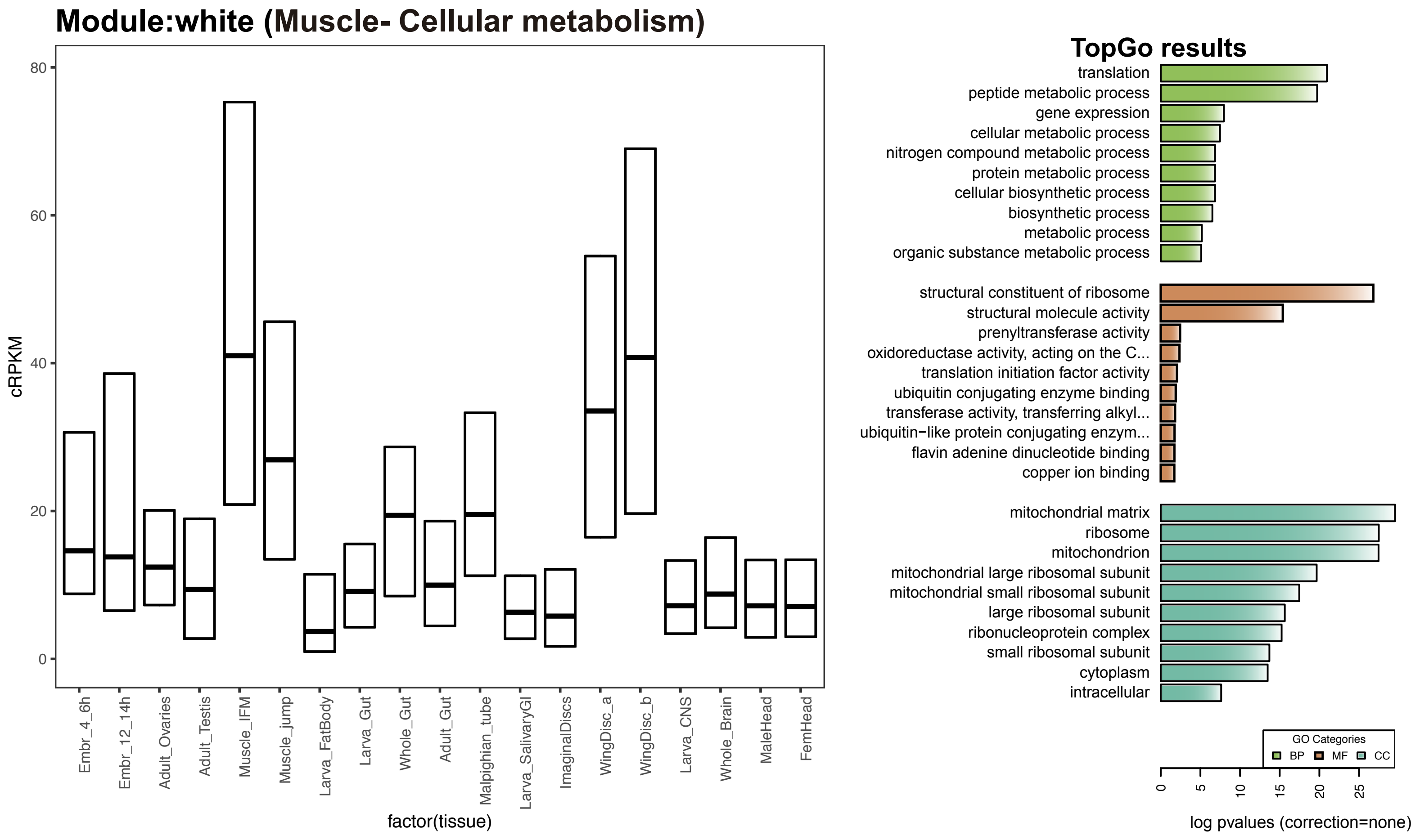


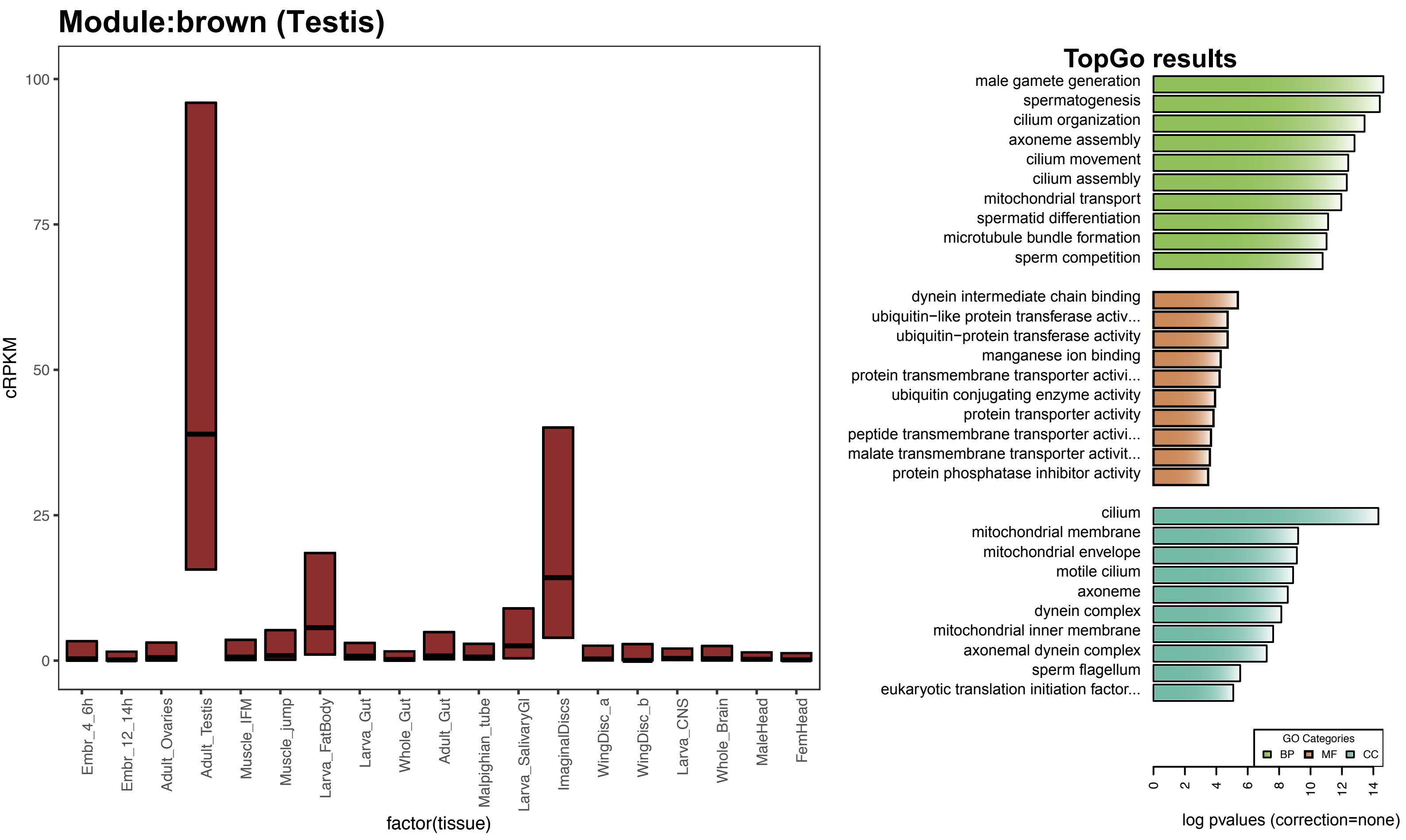

### *D. melanogaster* modules

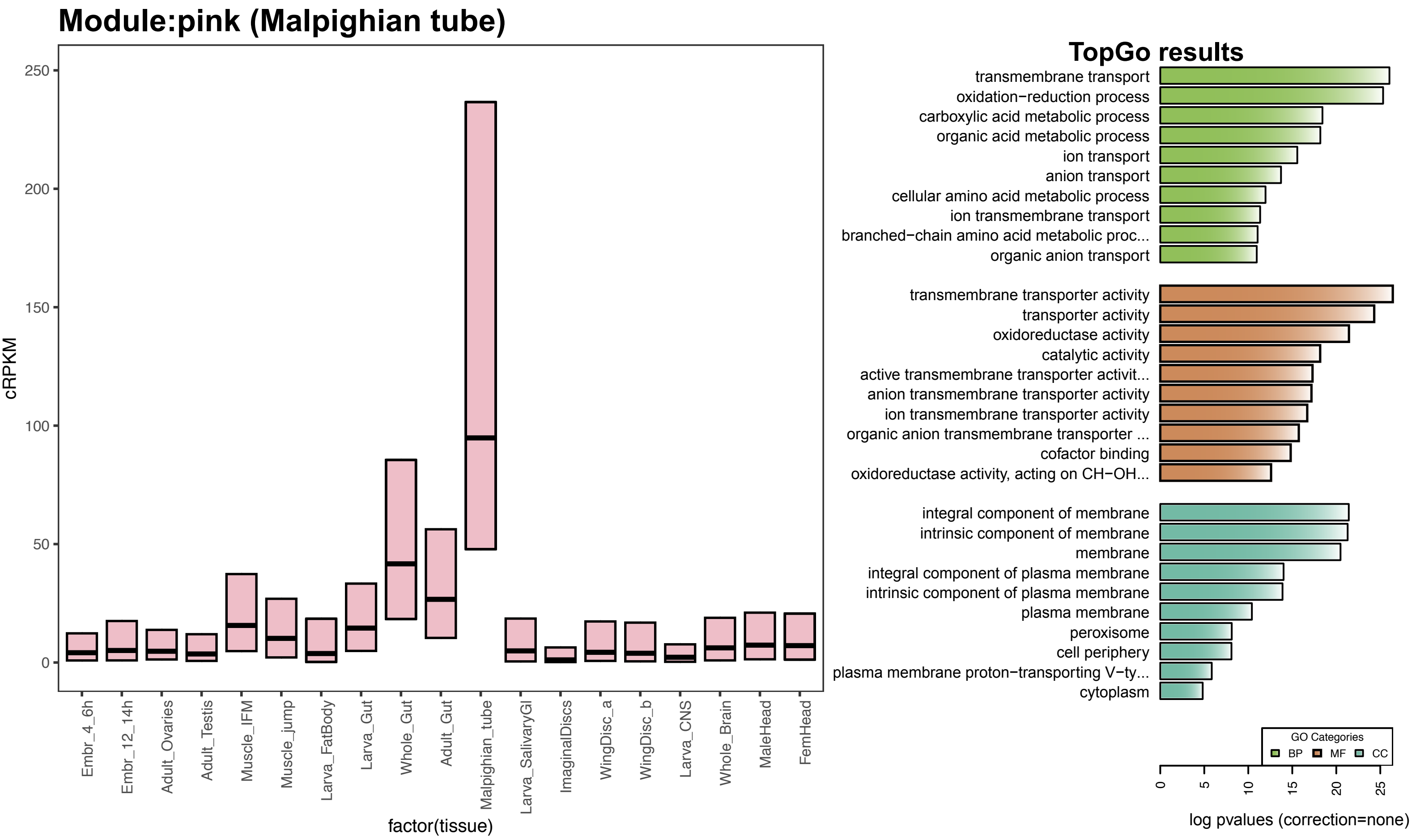


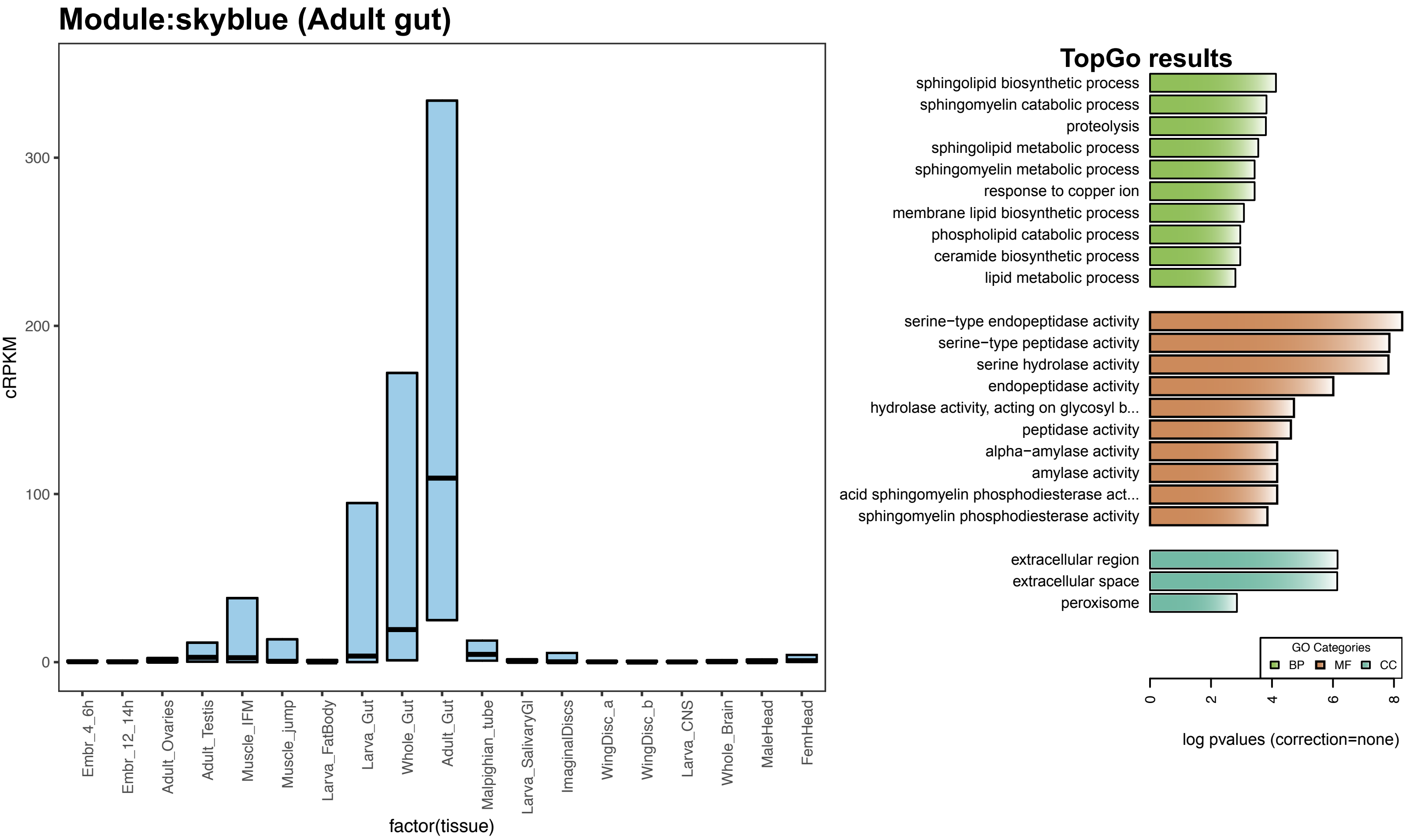

### *D. melanogaster* modules

#### Module:grey60 (Larval gut)

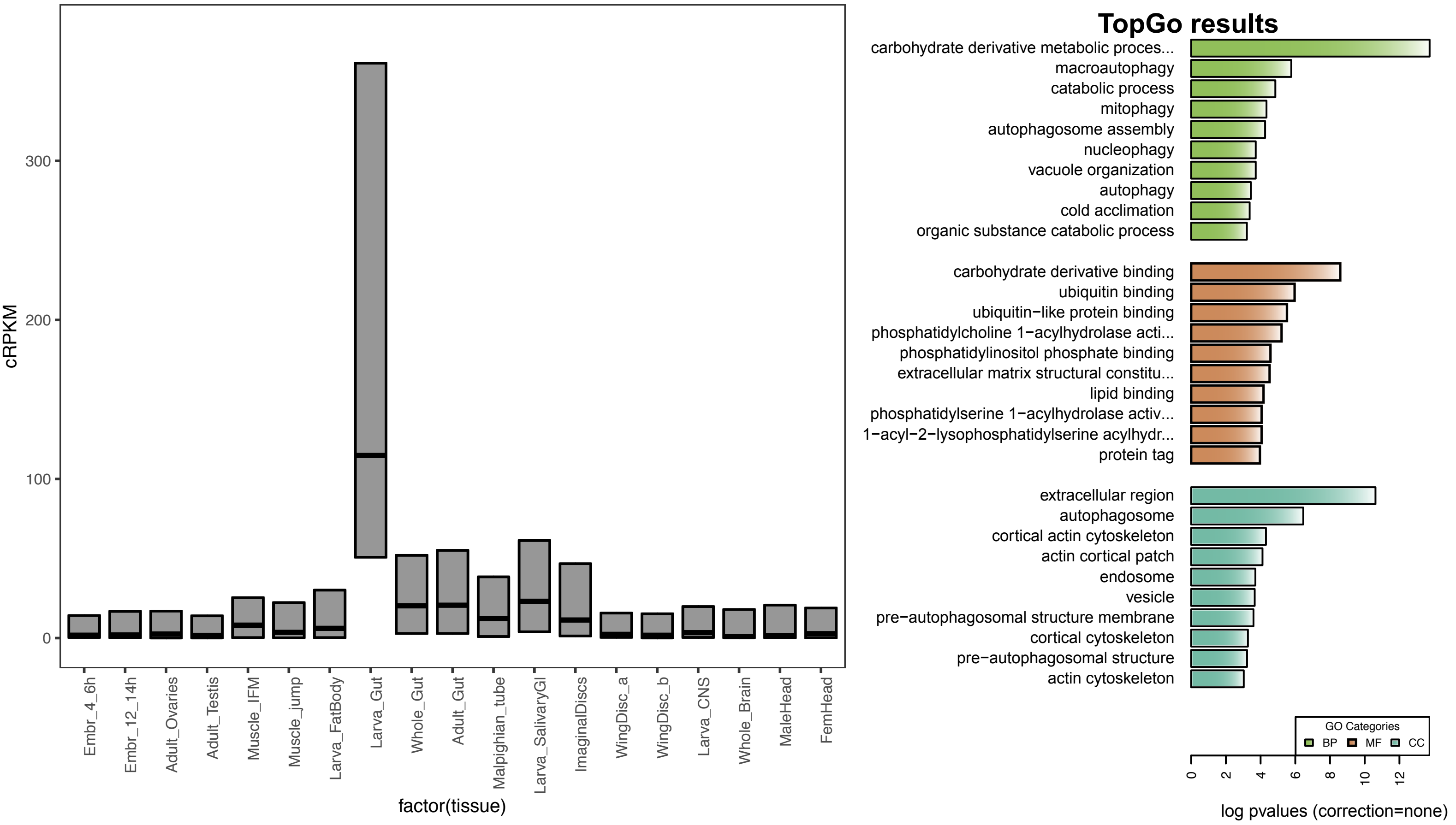

### *S. maritima* modules

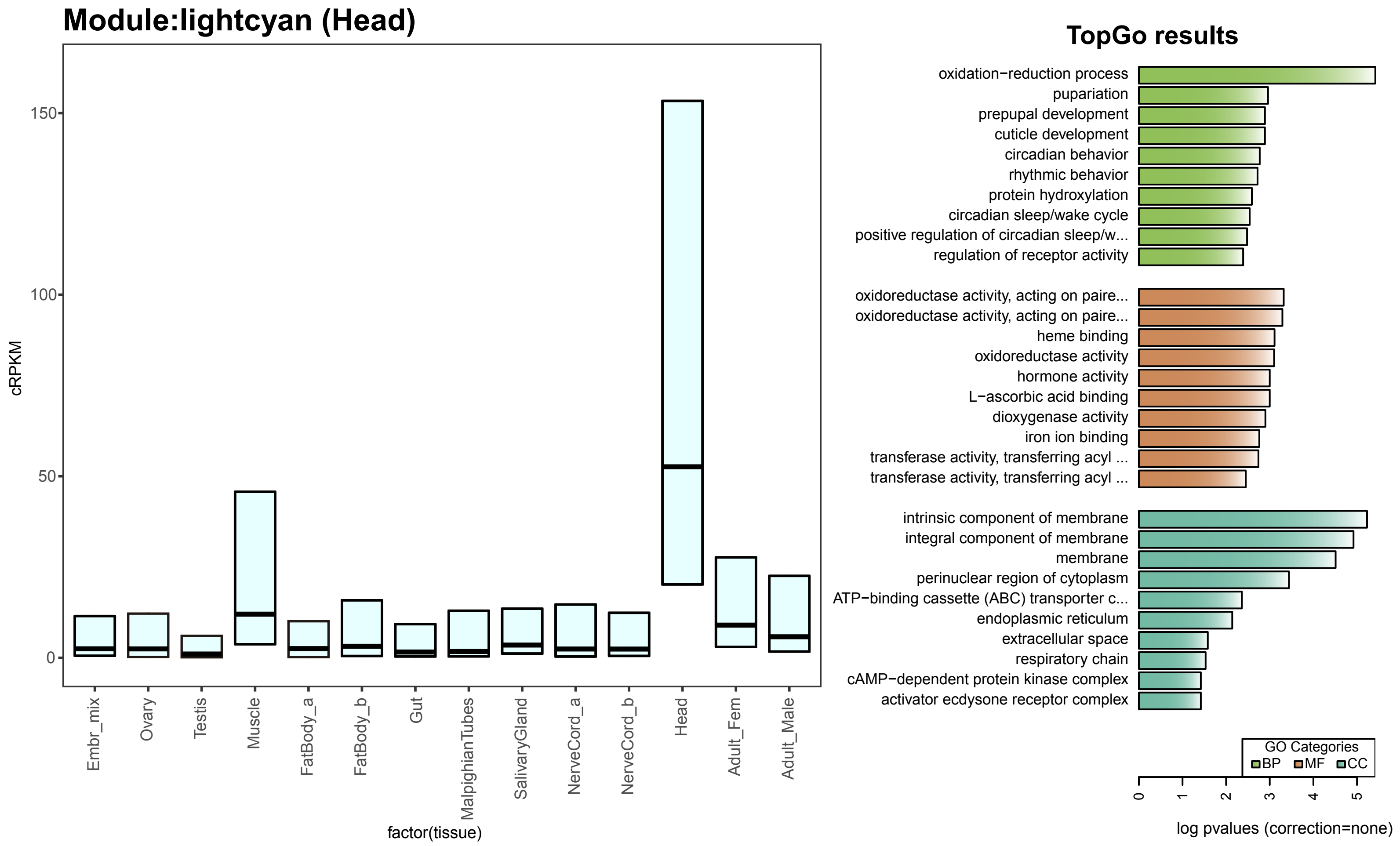


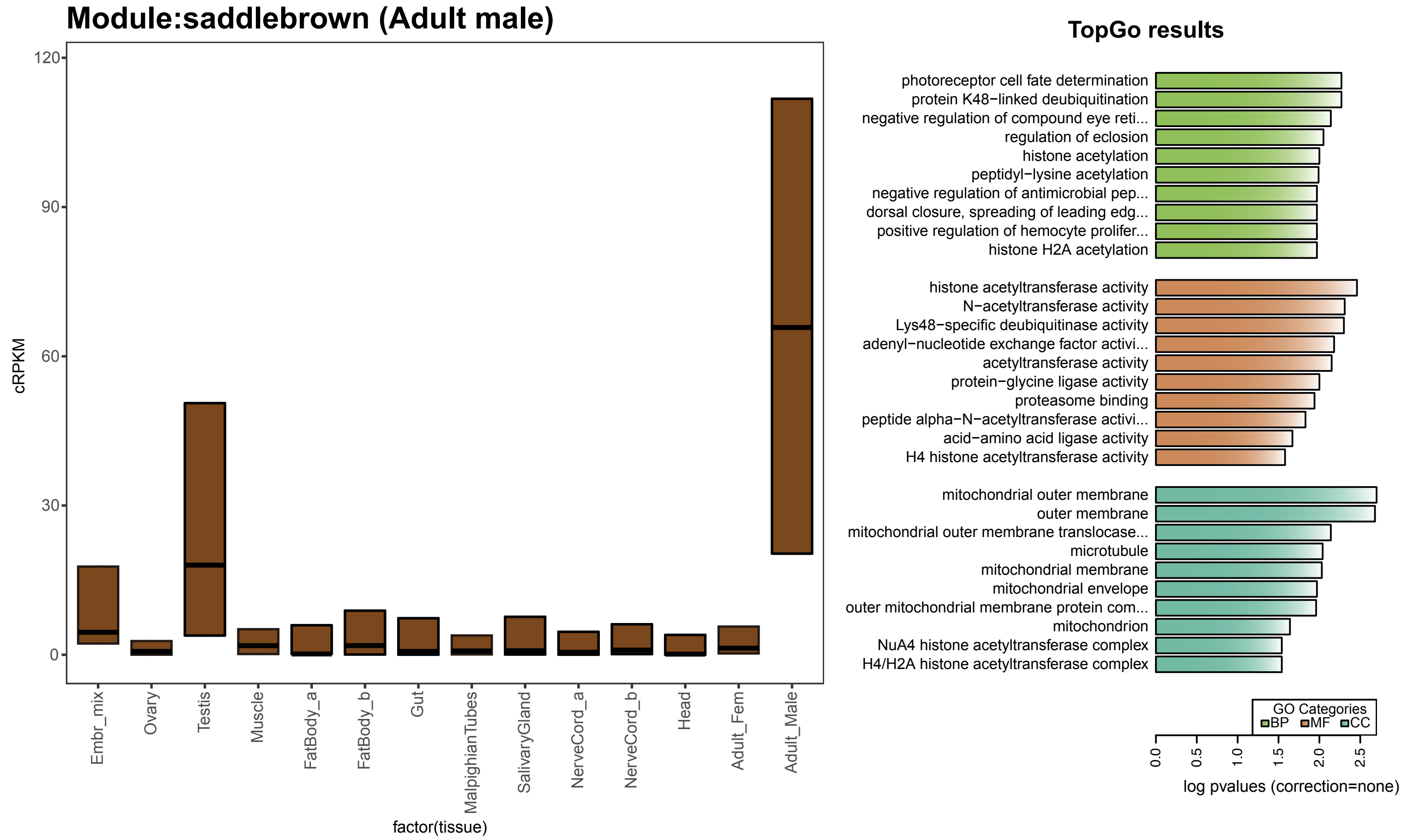

### *S. maritima* modules

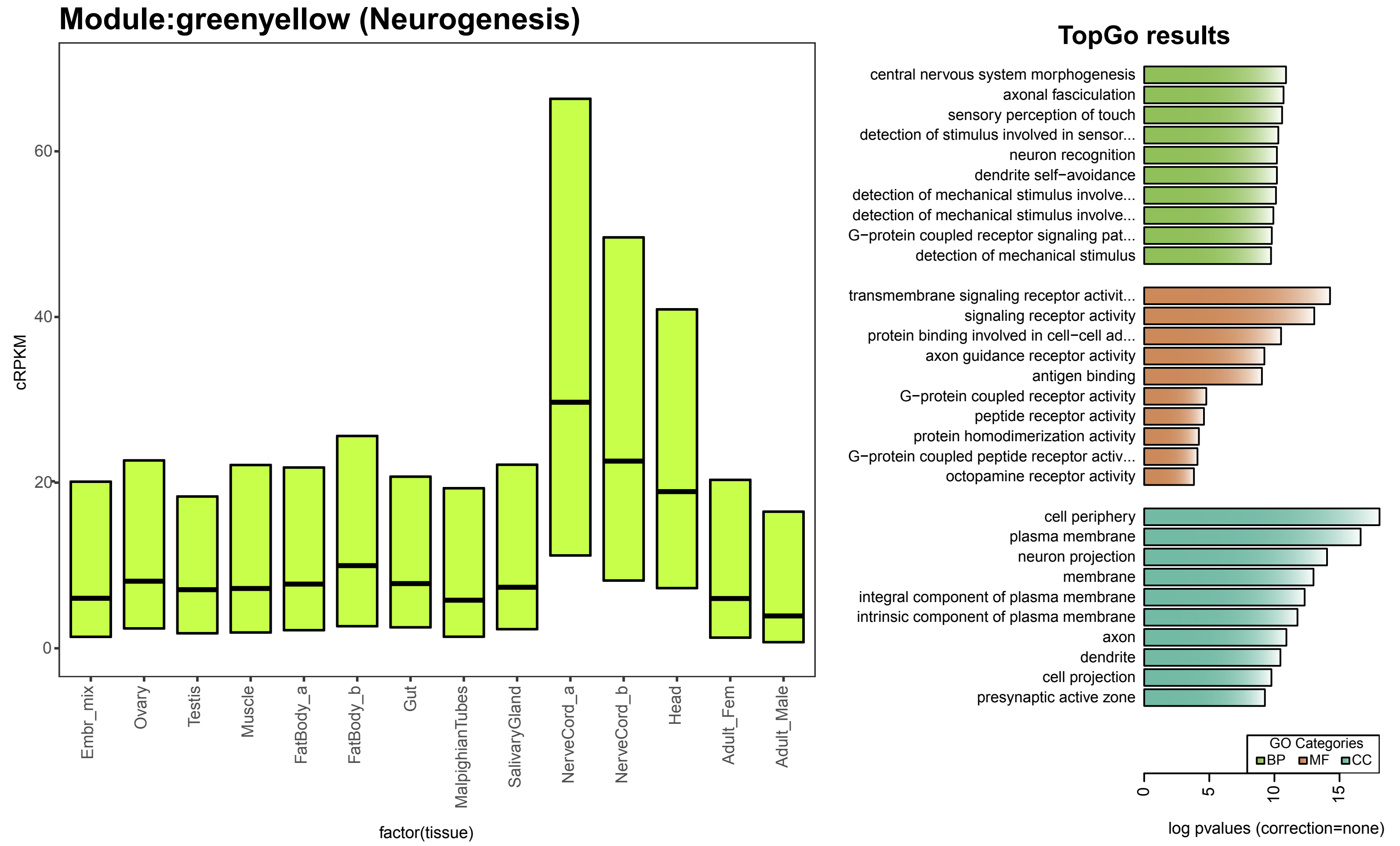


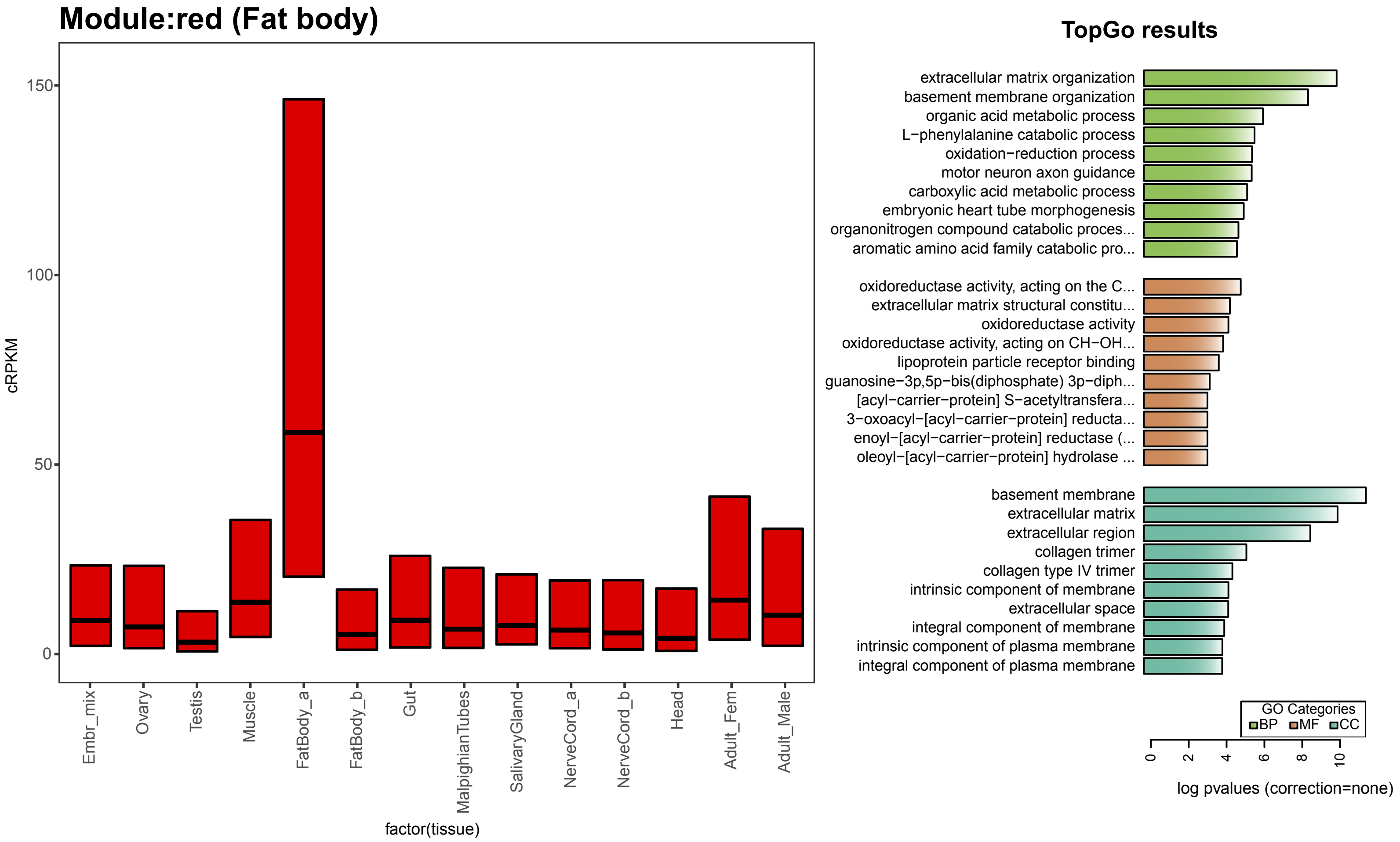

### *S. maritima* modules

#### Module:darkgreen (membrane)

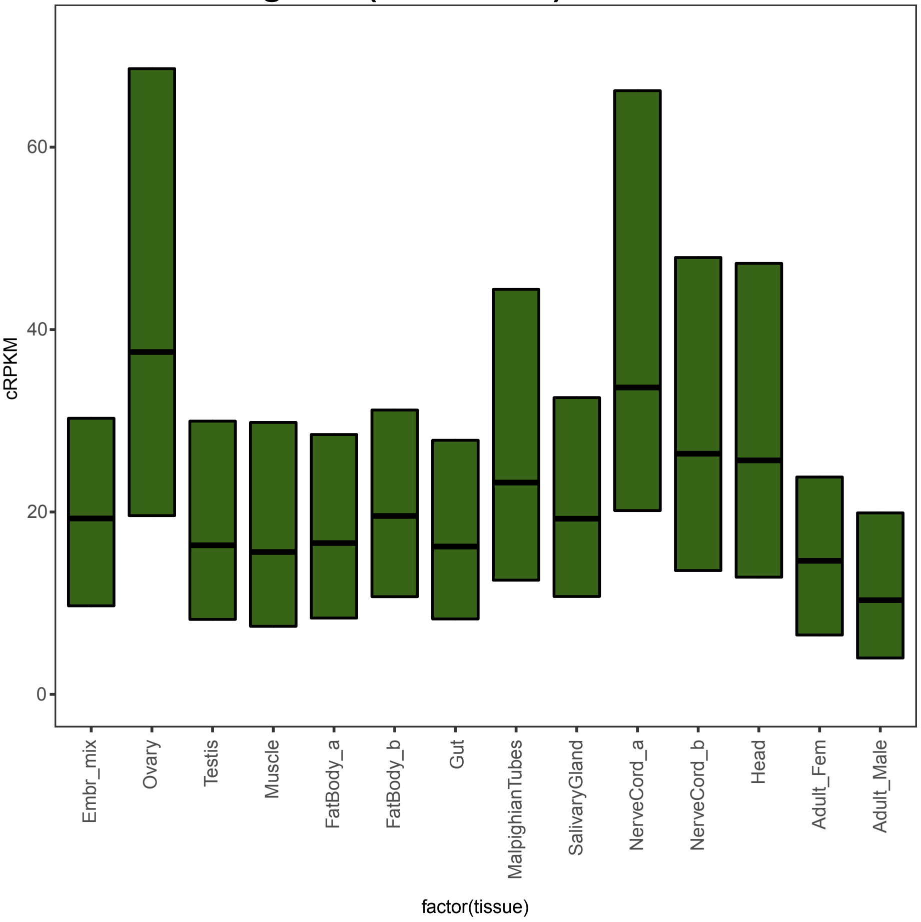


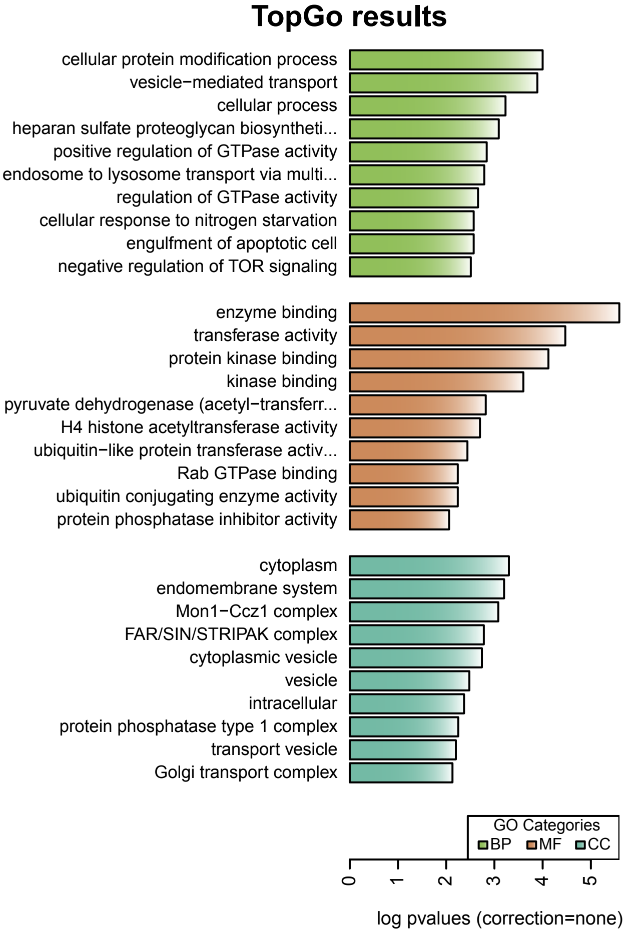


#### Module:skyblue (Embryo/Nerve cord)

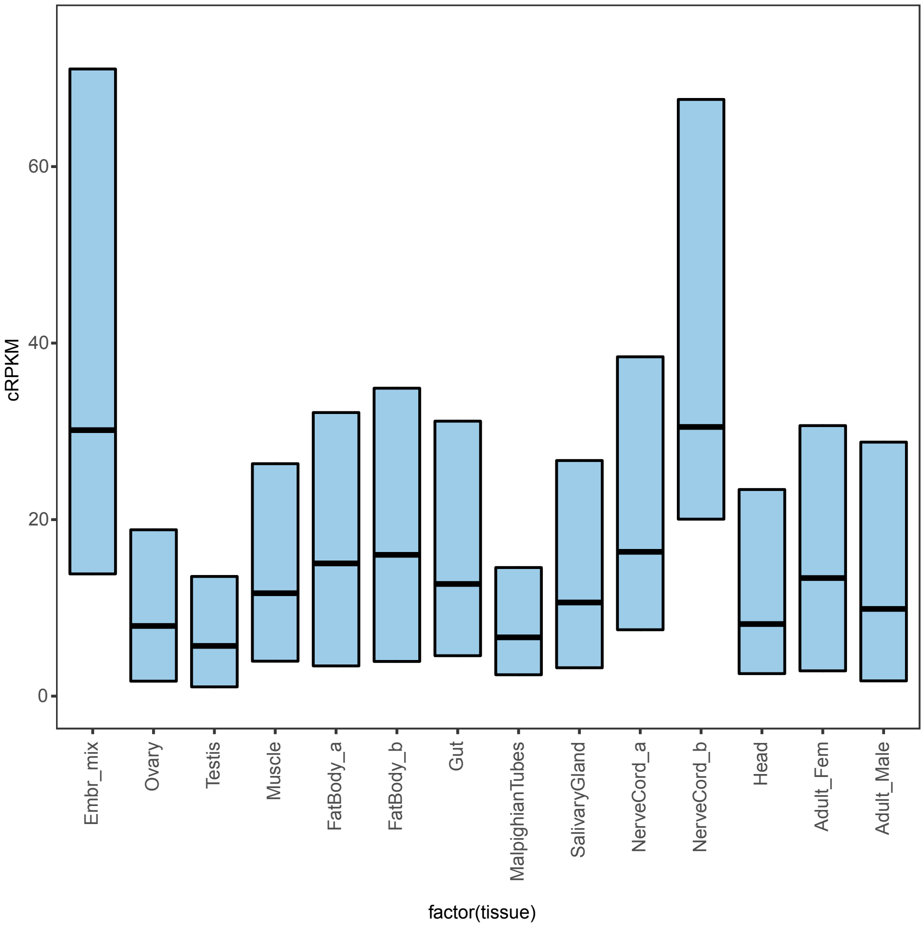


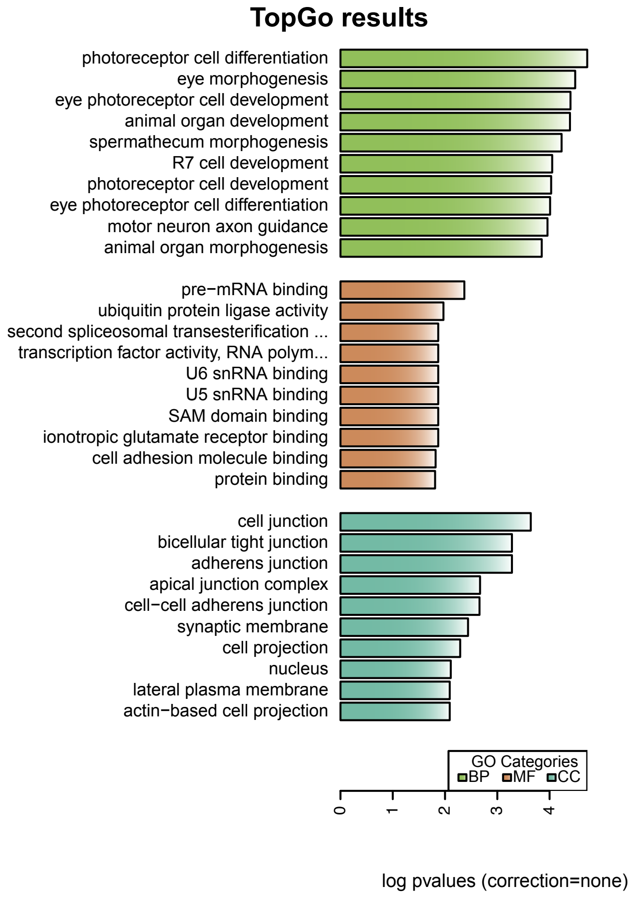

### *S. maritima* modules

#### Module:orange (Protein synthesis)

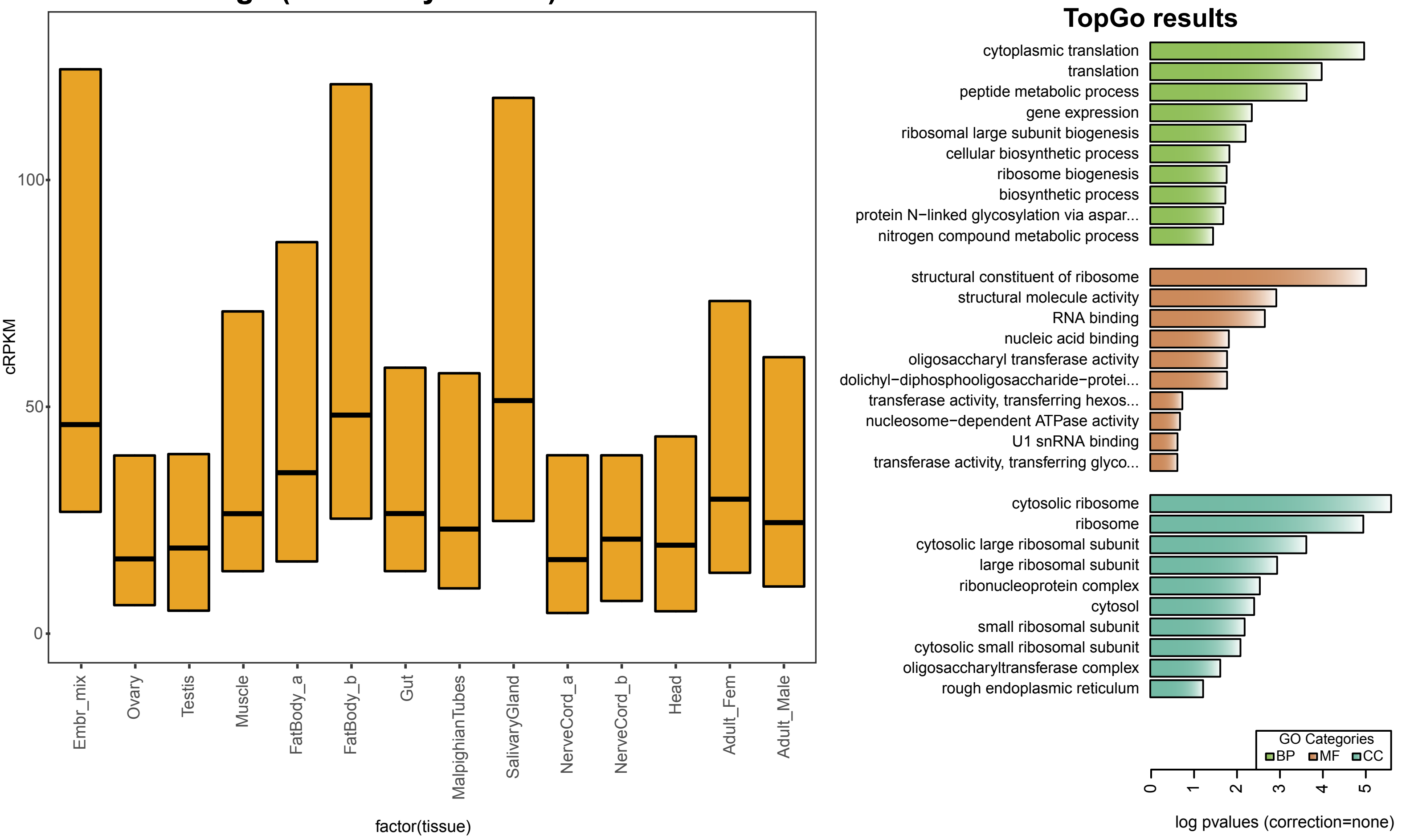


#### Module:black (Nerve Cord- Synapsis)

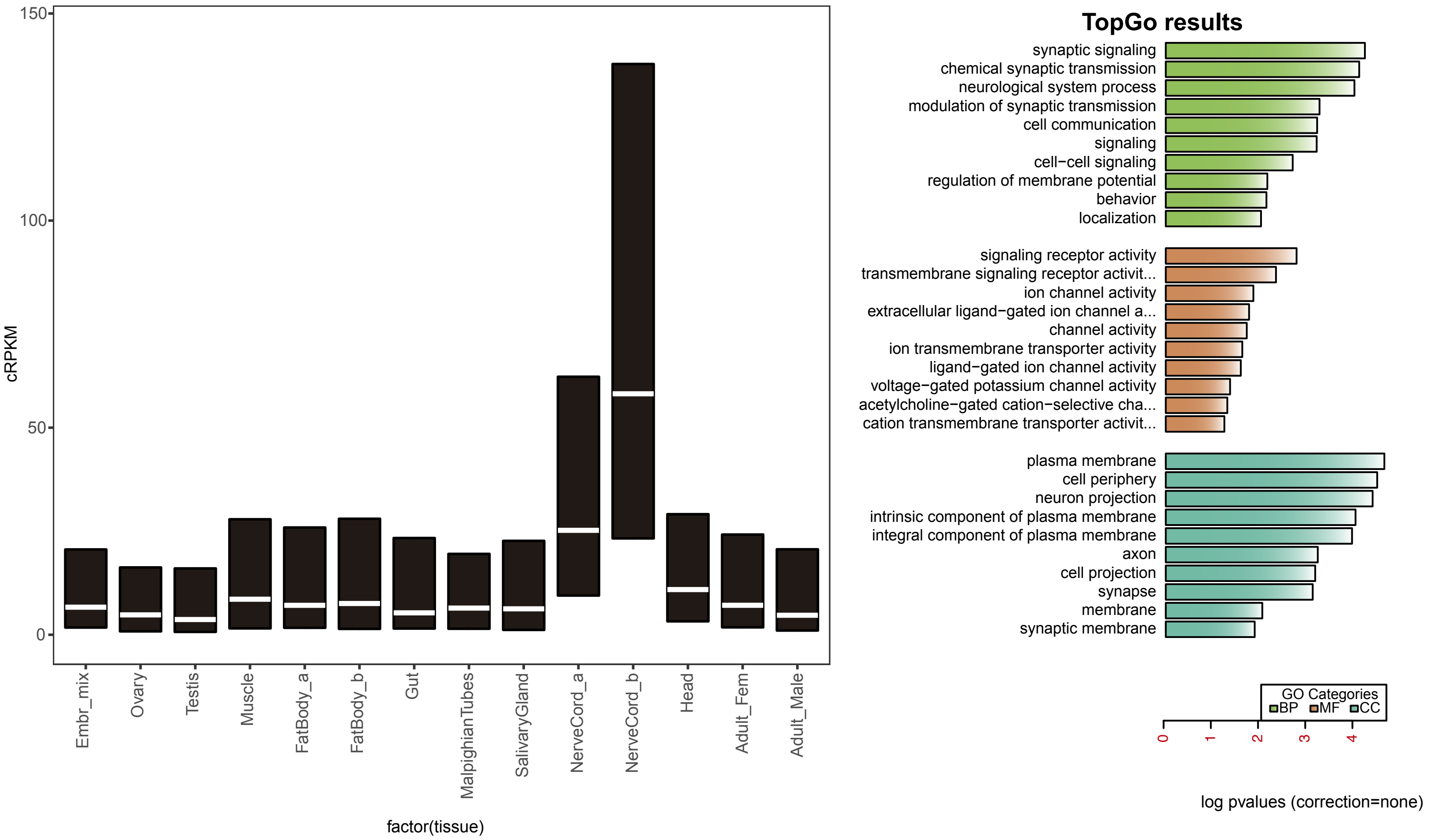

### *S. maritima* modules

#### Module:midnightblue (Splicing)

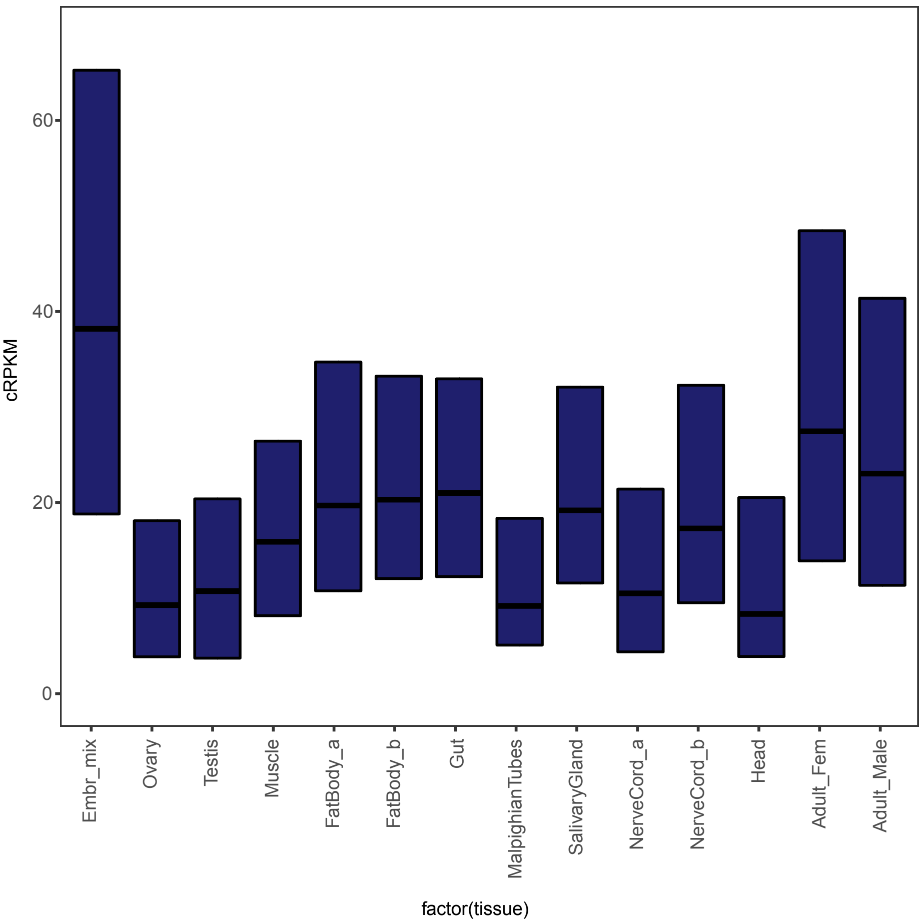


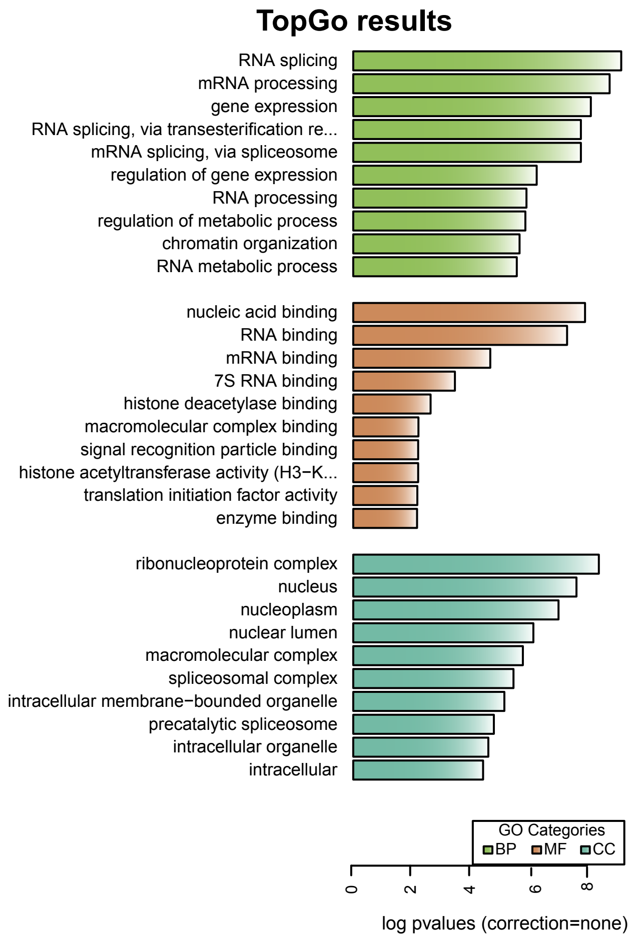


#### Module:blue (Gut)

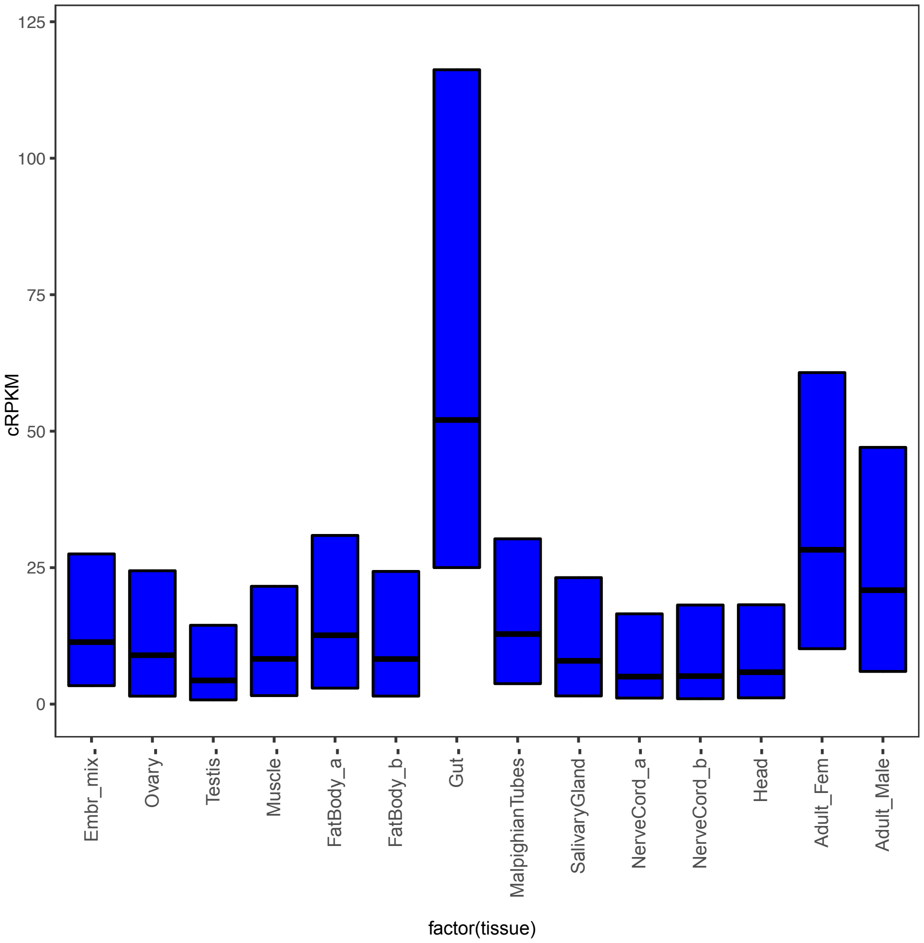


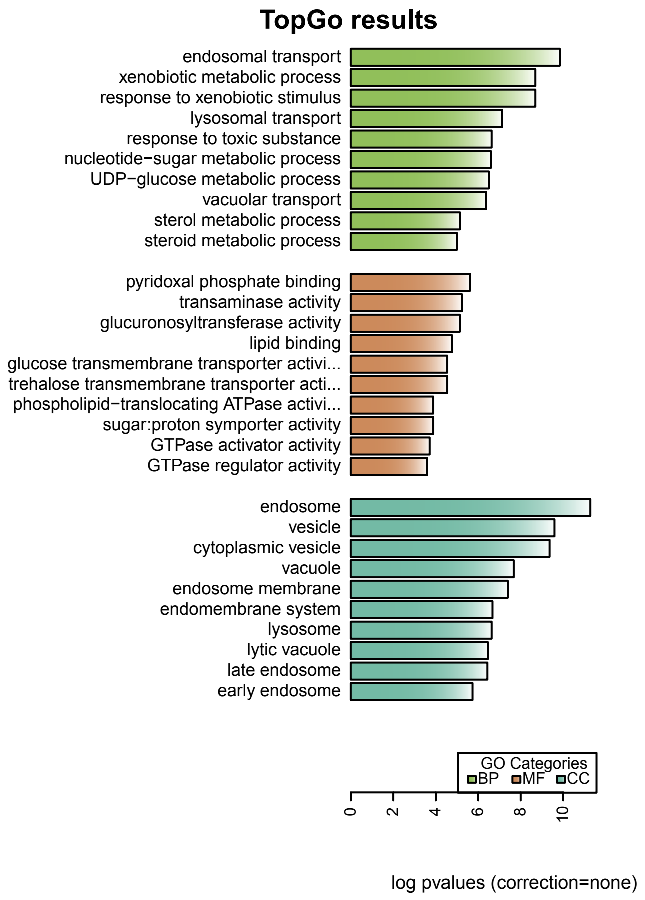

### *S. maritima* modules

#### Module:grey60 (Malpighian tubes)

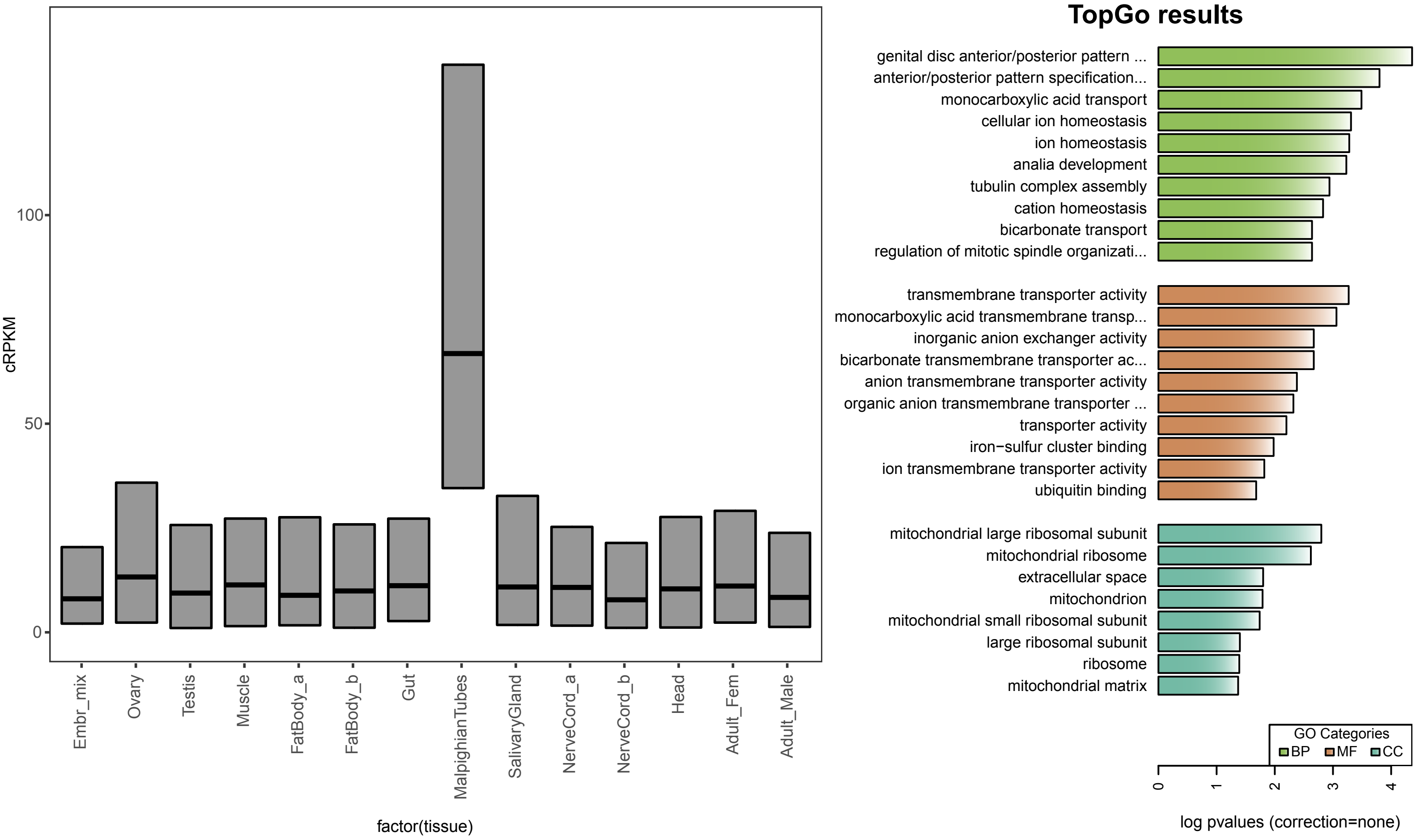


#### Module:purple (Fat body/Salivary gland- Translation)

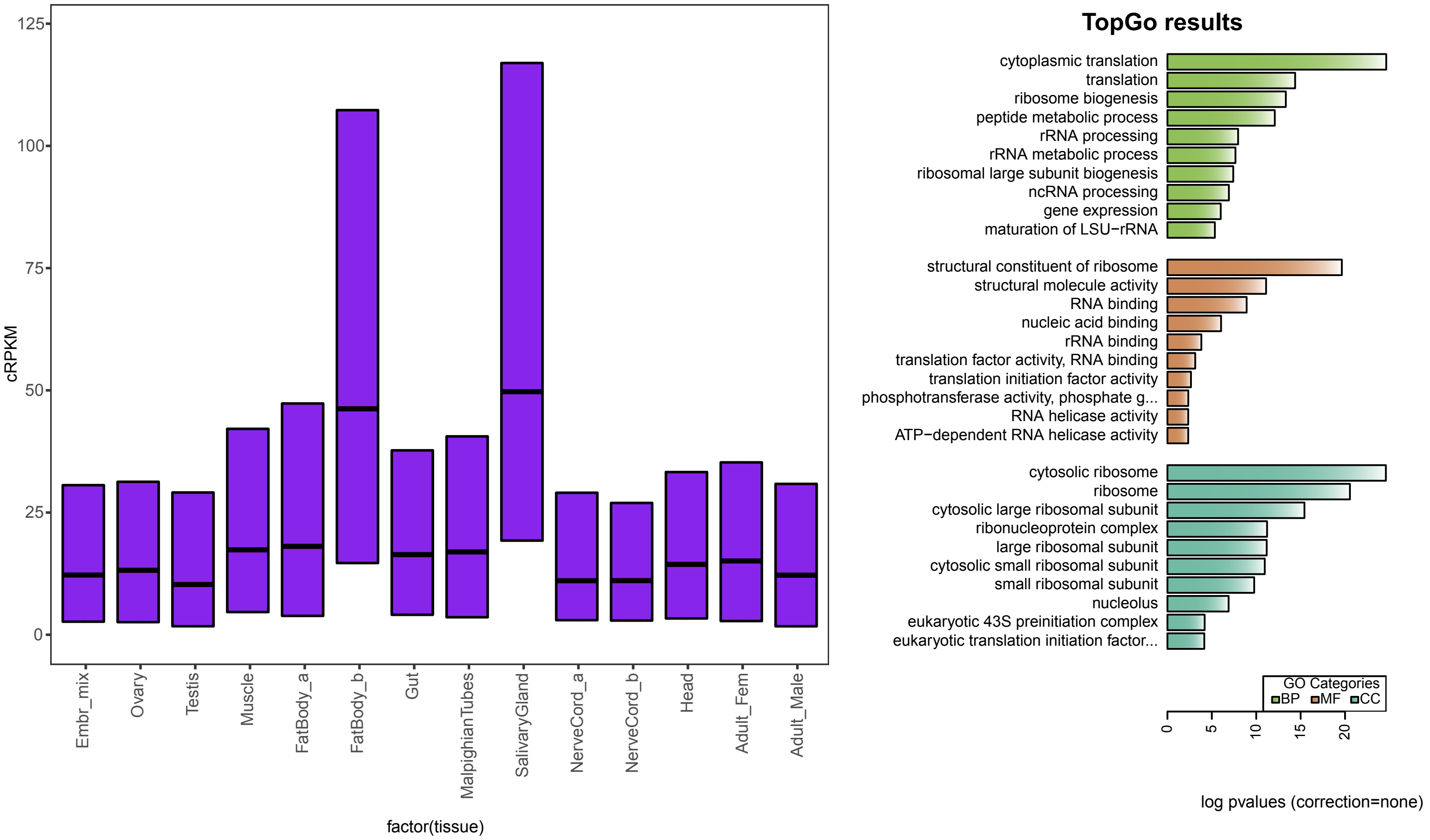

### *S. maritima* modules

#### Module:white (Proteasome)

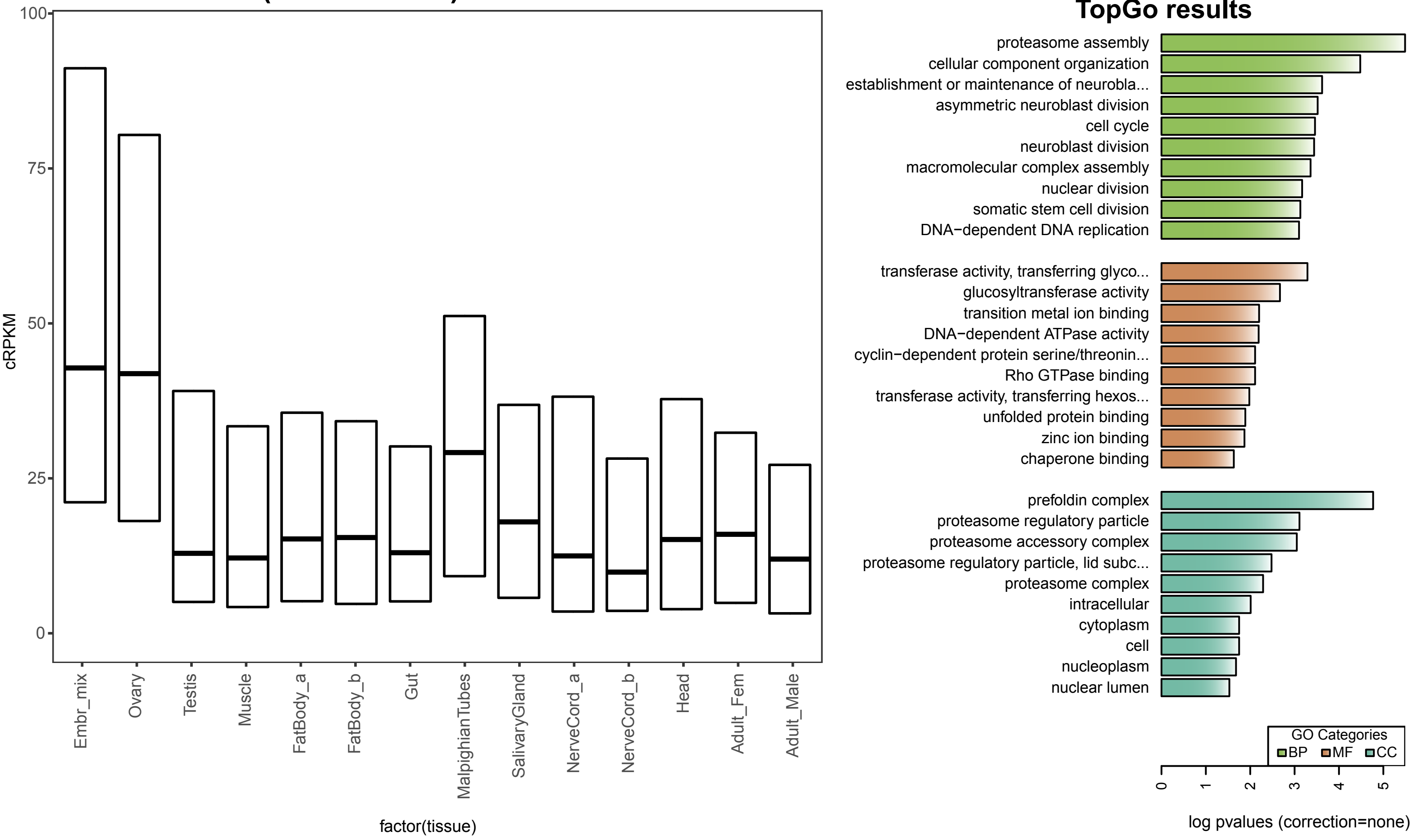


#### Module:lightgreen (Cellular respiration)

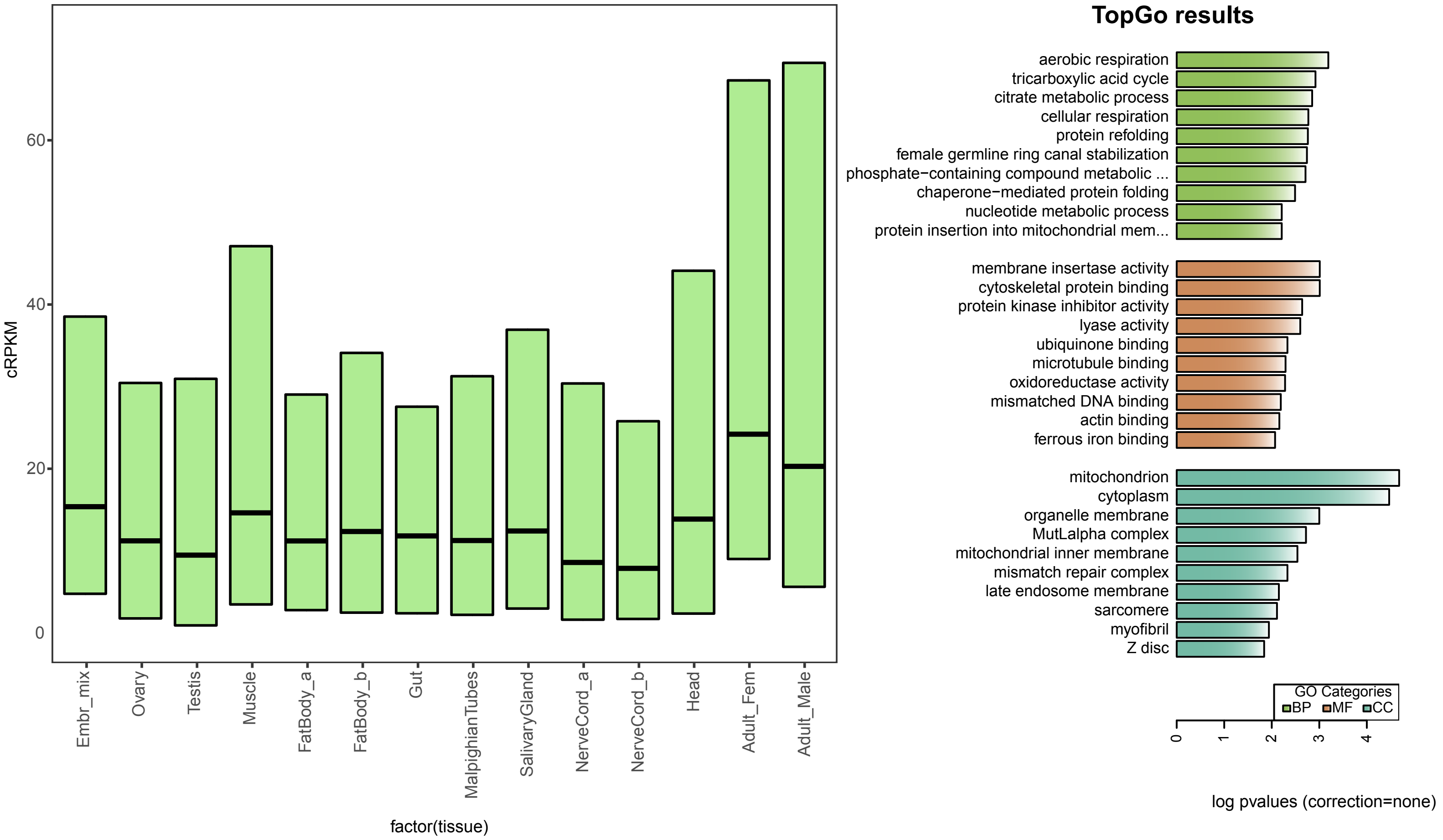

### *S. maritima* modules

#### Module:magenta (Ovaries- Replication)

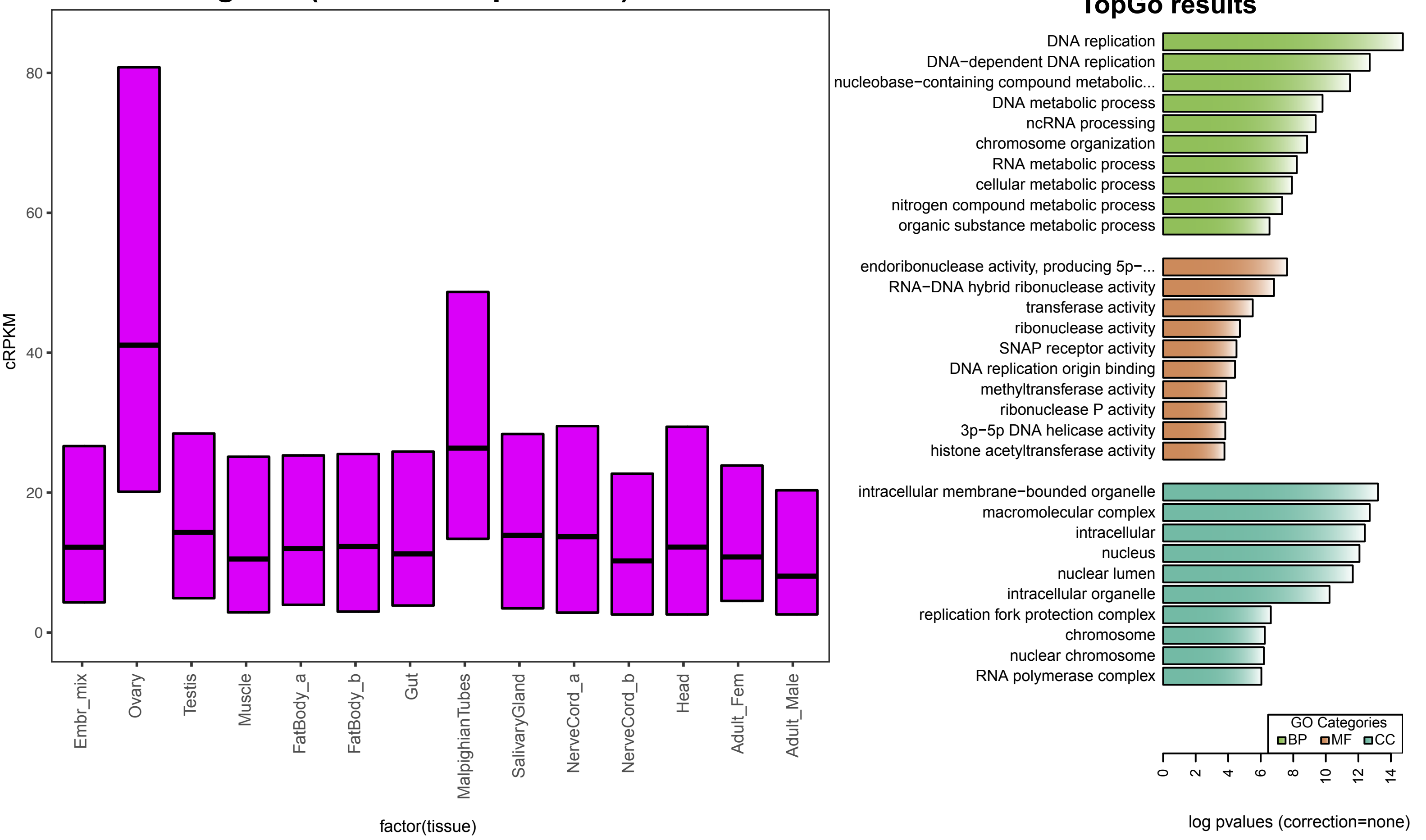


#### Module:lightyellow (Fatty acid metabolism)

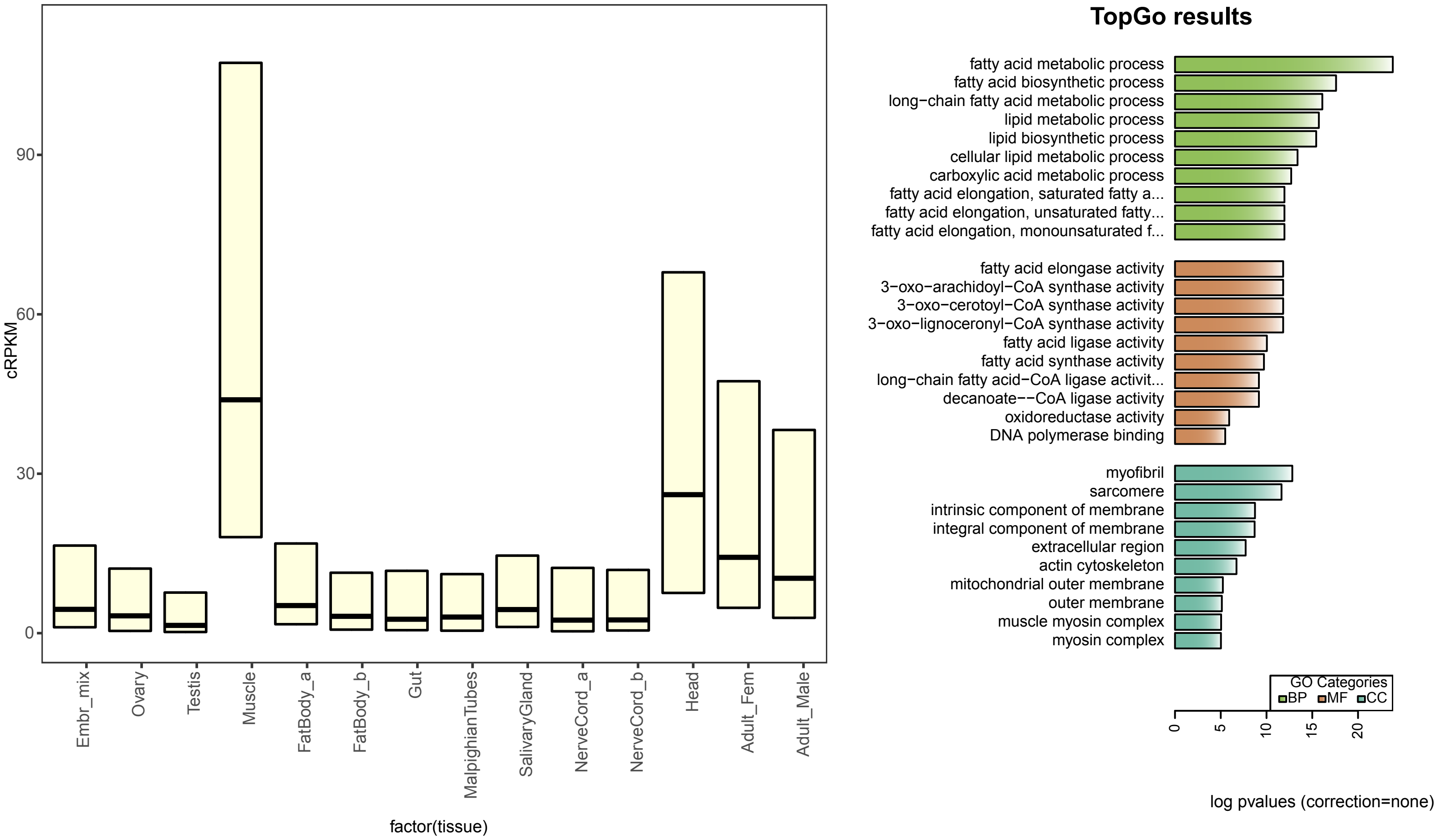

### *S. maritima* modules

#### Module:royalblue (Cellular metabolism)

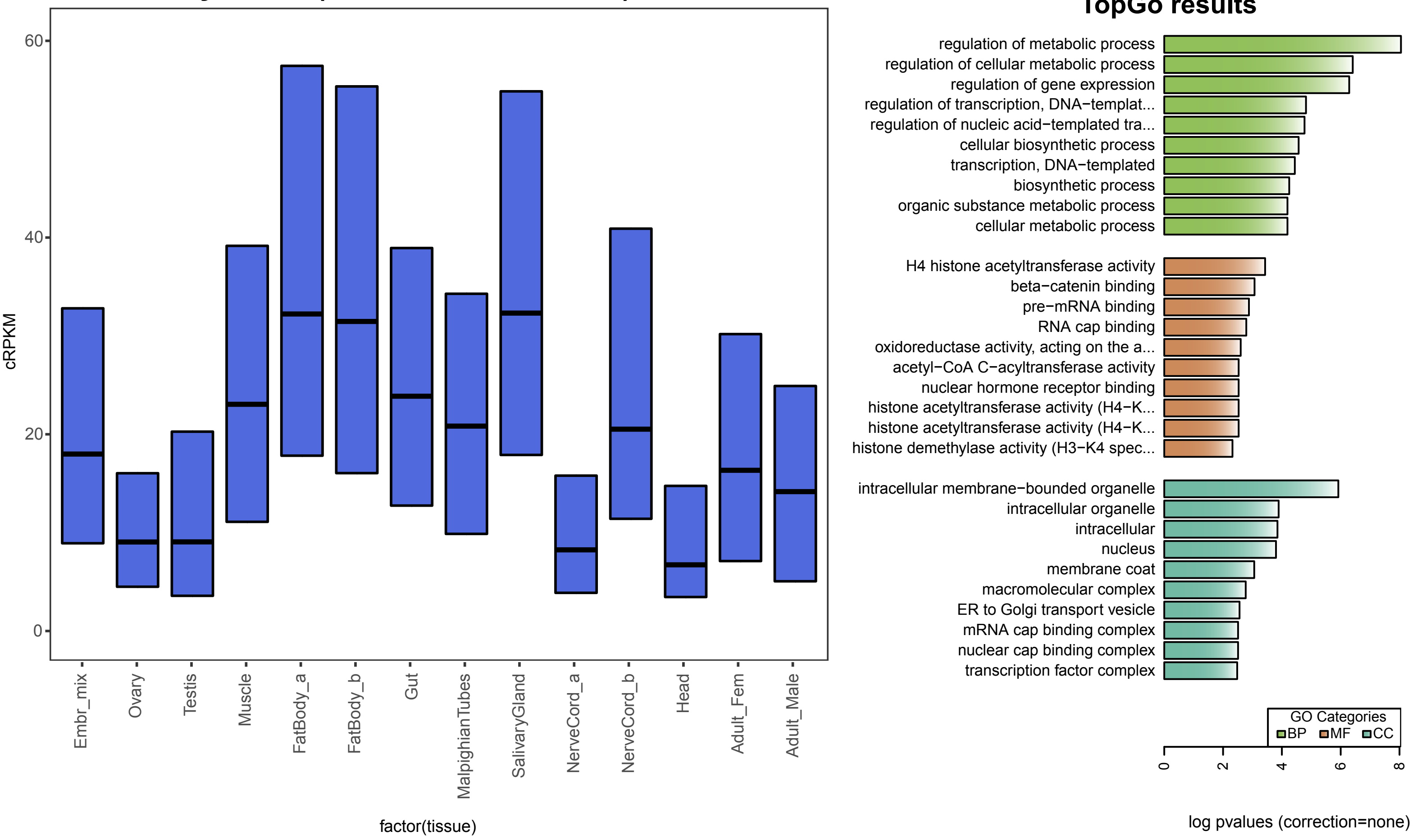


#### Module:darkorange (Fat Body- Post-translation modification)

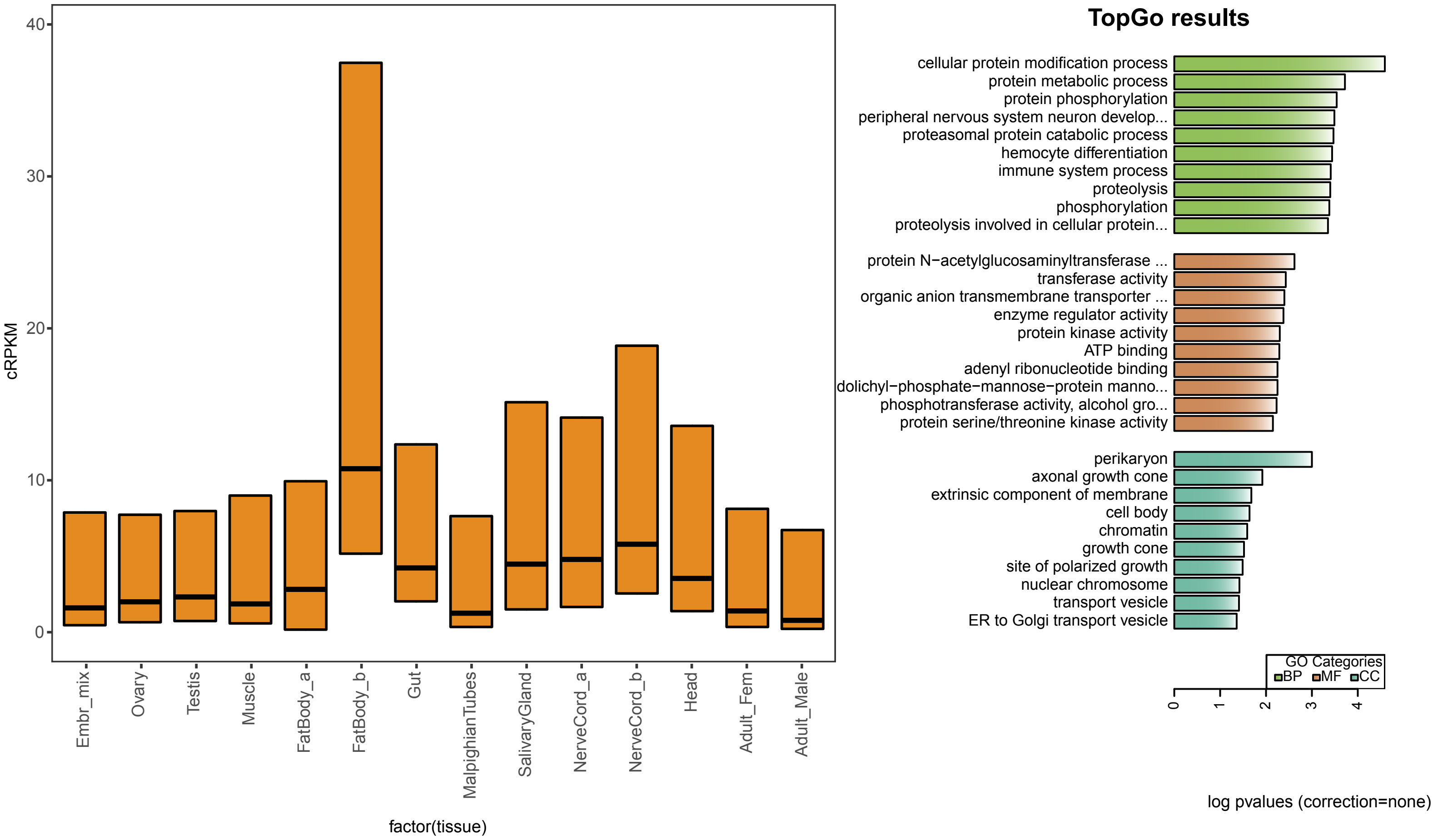

### *S. maritima* modules

#### Module:brown (Embryogenesis)

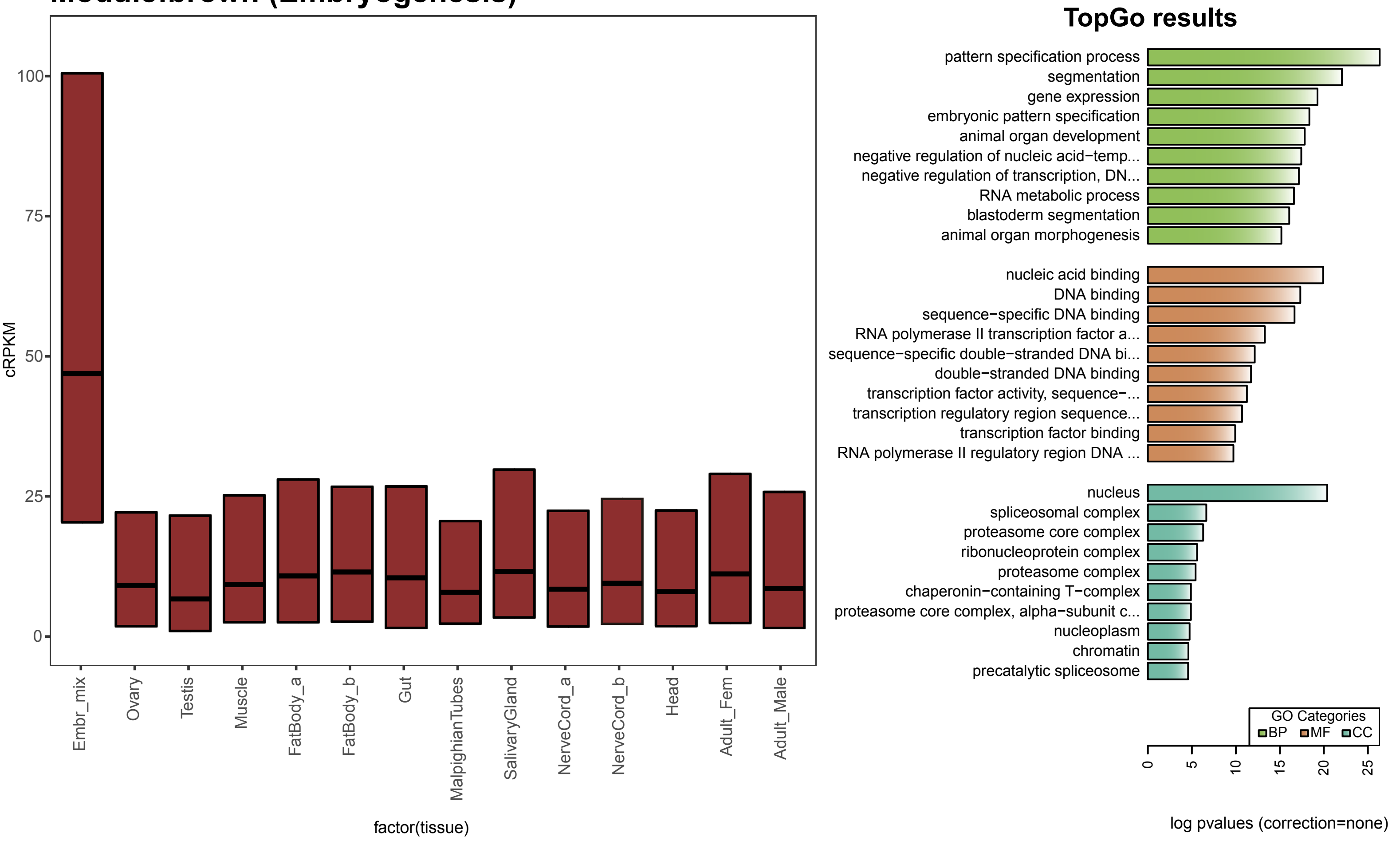


#### Module:turquoise (Testis- Cilium)

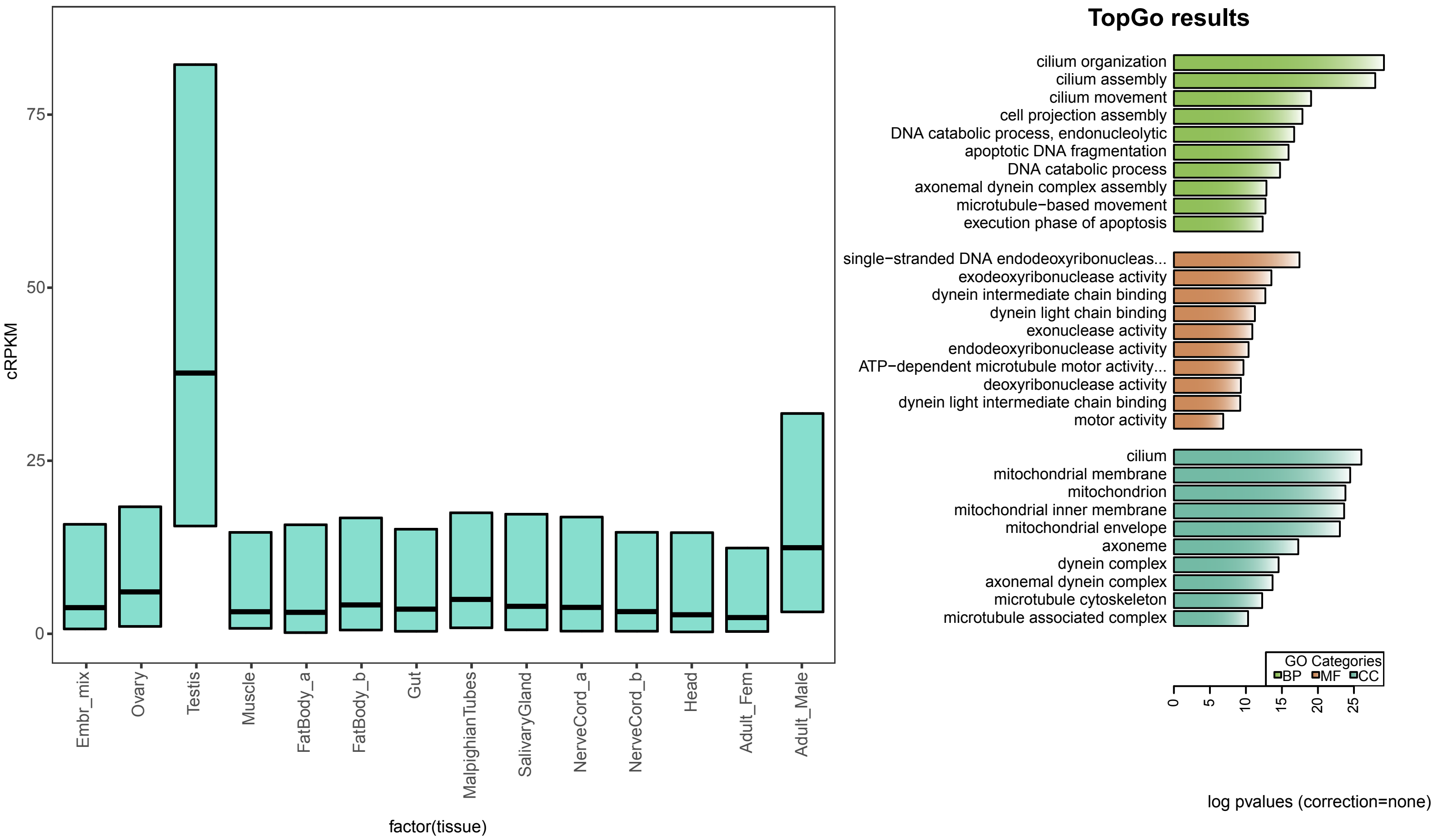

### *S. maritima* modules

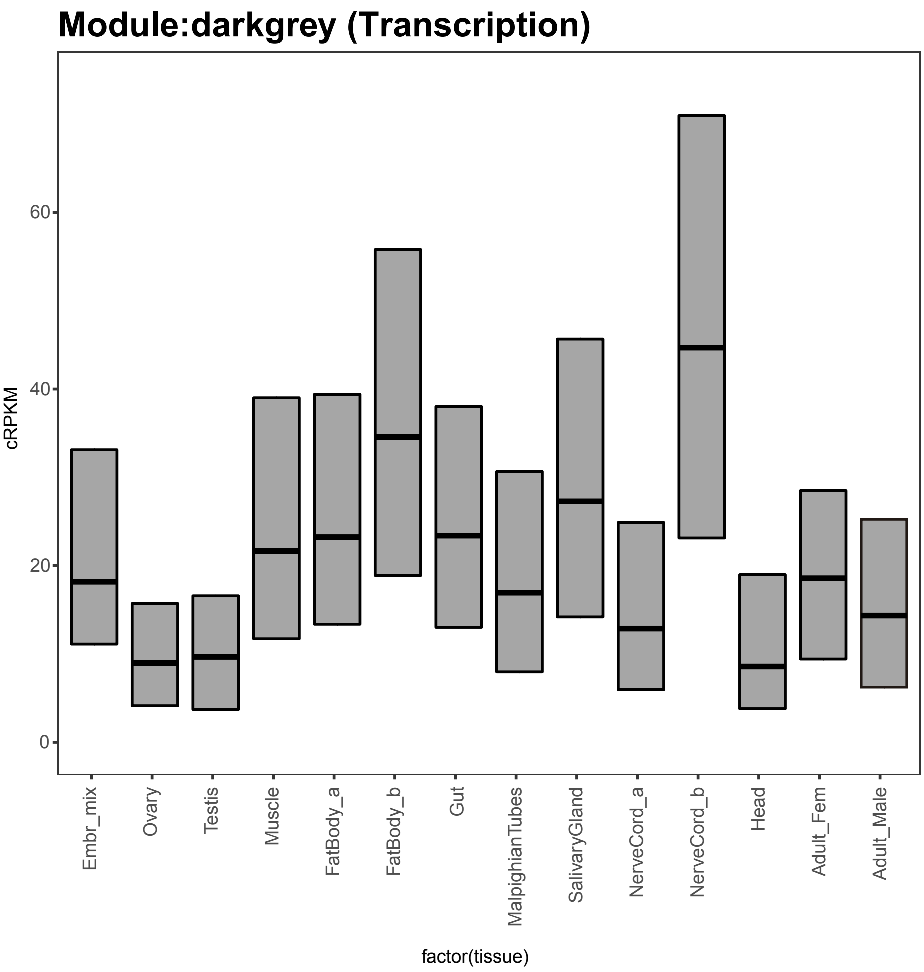


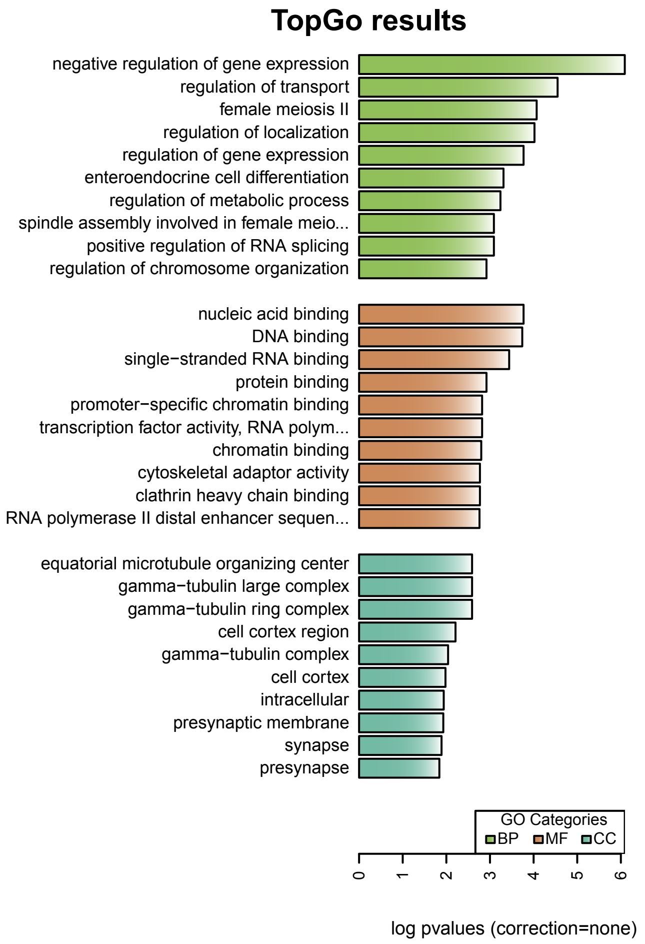
