## Supplementary information for "Genomic adaptations to aquatic and aerial life in mayflies and the origin of wings in insects"

**1. Genome sequencing and assembly**

1.1 Sample collection

1.2 Genomic DNA samples of *C. dipterum* for genome-wide sequencing

1.2.1 Illumina sequencing

1.2.2 Nanopore sequencing

1.3 Hybrid Assembly Strategy

1.4 RNA sequencing

1.3.1 PE RNA samples

1.3.2 SE RNA samples

1.5 Gene annotation

1.6 Repeat masking and TEs

1.7 Genome-wide alignment of Ephemeroptera and *D. melanogaster* genomes and PhastCons score calculation

**2. Resources for comparative genomics**

2.1 Gene family reconstruction and orthology assignment

**3. Comparative transcriptomics**

3.1 Transcriptomics clustering along life cycle stages (M-Fuzz)

3.2 Differentially gene expression analysis (DESeq2)

3.3 Weighted Gene Correlation Network Analysis (WGCNA)

**4. Chemosensory receptors**

4.1 Identification and annotation of chemosensory receptor complements

4.1.1 Functional and structural classification of CS sequences

4.2 Phylogenetic reconstruction

4.3 Gene Clustering analysis

4.4 Odorant Binding Proteins and Chemosensory Proteins

4.5 Gustatory and Odorant Receptors

4.6 Ionotropic Receptors

4.7 Sensory neuron membrane proteins

4.8 OBPs spatial gene expression patterns (in situ hybridizations)

**5. Opsin complement**

5.1 Identification and annotation of opsin genes

5.2 Gene trees

5.3 In hybridizations

**6. Phylostratrigraphy**

**7. Evolutionary origin of wings**

7.1 Functional test of shared genes

7.2 Tissue specific transcriptomics

### 1. Genome sequencing and assembly

#### 1.1 Sample collection

The samples were collected from an inbred line of *C. dipterum* kept in the laboratory for 7 generations [1].

#### 1.2 Genomic DNA samples of *C. dipterum* for genome-wide sequencing

##### 1.2.1 Illumina sequencing

DNA was extracted from 5 individual adult males using a standard Phenol:Chloroform protocol. The average amount of gDNA obtained per individual was 1 μg. The extracted DNA was used to build an Illumina sequencing library with fragment size 450bp. This library was sequenced using TruSeq Rapid SBS Kit v2 (Illumina Inc.), in paired end mode with a read length of 2x250bp and in two sequencing lanes of HiSeq2500 flowcell v2 (Illumina Inc.) according to standard Illumina operation procedures. A total of 160.89 Gb of raw sequence were produced. Primary data analysis, the image analysis, base calling and quality scoring of the run, was processed using the manufacturer's software Real Time Analysis and followed by generation of FASTQ sequence files by CASAVA. The FastQC reports suggested that quality decays dramatically towards the end of the read, specially affecting to some tiles during the last sequencing cycles. Noticeably, the average base quality scores are above 30 (1 error every 1000 base calls) in the first 180 bp of the reads.

##### 1.2.2 Nanopore sequencing

Five other samples, also of adult males from the same inbred line, were utilised to extract High-Molecular-Weight DNA, using MagAttract HMW DNA Kit (Qiagen) following manufacturers instructions, and build different library preparations for Oxford Nanopore Technologies (ONT) Sequencing. In total we performed 5 runs obtaining more than 7.56Gb of sequence. The ONT read length distribution comprises 25% of the reads being longer than 5.37 Kb, 50% longer than 3.51 and 95% longer than 1.64 Kb.

##### 1.2.3 PreQC Analyses

Before processing the reads, we estimated the genome size using three different methods. First an analysis of k-mers present in the sequence reads of the PE450 library was carried out using Jellyfish[2] to count k-mers of length 17. A peak k-mer depth was observed at 722-fold k-mer coverage (Supplementary Figure 1). A rough estimate of genome size can be made by dividing the total number of counted k-mers (150,314,958,593) by the k-mer coverage (722), which gives 208.19 Mb. Second, accounting for sequencing error, bias, and repetitive sequence using the program *gce*[3], we obtained a similar estimate more accurate of 208.35 Mb. Finally, SGA preqc [4] yielded a more conservative estimate of approximately 176.7Mb.

A detailed look at the 17-mer distribution indicated the presence of at least two contaminant genomes of different molarity in our sample. In order to detect such contaminants before assembling the mayfly genome, we ran Kraken [5] on the raw paired-end reads. This approach allowed us to detect and identify contaminant species in 4.21% of the reads (Supplementary Table 2).

1.2.4 Pre-processing the sequencing data

Due to the base quality distribution and the massive sequencing coverage of the PE library, we decided to trim the reads to 180bp and select those pairs with higher base quality along the read. Therefore, after trimming to 180 bp, we have used FASTX Toolkit 0.0.13 to keep those reads with a minimum base quality of 35 in at least 97% of their sequence. Finally, only valid pairs were kept for further filtering of contaminants.

Then, all reads were filtered by mapping (gem-mapper[6] with up to 2% mismatches) against a contamination database that included phiX, Univec sequences, *E. coli*, the complete mitochondrial genome of *Ephemera orientalis* and the genome of the contaminants detected with kraken [5] in the PE reads (Supplementary Table 3). These clean PE reads accounted for 55,502,519 2x180bp pairs, representing a sequencing coverage of 95.9x. The nanopore reads were also filtered using the same database, but adjusting the parameters to their higher error rate and lengths.

1.3 Hybrid Assembly Strategy

We ran a hybrid assembly using MaSuRCA v3.2.3 [7,8] with 95.9x short-read illumina and 36.34x ONT long-read coverage. All the options were set to default, but activating the usage of the linking mates and choosing the *Celera* assembler for contigging and scaffolding. The final assembly, *clodip*2, accounted for a total of 180,286,980 bp, had contig N50 461,411 bp and scaffold N50 434,950 bp. Regarding to gene completeness it is between 96.77% using CEGMA v2.5 [9] and 98.2% using BUSCO v3 [10].

1.4 RNA sequencing

A comprehensive collection of 35 RNA-seq datasets of multiple developmental stages and dissected tissues and organs was generated using the Illumina technology.

Four embryonic stages (E.4: 4 dpf or germ disc stage, E.6: 6 dpf or segmented embryo, E.10: 10 dpf or revolution stage, E.14: 14 dpf or pre-nymph), heads from nymphs at three stages: early, mid and late nymph, adult male heads, adult female heads, nymphal gills, nymphal wing pads, nymphal gut, nymphal malpighian tubes, adult muscle, testis, ovaries and adult brain samples were obtained from the *C. dipterum* inbred line kept in the laboratory. Organs and embryos were processed immediately after dissection and RNA was extracted using RNeasy Mini Kit (Qiagen) or RNAqueous™-Micro Total RNA Isolation Kit (Ambion) following manufacturers instructions.

1.4.1 PE RNA samples

Illumina libraries were constructed from RNA samples of four embryonic stages, adult male heads, adult female heads, nymphal gills, nymphal wing pads, adult muscle, testis and ovaries and sequenced at the CNAG-CRG using Illumina HiSeq2500 machine, the average of reads obtained was 50 million of 100-nt paired-end non-strand-specific reads. In addition, samples of gut (x2), malpighian tube (x2) and female brain (x2), were sequenced at the CRG Genomics facility on an Illumina HiSeq2500 machine generating an average of 50 million 100-nt paired ended strand specific reads.

1.4.2 SE RNA samples

RNA obtained from the heads of three nymphal stages was used to generate Illumina libraries through a standard Illumina TruSeq protocol. We obtained an average of 10 million reads of 50-nt single end non-strand specific.

1.5 Gene annotation

Reads from all transcriptomes were assembled altogether using Trinity [11] and subsequently aligned to the genome using the PASA pipeline [12]. After conceptual translation, high-quality transcripts were selected as a training set to build a hidden-markov profile for *de novo* gene prediction in Augustus [13].

Reads were aligned to the genome using STAR with default parameters [14] and transcriptome assembled using Stringtie for each sample [15]. The assemblies derived from all samples were merged using Taco which ensures high accuracy fusion of transcriptomes [16]. This transcriptome assembly gather 31913 gene loci and 56639 transcripts. In parallel, alignments from all samples were merged in order to filter and summarise splice-junctions using the Portcullis tool (https://github.com/maplesond/portcullis).

Consensus transcriptome assembly and splice-junctions were converted into hints and provided to Augustus gene prediction tool which yielded 16364 evidence-based gene models. Isoforms and UTR regions were added by updating Augustus gene models against the PASA transcriptome databased for two subsequent rounds, resulting in 34699 transcripts for the 16364 genes. These models contain 4308 non-redundant PFAM models as assigned using the PfamScan tool.

1.6 Repeat masking and TEs

RepeatModeler, a *de novo* repeat identification program, version 1.0.11 [17] was run on the *C. dipterum* genome with default parameters. The consensus sequences reported by RepeatModeler which had not been properly classified (thus assigned to the Unknown class) were filtered using BLASTX from the BLAST+ 2.9.0 release against the nr database (downloaded using the update_blastdb.pl script on the 2019-07-30). Only elements whose hit title contained at least one element from a set of TE-related keywords (transpos, integras, retro, among others) were conserved. The consensus were added to the RepBase library [18] for Metazoa from the 20181026 update and this new library was used to mask the genome with RepeatMasker [19] version 4.0.9. The divergence distribution obtained by calculating the Kimura distance from each TE classified up to the family level to the consensus of its family. The larger number of differences between the identified copy and its consensus translates into a larger Kimura distance. The distribution was plotted (Figure 1B) using the helper scripts from RepeatMasker. Unclassified sequences were not included in this plot.

1.7 Genome-wide alignment of Ephemeroptera and *D. melanogaster* genomes and PhastCons score calculation

Genomes from *E. danica* and *D. melanogaster* were aligned to our *C. dipterum* genome assembly using lastz (--inner=2000 --ydrop=3400 --gappedthresh=6000 --hspthresh=2200). We processed the alignment according to UCSC guidelines [20] using the Kent utilities. We then used axtChain to transform pairwise Axt alignments to chain alignments, that were sorted and filtered with chainMergeSort and chainPreNet. Afterwards, we converted alignment chains to 'nets' with chainNet. A combination of the pairwise alignments into a single multiple alignment in *C. dipterum* coordinates was performed using multiz-tba. Finally, we calculated the background model from the longest scaffold using phyloFit [21] and the conservation scores and highly conserved regions were computed with PhastCons [22]

### 2. Resources for comparative genomics

2.1 Gene family reconstruction and orthology assignment

To obtain phylogeny-based orthology relationships between different taxa, the predicted proteomes of 14 species (*Apis mellifera, Cloeon dipterum, Daphnia pulex, Drosophila melanogaster, Ephemera danica, Folsomia candida, Homo sapiens, Ladona fulva, Pediculus humanus, Stegodyphus mimosarum, Strigamia maritima, Tigriopus californicus, Tribolium castaneum, Zootermopsis nevadensis*) representing major arthropod lineages and outgroups were used as input for OrthoFinder2 [23]. All versus all similarity searches were obtained using DIAMOND (0.9.15)[24] and an inflation parameter of 1.5 for the clustering. Then trees were computed using MAFFT(v7.221)[25] to align of all members of each orthogroup (LINSI mode), trimAl (v1.4)[26] to trim non-homologous regions (-gt 0.2), and RAxML-NG [27] for maximum likelihood phylogenetic reconstruction (--model LG+G4 --seed 12345) and parsed with OrthoFinder 2.

Gene gains and losses in major transition of arthropods evolution were inferred with a recently presented pipeline [28]. Similarity searches of protein coding genes were obtained using DIAMOND (0.9.25)[24]. Clustering was performed with MCL v14-137 with a granulosity parameter of 2 [29]. The patterns of gains and losses were reconstructed based on the taxonomic occupancy of the gene clusters (Supplementary Figure 1). The *C. dipterum* Pfam and GO annotations were used to inspect the functions of these genes.

### 3. Comparative transcriptomics

#### 3.1 Transcriptomics clustering along life cycle stages (M-Fuzz)

Mfuzz software [30] was used to perform soft clustering of genes according to developmental and life history expression dynamics in *C. dipterum.* We selected eight developmental and post-embryonic stages: E.4 (4 dpf embryos or germ disc stage), E.6 (6 dpf embryos or segmented embryo), E.10 (revolution stage embryos), E.14 (pre-nymph), early nymph heads (2.5 mm nymphs), mid nymph heads (4 mm nymphs), late nymph heads (5 mm nymphs) and adult heads. To be able to profile the eight RNA-seq datasets together (PE and SE libraries), we used "R1" datasets from the PE libraries. Datasets used were an average of the replicas for each time point to avoid experimental or sex bias. Datasets were normalised using "DESeq2" library and genes with low variability (coefficient of var < 3) across the datasets were removed (1475 genes filtered out) for the analysis with default parameters. Minimum centroid distance was calculated using "Dmin" function to obtain the optimum number of clusters, which resulted in 30 (Supplementary Figure 2).

Taking the *Drosophila melanogaster* orthologs, we performed Gene Ontology (GO) enrichment analysis for each of the clusters using DAVID [31]. These analyses allowed us to assign GO terms that reflected main biological and molecular processes occurring for each of the clusters. For instance, cluster 2, 3, 8, 21 or 27, which included genes with high expression during embryonic stages, showed enrichment in terms such splicing, dorsal closure, axon guidance, etc. associated to morphogenetic processes happening at these developmental points. By contrast, clusters whose genes are expressed during nymphal stages, such as cluster 1, cluster 7, cluster 9 or cluster 18 exhibited enrichment in categories related to chitin, cuticle formation and perception of chemical stimulus (Supplementary Figure 2, Supplementary Table 8).

#### 3.2 Differentially gene expression analysis (DESeq2)

We mapped adult male and adult female head RNA samples to the annotated transcriptome using hisat2 v2.1.0 [32]. Then, we obtained counted reads for each of the samples (two replicas each condition) with samtools utilities (<http://www.htslib.org/>) and htseq-count [33]. Differentially gene expression analysis was performed with "DESeq2" package in R [34] to detect whether there were differences in gene expression based on sex.

We obtained 3243 genes that were differentially expressed between male and female heads (padj < 0.05), and of them, 1406 genes were upregulated in female heads and 1837 were upregulated in male heads (Supplementary Table 10).

#### 3.3 Weighted Gene Correlation Network Analysis (WGCNA)

We used the cRPKM metric (corrected-for-mappability Reads Per Kilobasepair of uniquely mappable positions per Million mapped reads [35] to perform these gene expression analyses. We selected the transcript with largest number of exons for each of the protein coding genes annotated in *C. dipterum* genome. In the case of *D. melanogaster* and *S. maritima* we used 19 and 14 datasets from [36] (Supplementary Table 11) and calculated cRPKMs as described in [35].

To characterise modules of co-expressed genes across our RNAseq datasets that include developmental stages and nymphal and adult tissues and organs, we utilised WGCNA [37]. We selected 26, 19 and 14 samples of *C. dipterum*, *D. melanogaster* and *S. maritima* (Supplementary Table 11). We performed the analyses using as datasets genes that were present in a least two species in our family reconstructions and showed variance across samples (coef. var ≥ 1). In total, the number of genes included in each dataset were, 13720 in *C. dipterum,* 12463 in *D. melanogaster* and 13001 in *S. maritima* (Supplementary Table 11). After running WGCNA software with default parameters, we obtained 22, 17 and 21 modules for *C. dipterum, D. melanogaster* and *S. maritima*. Each module was designated with a tissue or a biological category related to gene expression across tissues and GO enrichment (Supplementary Data 1). Finally, we analysed the overlap between homologous groups for each pair of modules for each of the species in a pairwise manner. To evaluate the significance, we performed hypergeometric tests.

### 4. Chemosensory receptors

#### 4.1 Identification and annotation of chemosensory receptor complements

We created a dataset containing reference sequences for each chemosensory (CS) gene family (GR, OR, IR/iGluR, CD36/SNMP, OBP and CSP) from relative annotated insect genomes (Supplementary Figure 3[38-47]). In addition, we constructed specific HMM profiles for each CS gene family based on their Pfam profiles (see Supplementary table 1 in [48]).

We used the sequence database and the HMM profiles in the program BITACORA [49] to (i) identify new CS members, or to curate the already annotated ones, among the pre-compiled gene models of these families (obtained with automatic methods), and (ii) to generate new models (of previously undetected copies) from the genomic sequences. Briefly, we performed various iterative rounds of BLASTP and HMMER searches against the automatically annotated proteins of *C. dipterum and* curate incorrect and incomplete (when possible) gene models. Also, we used TBLASTN against the genomic sequence to identify novel (not annotated by the automatic methods) regions encoding CS proteins. We generated a GTF file containing our curated annotation of *C. dipterum* CS genes.

##### 4.1.1 Functional and structural classification of CS sequences

We classified the novel sequences in different categories based on structural and functional criteria [50]. First, we examined the presence of premature stop codons; these features could represent real non-functional copies (pseudogenes), errors in sequencing or genome assembly steps or inaccuracies in our automatic annotation step based on TBLASTN hits. All sequences encoding complete proteins (CPs) that were free of stop codons were included in the first category (CP set). Operationally, we considered a CP when its length was >80% of the corresponding average protein domain length. In addition, for the GR and OR families, we also required that the CP members contained a minimum of 5 of the 7 transmembrane domains (defined by the software TMHMM version 2.0c [51]; Phobius version 1.01[52]). For the CP IR/iGluR members, we required the presence of the ligand channel domains, namely, PF00060 (ligand-gated ion channel), present in all IR/iGluR subfamilies, i.e., kainate, AMPA, NMDA, conserved IRs (IR25a/IR8a), and divergent IRs [53]. The remaining sequences that were free of stop codons and did not pass the length filter criteria were classified as incomplete proteins (IP set). Finally, the CP and IP sequences exhibiting some in-frame stop codons (that could represent pseudogenes, among other features; Ψ) were incorporated into the Ψ data set. We renamed all proteins with the initials of the family name and a number starting from 100, except those with clear homology to *D. melanogaster* CS members which were named as in *Drosophila*. We added a p to the name of incomplete proteins to identify them.

We identified a total of 367 putative proteins across all CS gene families in the genome of *C. dipterum.* CS gene families are characterized as fast evolving and highly divergent both across insects and family and, therefore surprisingly, the 95.4% of the copies were already predicted in the automatic annotation, although some of them had incorrect gene models (i.e. extra exons badly assigned to the gene, missing exons…), while the remaining were identified from genome sequence. The use of deep transcriptome libraries from several stages and tissues in genome-wide automated gene modelling provided these gene models for most of the CS sequences.

#### 4.2 Phylogenetic reconstruction

We included in the phylogenetic analyses all CS genes identified in *C. dipterum*, the members of these families in the fruit fly *D. melanogaster*, and the GR, OR, IR and OBP proteins annotated in the *C. splendens* genome (Ioannidis, Simao et al. 2017). Moreover, we also included the OR annotated in *E.danica* genome [47] and the OBPs obtained from a preliminary sequence similarity-based search in *E.danica* genome assembly (https://i5k.nal.usda.gov/Ephemera_danica). The analysis of the OBPs and CSPs was performed with the mature proteins (after excluding the signal peptide identified using SignalP software; [54]). We used maft (with '--auto' option) to build family-specific multiple sequence alignments (MSA) [55]. The phylogenetic analysis was performed with IQ-TREE version 1.6.5, estimating automatically the protein substitution model for each tree [56]. Node support was estimated from 1000 ultrafast bootstrap replicates [57]. The phylogenetic tree images were created using the iTOL webserver [58]. Trees were rooted using outgroups according to available phylogenetic information. Additionally, we used OrthoMCL (v2.0.9; [59]) to infer orthology among the gene family members to compare with the relationships observed in the phylogeny.

#### 4.3 Gene Clustering Analysis

We used computer simulations to test whether the members of a CS gene family are physically clustered in the genome of *C. dipterum*. We assessed the clustering pattern of all annotated CS genes by computing the variance of the number of copies of a given family per scaffold (in 1,395 scaffolds). To build the null hypothesis of no clustering, we randomly choose from the total genome gene set a number of genes equal to the number of copies of the gene family we wanted to test, and calculated the variance of the number of genes per scaffold. We repeated this procedure 10,000 times until having a empirical distribution of this quantity based on all genes. The empirical *P*-value of being distributed across the genome just like any other gene, regardless of being a member of a CS family, was obtained by comparing the observed variance of CS genes per scaffold against this genome-wide distribution using the Empirical Cumulative Distribution Function (ecdf) in R (R Core Team 2016).

#### 4.4 Odorant Binding Proteins and Chemosensory Proteins

We found the largest OBP repertoire known in insects, with a total of 191 fragments encoding members of this family, being 167 of them complete and conserving the typical conserved cysteine pattern. Among the 24 partial OBP sequences, two are putative pseudogenes (sequences containing premature stop codons that truncate the sequence) and 22 incomplete OBP that could be either also pseudogenized sequences or just incomplete annotations or assemblies. The *C. dipterum* OBP sub genome far exceeds the 109 OBP genes identified in the cockroach *Blatella germanica* [60], and contrast greatly with the four OBP found in the genome of closely related species *Calopteryx splendens* [42], and the 39 putative genes of this family found in a preliminary search in *E. danica* genome.

We also identified 16 CSP proteins with complete gene models and the typical four cysteine pattern of this family. The phylogenetic tree shows that this genome encodes three highly divergent members of this family (CdipCSP107, CdipCSP108 and CdipCSP113; Supplementary Figure 2b). We could not establish any orthology relationship between *D. melanogaster* and *C. dipterum* CSP.

We determined that the members of the OBP family of *C. dipterum* are significantly clustered in the genome (*P*-value < 0.0001). This clustering pattern would be expected if new copies originate by tandem gene duplications and their genomic structure is maintained for long time periods by selection against rearrangements, as reported for this family in different insect species [61-63]. An alternative, not selective, explanation is a very recent burst of gene duplication in the lineage of *C. dipterum* (in the family or in the entire genome) that generated large tandem arrays of genes that still remain physically close in the genome. We observed several species-specific phylogenetic clades of these physically close copies in the OBP family tree with relatively short branch lengths, suggesting a very recent origin and, therefore supporting the second hypothesis; nevertheless, we also observe some basal lineages that group with *D. melanogaster* and *C. splendens* sequences, pointing to a much older origin and, therefore, suggesting some selective constraint against cluster rearrangements (Figure 2a).

The OBP tree also highlights the presence in *C. dipterum* of an ortholog of the DmelOBP73a protein, here named as CdipOBP73a, which is also present in *C. splendens*, CsplOBP1. This orthogroup is highly conserved across insects, indicating a critical (but already unknown) function in this group [41]. Additional orthologies, such as that of the OBP43a group, were also detected, although bootstrap support was low.

4.5 Gustatory and Odorant Receptors

The genome annotation of chemoreceptors was more challenging as they are 4-fold longer than OBPs and CSPs and their expression is more restricted (most of them are specific to chemosensory appendages). We found that almost half of the sequences encoding chemoreceptors are partial fragments, including also some putative pseudogenes. Nonetheless, we identified 64 GR proteins which would encode, at least, 56 receptors, being 26 of them complete. Regarding OR, we found 50 sequences encoding at least 43 proteins, 29 with full-length models. Interestingly, and unlike most insects, in *C. dipterum* the number of OR and OBP proteins, both involved in olfaction, is very different, mainly due to the large number of extracellular soluble proteins.

The phylogenetic analysis of chemoreceptors uncovered the presence in *C. dipterum* of the highly conserved OR co-receptor, known as ORCO and of 49 specific OR (Supplementary Figure 3c and f). The OR copies of the mayfly seem to be old and, given that the number of OR annotated in the other Ephemeroptera, *E. Danica*, is quite similar (46; [47]), an expansion of this family in the ancestor of these two lineages would be the most plausible hypothesis for the observed differences between them and other sister lineages.

For the GR, we found a copy similar to the *Drosophila* receptors for sugar taste (here CdipGR124), a group of proteins conserved in all insects and also found in the crustacean *D. pulex* [64]. Two additional GR found in this study, CdipGR133 and CdipGR156, are similar to the GR characterized as the $CO_2$ receptors in *Drosophila*. Apart from these highly conserved sequences, we identified some phylogenetic-based candidates to be the fructose receptor or to participate in bitter detection, although the key nodes were largely unsupported. The remaining GR grouped in species-specific clades and its function is completely unknown. Noticeably several of these GR are expressed in the pre-nymphal embryo and in nymphal heads, so these receptors probably also mediate gustation in *C. dipterum*.

4.6 Ionotropic Receptors

Of the 34 IR/iGluR genes identified in *C. dipterum* (26 complete sequences), 11 are canonical iGluRs, while 23 likely encode ionotropic (putative chemosensory) receptors (Supplementary Figure 3). Particularly, we identified some IR members known to be conserved across insects, such as the co-receptor IR8a, which is duplicated in the mayfly ( CdipIR8a_1 and CdipIR8a_2), IR25a and IR76b,IR93a, IR21a, IR40a, IR68a, the three first associated in thermo- and hygrosensory mechanisms [65]. Among the IR members identified in *C. splendens*, only IR75c, which respond to various acids (Prieto-Godino, Rytz et al. 2017) seems to be missing in *C. dipterum*. The phylogenetic analysis also uncovers a small burst of ancient CdipIRs similar in number to the observed in *C. splendens* (Supplementary Figure 3e). These divergent IRs could be implicated in chemosensory functions but also in the response to other stimuli, such as temperature or humidity, as has been documented in *D. melanogaster* [65]. The diversity of tissues in which these receptors are expressed predicts their broad range of functions. We observed particular IR copies highly expressed in adult heads (IR40a and IR93a), gills (IR68a, IR107, IR8a_1, IR103 and IR114p) and early embryos (E4; IR109, IR112p, IR25a), while canonical iGluRs were found majorly expressed in whole heads and brains.

4.7 Sensory neuron membrane proteins

We identified 12 complete genes encoding CD36-SNMP proteins, and no evidence of partial fragments or pseudogenes in this family. Only SNMP, found to be expressed in specific *Drosophila* pheromone-responding sensory neurons, has been related with chemoreception, playing a key role in the sensory perception of an important fruit fly pheromone by facilitating the contact between the ligand and the membrane receptor [66]. The phylogenetic analysis indicate that the only copy of this family conserved in *C. dipterum* is SNMP2, which is represented by five divergent paralogs. Interestingly, four of these five genes are mostly expressed during embryonic stages, in particular at the segmented embryo stage (E6).

4.8 OBPs spatial gene expression patterns (in situ hybridizations)

We selected some of the OBP genes that showed high expression in the gills RNA-seq datasets to investigate the spatial expression pattern within this organ. We designed and generated RNA probes against *OBP199, OBP260* and *OBP219* (Supplementary Table 14). After whole trunks were fixed in FA 4% at 4º C o.n., gills were separated from the body wall and cuticle was carefully removed. Gills were incubated at 60º C with RNA probes after post-fixation in Methanol and Proteinase K treatment. Anti-digoxinenin-AP (*OBP260*, for 1 h 30 min; Roche) or anti-digoxinenin-POD (*OBP199, OBP219,* o.n. at 4º C; Roche) antibodies were used to detect our DIG-labelled probes. *OBP199* and *OBP219* gills were incubated with Tyramide Signal Amplification (TSA) 1:100 in borate buffer for 1 hour. After washing TSA reaction, gills were stained with DAPI

1:10000 and Goat anti-HRP-Cy3 1:100 (Jackson Immunoresearch) at 4º C, o.n. Image acquisition was done with Leica SPE confocal microscope. Images were processed with Fiji [67]. For *OBP260* in situ hybridization standard protocol was followed, NBT/BCIP was used as chromogenic reagent.

### 5. Opsin complement

5.1 Identification and annotation of opsin genes

A total 1247 opsins from all the major metazoans groups were used as seed in the BLAST research the predicted protein sequences of *Cloeon dipterum, Ephemera danica* (https://www.hgsc.bcm.edu/arthropods/mayfly-genome-project) *and Ladona fulva* (https://www.hgsc.bcm.edu/arthropods/scarce-chaser-genome-project). All the genes with e-value < $10^{-10}$ were retained for further analysis. To be considered an opsin it was required that to have the retinal binding domain or that (for shorted sequences) the first BLAST again Uniprot was an opsin. These sequences were then merged with those from [28,68,69]. To this set of sequences, additional mayfly LWS, UV and Blue opsin sequences from transcriptome assemblies for *Baetis sp. EP001* and *Epeorus sp. EP006* were obtained from [68,69] and for *Baetis sp. AD2013* from [70]. Sequences derived from the previously mentioned transcriptomes that were shorter than 100 aas were not included in subsequent analyses. We also searched a *Baetis rhodani* shotgun whole genome assembly [71]. The fragmentary nature of this assembly precluded confident annotation of opsin genes, but allowed us to find one candidate exon of a putative ortholog of *C. dipterum* UV-Ops1, that was too short to be included in the phylogenetic analyses (Supplementary Figure 5, Supplementary Table 15). To this dataset, we finally added the sequences for an additional 27 species covering seven orders were retrieved from [72].

All mayfly LWS, UV and Blue opsin sequences were carefully inspected and curated to detect gene annotation errors such fusions and fissions of gene models, missing or spurious exons and identical sequences derived from haplotypes of the same gene. These errors were corrected using available transcriptomic data and taking into account sequence and intron position conservation with opsin genes from other species and other opsin paralogs within the same species (Supplementary Table 15). Cd-hit was used to remove the redundant sequences resulting 687 insect opsins sequences. The alignment was performed using MAFFT with default parameters and regions with more the 70% of gaps were removed using TRIMAL. Phylogenetic reconstruction was performed using Ultrafast bootstrap with 1000 replicates, aLRT Bootstrap and aBayes [73,74] suing IQ-Tree-1.6.7 [75] under LG+G4+F model. Furthermore, we produced reduced version of the dataset where specific duplication from dragonflies were excluded. This data encompassed 359 sequences and was subject to a Bayesian phylogenetic reconstruction using Phylobayes MPI under GTR-G4 model and iqtree-1.6.7 under LG+G4+F [76,77]. In all the phylogenetic analyses the trees were rooted using the melatonin receptor which represent opsin closest outgroup [28].

5.2 Uv-Ops *in situ* hybridizations

Specific primers were designed to generate DIG-labelled probes against *UV-Ops2* and *UV-Ops4* (Supplementary Table 14). After o.n. fixation of the heads in FA 4% at 4º C, dissected retinas were bleached using Formamide Solution [78] for 16 hours. The following steps were as described for *OBP219* and *OBP199* genes. Briefly, we post-fixated the retinas and we treated them with Proteinase K for 15 min. The hybridization was carried out at 60º C o.n. The next day, retinas were incubated with anti-digoxigenin-POD at 4º C o.n. Finally, retinas were incubated with 1:100 TSA in borate buffer. Leica SPE confocal microscope was used to acquire images that were processed with Fiji [67].

### 6. Phylostratigraphy

To classify genes by origin (phylostratigraphy) we used an expanded dataset of 28 species (*Acanthoscurria geniculata, Apis mellifera, Caenorhabditis elegans, Cloeon dipterum, Daphnia pulex, Drosophila melanogaster, Ephemera danica, Heliconius melpomene, Holacanthella duospinosa, Homo sapiens, Ixodes scapularis, Ladona fulva, Laodelphax striatella, Lepidurus arcticus, Limulus polyphemus, Lingula anatina, Locusta migratoria, Nematostella vectensis, Parhyale hawaiensis, Pediculus humanus, Penaeus vannamei, Ramazzottius varieornatus, Sinella curviseta, Stegodyphus mimosarum, Strigamia maritima, Tigriopus californicus, Tribolium castaneum, Zootermopsis nevadensis*) aiming to sample more densely arthropod lineages and key taxonomic clades. OrthoFinder2 [23] with DIAMOND [24] as search engine and an inflation parameter of 2 were used to compute orthogroups. A higher inflation parameter was required to increase granularity aiming at capturing more defined orthogroups. Then, the orthogroups with sequences from *Cloeon* and *Drosophila* were selected, and each gene for each species was assigned an age based on the furthest species that had a member of the same orthogroup (present in *Cloeon* and *Nematostella* -> phylostratum Planulozoa). This resulted in 13 phylostratums for *Drosophila* and 13 for *Cloeon* (see Supplementary Table 5). To compute enrichment tests on gene age, the proportion of gene ages/phylostratums in a subset of interest (e.g. genes in WGCNA module "wings") was compared to the background proportion of ages of the species of interest using a fisher exact test fisher.test (alternative = "two.sided") in R. The list of p-values for each phylostratum were corrected for multiple-testing using p.adjust(method = "BH") in R, and only those enrichments with a adjusted p-value < 0.01 were classified as significant.

### 7. Evolutionary origin of wings

#### 7.1 Functional test of shared genes

A set of genes with unknown function (Flybase) or GO term assigned was selected to test functionally in *Drosophila* wings. VDRC lines (see Supplementary Table 13) were crossed to *yw; nub-Gal4;* + line to expressed the RNAi constructs specifically in the wing. Crosses were kept at 25º C for 48 hours and then switched to 29º C. Wings were dissected from adult females and mounted in Hoyer's/Lactic medium to image capture.

#### 7.2 Tissue specific transcriptomics

To characterise similarities between tissue-specific transcriptomes, we calculated which genes where expressed in wing pads and one of the other tissues preferentially, according to cRPKM (Minimum fraction of the minimum expr of test group that has to separate both distributions). We considered that the minimum expression of the test group was 20 and that the difference between the test group (wing pad and second tissue) with the rest of the tissues was at least of 30 % (Supplementary Figure 6d, e). This analysis resulted in 98 genes with wing pad-specific expression and from those, in 42 of the cases, the second tissue was the gills.


1 Almudi, I. *et al.* Establishment of the mayfly Cloeon dipterum as a new model system to investigate insect evolution. *Evodevo* **10**, 6, doi:10.1186/s13227-019-0120-y (2019).

2 Marcais, G. & Kingsford, C. A fast, lock-free approach for efficient parallel counting of occurrences of k-mers. *Bioinformatics* **27**, 764-770, doi:10.1093/bioinformatics/btr011 (2011).

3 Liu, B. *et al.* Estimation of genomic characteristics by analyzing k-mer frequency in de novo genome projects. *arXiv:1308.2012* (2013).

4 Simpson, J. T. Exploring genome characteristics and sequence quality without a reference. *Bioinformatics* **30**, 1228-1235, doi:10.1093/bioinformatics/btu023 (2014).

5 Wood, D. & Salzberg, S. Kraken: ultrafast metagenomic sequence classification using exact alignments. *Genome Biology* **15**, R46 (2014).

6 Marco-Sola, S., Sammeth, M., Guigo, R. & Ribeca, P. The GEM mapper: fast, accurate and versatile alignment by filtration. *Nat Methods*, doi:10.1038/nmeth.2221 (2012).

7 Zimin, A. V. *et al.* Hybrid assembly of the large and highly repetitive genome of Aegilops tauschii, a progenitor of bread wheat, with the mega-reads algorithm. *bioRxiv*, doi:10.1101/066100 (2016).

8 Zimin, A. V. *et al.* The MaSuRCA genome assembler. *Bioinformatics* **29**, 2669-2677, doi:10.1093/bioinformatics/btt476 (2013).

9 Parra, G., Bradnam, K. & Korf, I. CEGMA: a pipeline to accurately annotate core genes in eukaryotic genomes. *Bioinformatics* **23**, 1061-1067, doi:10.1093/bioinformatics/btm071 (2007).

10 Simão, F. A., Waterhouse, R. M., Ioannidis, P., Kriventseva, E. V. & Zdobnov, E. M. BUSCO: assessing genome assembly and annotation completeness with single-copy orthologs. *Bioinformatics* **31**, doi:10.1093/bioinformatics/btv351 (2015).

11 Grabherr, M. G. *et al.* Full-length transcriptome assembly from RNA-Seq data without a reference genome. *Nat Biotechnol* **29**, 644-652, doi:10.1038/nbt.1883 (2011).

12 Hass, J., Blaschke, S., Rammsayer, T. & Herrmann, J. M. A neurocomputational model for optimal temporal processing. *J Comput Neurosci* **25**, 449-464, doi:10.1007/s10827-008-0088-4 (2008).

13 Stanke, M., Tzvetkova, A. & Morgenstern, B. AUGUSTUS at EGASP: using EST, protein and genomic alignments for improved gene prediction in the human genome. *Genome Biol* **7 Suppl 1**, S11 11-18, doi:10.1186/gb-2006-7-s1-s11 (2006).

14 Dobin, A. *et al.* STAR: ultrafast universal RNA-seq aligner. *Bioinformatics* **29**, 15-21, doi:10.1093/bioinformatics/bts635 (2013).

15 Pertea, M. *et al.* StringTie enables improved reconstruction of a transcriptome from RNA-seq reads. *Nat Biotechnol* **33**, 290-295, doi:10.1038/nbt.3122 (2015).

16 Niknafs, Y. S., Pandian, B., Iyer, H. K., Chinnaiyan, A. M. & Iyer, M. K. TACO produces robust multisample transcriptome assemblies from RNA-seq. *Nat Methods* **14**, 68-70, doi:10.1038/nmeth.4078 (2017).

17 Smit, A. & Hubley, R. RepeatModeler Open-1.0. . http://www.repeatmasker.org/ (2008).

18 Bao, W., Kojima, K. K. & Kohany, O. Repbase Update, a database of repetitive elements in eukaryotic genomes. *Mob DNA* **6**, 11, doi:10.1186/s13100-015-0041-9 (2015).

19 Smit, A., Hubley, R. & Green, P. RepeatMasker Open-4.0. http://www.repeatmasker.org/ (2013).

20 Blanchette, M. *et al.* Aligning multiple genomic sequences with the threaded blockset aligner. *Genome Res* **14**, 708-715, doi:10.1101/gr.1933104 (2004).

21 Roth, A. C., Gonnet, G. H. & Dessimoz, C. Algorithm of OMA for large-scale orthology inference. *BMC Bioinformatics* **9**, 518, doi:10.1186/1471-2105-9-518 (2008).

22 Margulies, E. H., Blanchette, M., Program, N. C. S., Haussler, D. & Green, E. D. Identification and characterization of multi-species conserved sequences. *Genome Res* **13**, 2507-2518, doi:10.1101/gr.1602203 (2003).

23 Emms, D. M. & Kelly, S. OrthoFinder: phylogenetic orthology inference for comparative genomics. *bioRxiv*, 466201, doi:10.1101/466201 (2019).

24 Buchfink, B., Xie, C. & Huson, D. H. Fast and sensitive protein alignment using DIAMOND. *Nature Methods* **12**, 59, doi:10.1038/nmeth.3176

https://www.nature.com/articles/nmeth.3176 - supplementary-information (2014).

25 Katoh, K. & Standley, D. M. MAFFT multiple sequence alignment software version 7: improvements in performance and usability. *Molecular biology and evolution* **30**, 772-780, doi:10.1093/molbev/mst010 (2013).

26 Capella-Gutiérrez, S., Silla-Martínez, J. M. & Gabaldón, T. trimAl: a tool for automated alignment trimming in large-scale phylogenetic analyses. *Bioinformatics* **25**, 1972-1973, doi:10.1093/bioinformatics/btp348 (2009).

27 Kozlov, A. M., Darriba, D., Flouri, T., Morel, B. & Stamatakis, A. RAxML-NG: a fast, scalable and user-friendly tool for maximum likelihood phylogenetic inference. *Bioinformatics*, doi:10.1093/bioinformatics/btz305 (2019).

28 Feuda, R., Hamilton, S. C., McInerney, J. O. & Pisani, D. Metazoan opsin evolution reveals a simple route to animal vision. *Proceedings of the National Academy of Sciences* **109**, 18868-18872, doi:10.1073/pnas.1204609109 (2012).

29 Enright, A. J., Van Dongen, S. & Ouzounis, C. A. An efficient algorithm for large-scale detection of protein families. *Nucleic acids research* **30**, 1575-1584, doi:10.1093/nar/30.7.1575 (2002).

30 Kumar, L. & E Futschik, M. Mfuzz: a software package for soft clustering of microarray data. *Bioinformation* **2**, 5-7, doi:10.6026/97320630002005 (2007).

31 Huang, D. W., Sherman, B. T. & Lempicki, R. A. Systematic and integrative analysis of large gene lists using DAVID bioinformatics resources. *Nature Protocols* **4**, 44, doi:10.1038/nprot.2008.211 https://www.nature.com/articles/nprot.2008.211 - supplementary-information (2008).

32 Kim, D., Paggi, J. M., Park, C., Bennett, C. & Salzberg, S. L. Graph-based genome alignment and genotyping with HISAT2 and HISAT-genotype. *Nature Biotechnology* **37**, 907-915, doi:10.1038/s41587-019-0201-4 (2019).

33 Anders, S., Pyl, P. T. & Huber, W. HTSeq—a Python framework to work with high-throughput sequencing data. *Bioinformatics* **31**, 166-169, doi:10.1093/bioinformatics/btu638 (2014).

34 Love, M. I., Huber, W. & Anders, S. Moderated estimation of fold change and dispersion for RNA-seq data with DESeq2. *Genome Biol* **15**, 550, doi:10.1186/s13059-014-0550-8 (2014).

35 Labbé, R. M. *et al.* A Comparative Transcriptomic Analysis Reveals Conserved Features of Stem Cell Pluripotency in Planarians and Mammals. *STEM CELLS* **30**, 1734-1745, doi:10.1002/stem.1144 (2012).

36 Torres-Méndez, A. *et al.* A novel protein domain in an ancestral splicing factor drove the evolution of neural microexons. *Nature Ecology & Evolution* **3**, 691-701, doi:10.1038/s41559-019-0813-6 (2019).

37 Langfelder, P. & Horvath, S. WGCNA: an R package for weighted correlation network analysis. *BMC Bioinformatics* **9**, 559, doi:10.1186/1471-2105-9-559 (2008).

38 Harrison, M. C. *et al.* Hemimetabolous genomes reveal molecular basis of termite eusociality. *Nature Ecology & Evolution* **2**, 557-566, doi:10.1038/s41559-017-0459-1 (2018).

39 Benton, R., Vannice, K. S., Gomez-Diaz, C. & Vosshall, L. B. Variant ionotropic glutamate receptors as chemosensory receptors in Drosophila. *Cell* **136**, 149-162, doi:10.1016/j.cell.2008.12.001 (2009).

40 Vogt, R. G. *et al.* The insect SNMP gene family. *Insect Biochem Mol Biol* **39**, 448-456, doi:10.1016/j.ibmb.2009.03.007 (2009).

41 Vieira, F. G. & Rozas, J. Comparative genomics of the odorant-binding and chemosensory protein gene families across the Arthropoda: origin and evolutionary history of the chemosensory system. *Genome Biol Evol* **3**, 476-490, doi:10.1093/gbe/evr033 (2011).

42 Ioannidis, P. *et al.* Genomic Features of the Damselfly Calopteryx splendens Representing a Sister Clade to Most Insect Orders. *Genome Biol Evol* **9**, 415-430, doi:10.1093/gbe/evx006 (2017).

43 Missbach, C. *et al.* Evolution of insect olfactory receptors. *Elife* **3**, e02115, doi:10.7554/eLife.02115 (2014).

44 Kirkness, E. F. *et al.* Genome sequences of the human body louse and its primary endosymbiont provide insights into the permanent parasitic lifestyle. *Proc Natl Acad Sci U S A* **107**, 12168-12173, doi:10.1073/pnas.1003379107 (2010).

45 Terrapon, N. *et al.* Molecular traces of alternative social organization in a termite genome. *Nat Commun* **5**, 3636, doi:10.1038/ncomms4636 (2014).

46 Wu, C. *et al.* Analysis of the genome of the New Zealand giant collembolan (Holacanthella duospinosa) sheds light on hexapod evolution. *BMC Genomics* **18**, 795-795, doi:10.1186/s12864-017-4197-1 (2017).

47 Brand, P. *et al.* The origin of the odorant receptor gene family in insects. *eLife* **7**, e38340, doi:10.7554/eLife.38340 (2018).

48 Frias-Lopez, C. *et al.* Comparative analysis of tissue-specific transcriptomes in the funnel-web spider Macrothele calpeiana (Araneae, Hexathelidae). *PeerJ* **3**, e1064, doi:10.7717/peerj.1064 (2015).

49 Vizueta, J., Sánchez-Gracia, A. & Rozas, J. BITACORA: A comprehensive tool for the identification and annotation of gene families in genome assemblies. *bioRxiv*, 593889, doi:10.1101/593889 (2019).

50 Vizueta, J., Rozas, J. & Sánchez-Gracia, A. Comparative Genomics Reveals Thousands of Novel Chemosensory Genes and Massive Changes in Chemoreceptor Repertories across Chelicerates. *Genome biology and evolution* **10**, 1221-1236, doi:10.1093/gbe/evy081 (2018).

51 Krogh, A., Larsson, B., von Heijne, G. & Sonnhammer, E. L. Predicting transmembrane protein topology with a hidden Markov model: application to complete genomes. *J Mol Biol* **305**, 567-580, doi:10.1006/jmbi.2000.4315 (2001).

52 Kall, L., Krogh, A. & Sonnhammer, E. L. A combined transmembrane topology and signal peptide prediction method. *J Mol Biol* **338**, 1027-1036, doi:10.1016/j.jmb.2004.03.016 (2004).

53 Croset, V. *et al.* Ancient protostome origin of chemosensory ionotropic glutamate receptors and the evolution of insect taste and olfaction. *PLoS Genet* **6**, e1001064, doi:10.1371/journal.pgen.1001064 (2010).

54 Petersen, T. N., Brunak, S., von Heijne, G. & Nielsen, H. SignalP 4.0: discriminating signal peptides from transmembrane regions. *Nat Methods* **8**, 785-786, doi:10.1038/nmeth.1701 (2011).

55 Katoh, K. & Standley, D. M. MAFFT multiple sequence alignment software version 7: improvements in performance and usability. *Molecular biology and evolution* **30**, 772-780, doi:10.1093/molbev/mst010 (2013).

56 Nguyen, L. T., Schmidt, H. A., von Haeseler, A. & Minh, B. Q. IQ-TREE: a fast and effective stochastic algorithm for estimating maximum-likelihood phylogenies. *Molecular biology and evolution* **32**, 268-274, doi:10.1093/molbev/msu300 (2015).

57 Hoang, D. T. *et al.* MPBoot: fast phylogenetic maximum parsimony tree inference and bootstrap approximation. *BMC evolutionary biology* **18**, 11-11, doi:10.1186/s12862-018-1131-3 (2018).

58 Letunic, I. & Bork, P. Interactive Tree Of Life (iTOL): an online tool for phylogenetic tree display and annotation. *Bioinformatics* **23**, 127-128, doi:10.1093/bioinformatics/btl529 (2007).

59 Li, L., Stoeckert, C. J., Jr. & Roos, D. S. OrthoMCL: identification of ortholog groups for eukaryotic genomes. *Genome Res* **13**, 2178-2189, doi:10.1101/gr.1224503 (2003).

60 Robertson, H. M., Baits, R. L., Walden, K. K. O., Wada-Katsumata, A. & Schal, C. Enormous expansion of the chemosensory gene repertoire in the omnivorous German cockroach Blattella germanica. *Journal of Experimental Zoology Part B: Molecular and Developmental Evolution* **330**, 265-278, doi:10.1002/jez.b.22797 (2018).

61 Librado, P. & Rozas, J. Uncovering the functional constraints underlying the genomic organization of the odorant-binding protein genes. *Genome Biol Evol* **5**, 2096-2108, doi:10.1093/gbe/evt158 (2013).

62 Mei, T., Fu, W.-B., Li, B., He, Z.-B. & Chen, B. Comparative genomics of chemosensory protein genes (CSPs) in twenty-two mosquito species (Diptera: Culicidae): Identification, characterization, and evolution. *PloS one* **13**, e0190412-e0190412, doi:10.1371/journal.pone.0190412 (2018).

63 Sanchez-Gracia, A., Vieira, F. G. & Rozas, J. Molecular evolution of the major chemosensory gene families in insects. *Heredity (Edinb)* **103**, 208-216, doi:10.1038/hdy.2009.55 (2009).

64 Penalva-Arana, D. C., Lynch, M. & Robertson, H. M. The chemoreceptor genes of the waterflea Daphnia pulex: many Grs but no Ors. *BMC Evol Biol* **9**, 79, doi:10.1186/1471-2148-9-79 (2009).

65 Knecht, Z. A. *et al.* Distinct combinations of variant ionotropic glutamate receptors mediate thermosensation and hygrosensation in Drosophila. *eLife* **5**, e17879, doi:10.7554/eLife.17879 (2016).

66 Gomez-Diaz, C. *et al.* A CD36 ectodomain mediates insect pheromone detection via a putative tunnelling mechanism. *Nat Commun* **7**, 11866, doi:10.1038/ncomms11866 (2016).

67 Schindelin, J. *et al.* Fiji: an open-source platform for biological-image analysis. *Nature Methods* **9**, 676-682, doi:10.1038/nmeth.2019 (2012).

68 Futahashi, R. *et al.* Extraordinary diversity of visual opsin genes in dragonflies. *Proceedings of the National Academy of Sciences* **112**, E1247-E1256, doi:10.1073/pnas.1424670112 (2015).

69 Suvorov, A. *et al.* Opsins have evolved under the permanent heterozygote model: insights from phylotranscriptomics of Odonata. *Mol Ecol* **26**, 1306-1322, doi:10.1111/mec.13884 (2017).

70 Misof, B. *et al.* Phylogenomics resolves the timing and pattern of insect evolution. *Science* **346**, 763-767, doi:10.1126/science.1257570 (2014).

71 Macdonald, H. C., Ormerod, S. J. & Bruford, M. W. Enhancing capacity for freshwater conservation at the genetic level: a demonstration using three stream macroinvertebrates. *Aquatic Conservation: Marine and Freshwater Ecosystems* **27**, 452-461, doi:10.1002/aqc.2691 (2017).

72 Feuda, R., Marletaz, F., Bentley, M. A. & Holland, P. W. Conservation, Duplication, and Divergence of Five Opsin Genes in Insect Evolution. *Genome Biol Evol* **8**, 579-587, doi:10.1093/gbe/evw015 (2016).

73 Hoang, D. T., Chernomor, O., von Haeseler, A., Minh, B. Q. & Vinh, L. S. UFBoot2: Improving the Ultrafast Bootstrap Approximation. *Molecular biology and evolution* **35**, 518-522, doi:10.1093/molbev/msx281 (2017).

74 Anisimova, M., Gil, M., Dufayard, J.-F., Dessimoz, C. & Gascuel, O. Survey of branch support methods demonstrates accuracy, power, and robustness of fast likelihood-based approximation schemes. *Systematic biology* **60**, 685-699, doi:10.1093/sysbio/syr041 (2011).

75 Nguyen, L.-T., Schmidt, H. A., von Haeseler, A. & Minh, B. Q. IQ-TREE: A Fast and Effective Stochastic Algorithm for Estimating Maximum-Likelihood Phylogenies. *Molecular biology and evolution* **32**, 268-274, doi:10.1093/molbev/msu300 (2014).

76 Le, S. Q. & Gascuel, O. An Improved General Amino Acid Replacement Matrix. *Molecular biology and evolution* **25**, 1307-1320, doi:10.1093/molbev/msn067 (2008).

77 Lartillot, N., Rodrigue, N., Stubbs, D. & Richer, J. PhyloBayes MPI: Phylogenetic Reconstruction with Infinite Mixtures of Profiles in a Parallel Environment. *Systematic Biology* **62**, 611-615, doi:10.1093/sysbio/syt022 (2013).

78 King, R. S. & Newmark, P. A. In situ hybridization protocol for enhanced detection of gene expression in the planarian Schmidtea mediterranea. *BMC developmental biology* **13**, 8-8, doi:10.1186/1471-213X-13-8 (2013).
